# Localizing components of shared transethnic genetic architecture of complex traits from GWAS summary data

**DOI:** 10.1101/858431

**Authors:** Huwenbo Shi, Kathryn S. Burch, Ruth Johnson, Malika K. Freund, Gleb Kichaev, Nicholas Mancuso, Astrid M. Manuel, Natalie Dong, Bogdan Pasaniuc

## Abstract

Despite strong transethnic genetic correlations reported in the literature for many complex traits, the non-transferability of polygenic risk scores across populations suggests the presence of population-specific components of genetic architecture. We propose an approach that models GWAS summary data for one trait in two populations to estimate genome-wide proportions of population-specific/shared causal SNPs. In simulations across various genetic architectures, we show that our approach yields approximately unbiased estimates with in-sample LD and slight upward-bias with out-of-sample LD. We analyze 9 complex traits in individuals of East Asian and European ancestry, restricting to common SNPs (MAF > 5%), and find that most common causal SNPs are shared by both populations. Using the genome-wide estimates as priors in an empirical Bayes framework, we perform fine-mapping and observe that high-posterior SNPs (for both the population-specific and shared causal configurations) have highly correlated effects in East Asians and Europeans. In population-specific GWAS risk regions, we observe a 2.8x enrichment of shared high-posterior SNPs, suggesting that population-specific GWAS risk regions harbor shared causal SNPs that are undetected in the other GWAS due to differences in LD, allele frequencies, and/or sample size. Finally, we report enrichments of shared high-posterior SNPs in 53 tissue-specific functional categories and find evidence that SNP-heritability enrichments are driven largely by many low-effect common SNPs.

## Introduction

Genetic and phenotypic variations among humans have been shaped by many factors, including migration histories, geodemographic events, and environmental background^1–5^. As a result, the underlying genetic architecture of a given complex trait – defined here in terms of ‘polygenicity’ (the number of variants with nonzero effects)^6–10^ and the coupling of causal effect sizes with minor allele frequency (MAF)^11,12^, linkage disequilibrium (LD)^13–15^, and other genomic features^16^ – varies among ancestral populations. While the vast majority of genome-wide association studies (GWAS) to date have been performed in individuals of European descent^17–20^, growing numbers of studies performed in individuals of non-European ancestry^21–27^ have created opportunities for well-powered transethnic genetic studies^21,22,24,26,28–33^.

Risk regions identified through GWAS tend to replicate across populations^17,21,22,33–35^, indicating that complex traits have shared genetic components among populations. Indeed, for certain post-GWAS analyses such as disease mapping^23,31,36^ and statistical fine-mapping^28,37–40^, under the assumption that two populations share one or more causal variants, population-specific LD patterns can be leveraged to improve performance over approaches that model a single population. On the other hand, several studies have shown that heterogeneity in genetic architectures limits transferability of polygenic risk scores (PRS) across populations^5,41–48^; critically, if applied in a clinical setting, existing PRS may exacerbate health disparities among ethnic groups^49^. The population-specificity of existing PRS as well as estimates of transethnic genetic correlations less than one reported in the literature^30,50–53^ indicate that (1) LD tagging and allele frequencies of shared causal variants vary across populations, (2) that a sizeable number of causal variants are population-specific, and/or (3) that causal effect sizes vary across populations due to, for example, different gene-environment interactions. For example, due to population-specific LD, a single genetic variant that is significantly associated with a trait in two populations may actually be tagging distinct population-specific causal variants (Figure 1). Conversely, two distinct associations in two populations may be driven by the same underlying causal variants (i.e. colocalization). Thus, identifying shared and population-specific components of genetic architecture could help improve transethnic analyses (e.g., transferability of PRS across populations^19,41,42,45,46^) and uncover novel disease etiologies.

**Figure 1:**
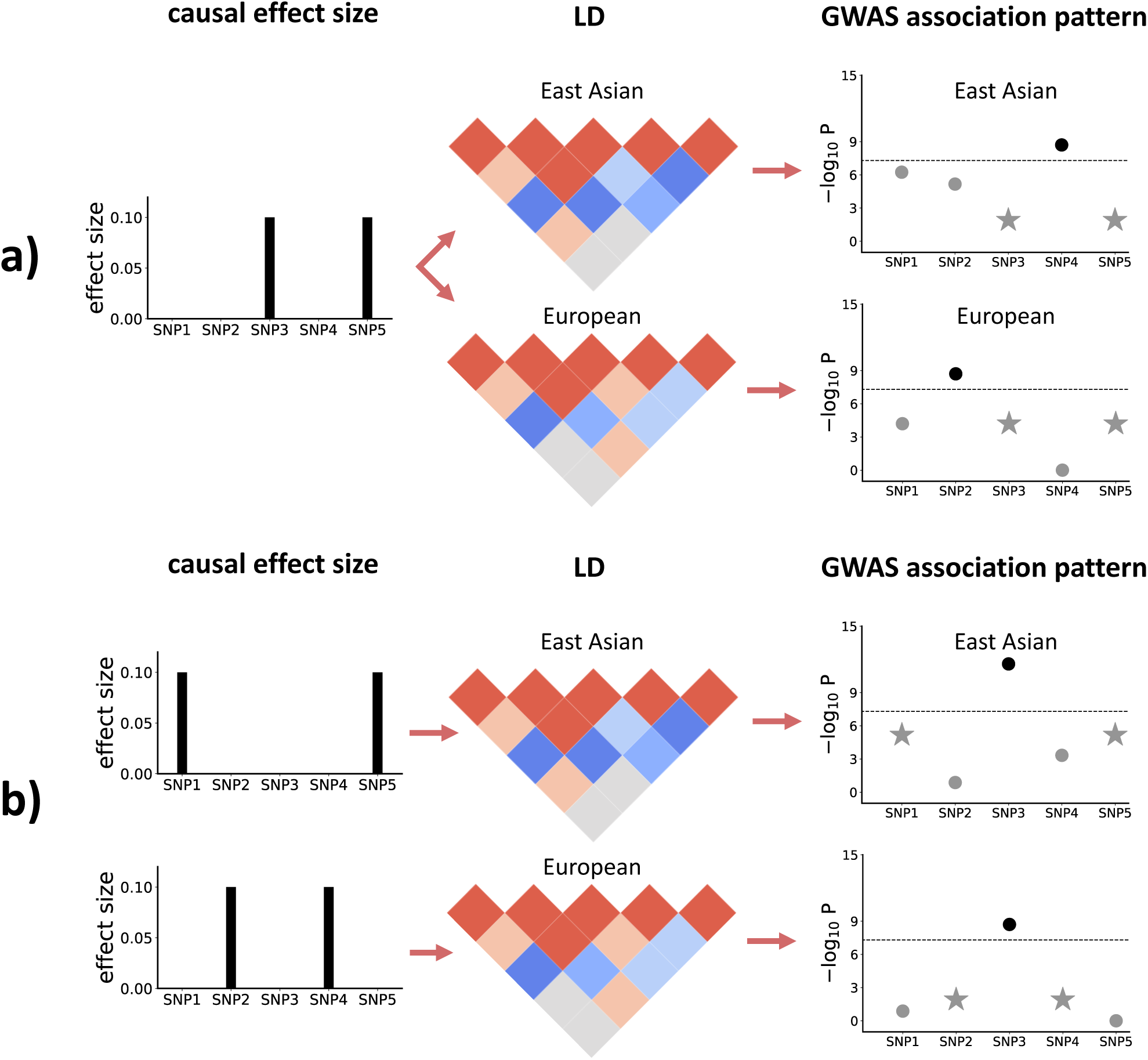
Toy examples to illustrate how population-specific LD patterns affect GWAS associations. a) SNPs 3 and 5 are causal in both East Asians and Europeans and have the same population-specific causal effect size of 0.1. However, due to different LD patterns in East Asians and Europeans, SNPs 2 and 4 are observed to be GWAS-significant, respectively. b) Different SNPs are causal in East Asians (SNPs 1 and 5) and Europeans (SNPs 2 and 4). However, due to population-specific LD, SNP 3 is observed to be GWAS-significant in both populations. The stars in the rightmost plots represent the SNPs with true nonzero effects; the GWAS-significant SNP is highlighted in a darker color.

In this work, we introduce PESCA (Population-spEcific/Shared Causal vAriants), an approach that requires only GWAS summary association statistics and ancestry-matched estimates of LD to infer genome-wide proportions of population-specific and shared causal variants for a single trait in two populations. These estimates are then used as priors in an empirical Bayes framework to localize and test for enrichment of population-specific/shared causal variants in regions of interest. In this context, a “causal variant” is a variant measured in the given GWAS that either has a nonzero effect on the trait (e.g., a nonsynonymous variant that alters protein folding) or tags a nonzero effect at an unmeasured variant through LD. It is therefore important to note that the set of “causal variants” that PESCA aims to identify is defined with respect to the set of variants included in the GWAS and can contain variants with indirect nonzero effects that are statistical rather than biological in nature (this is analogous to the definition of SNP-heritability, which is also a function of a specific set of SNPs^11,54–56^). Through extensive simulations, we show that our method yields approximately unbiased estimates of the proportions of population-specific/shared causal variants if in-sample LD is used and slightly upward-biased estimates if LD is estimated from an external reference panel. We then show that using these estimates as priors to perform fine-mapping (Methods) produces well-calibrated per-SNP posterior probabilities and enrichment test statistics. We note that the definition of enrichment used here is related to, but conceptually distinct from, definitions of SNP-heritability enrichment^13,16^. Under our framework, an enrichment of causal SNPs greater than 1 indicates that, compared to the genome-wide background, there are more causal SNPs in that region than expected^57,58^ (Methods). In contrast, an enrichment of SNP-heritability greater than 1 indicates that the average per-SNP effect size in the region is larger than the genome-wide average per-SNP effect size.

We apply our approach to publicly available GWAS summary statistics for 9 complex traits and diseases in individuals of East Asian (EAS) and European (EUR) ancestry (average N_EAS_ = 94,621, N_EUR_ = 103,507) (Table 1), restricting to common SNPs (MAF > 5%) and using 1000 Genomes^59^ to estimate ancestry-matched LD. On average across the 9 traits, we estimate that approximately 80% (S.D. 15%) of common SNPs that are causal in EAS and 84% (S.D. 8%) of those in EUR are shared by the other population. Consistent with previous studies based on SNP-heritability^55,60^, we find that high-posterior SNPs are distributed uniformly across the genome. We observe that population-specific GWAS risk regions have, on average across the 9 traits, a 2.8x enrichment of shared high-posterior SNPs relative to the genome-wide background, suggesting that many EAS-specific and EUR-specific GWAS risk regions harbor shared causal SNPs that are undetected in the other population due to differences in LD, allele frequencies, and/or GWAS sample size. The effects of SNPs with posterior probability > 0.8 of being causal (for any causal configuration) are highly correlated between EAS and EUR, concordant with replication slopes between EAS and EUR marginal effects close to 1 that have been reported for several complex diseases^33^ and with strong transethnic genetic correlations previously reported for the same traits analyzed in this work (average 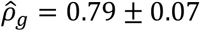 s.e.m. across the 9 traits)^51^. Finally, we show that regions flanking genes that are specifically expressed in trait-relevant tissues^61^ harbor a disproportionate number of shared high-posterior SNPs – many of the same tissue-specific gene sets are also enriched with SNP-heritability, implying that SNP-heritability enrichments are driven by many low-effect SNPs rather than a small number of high-effect SNPs. Our results suggest that common causal SNPs have similar etiological roles in EAS and EUR and that transferability of PRS and other GWAS findings across populations can be improved by explicitly correcting for population-specific LD and allele frequencies.

**Table 1:**
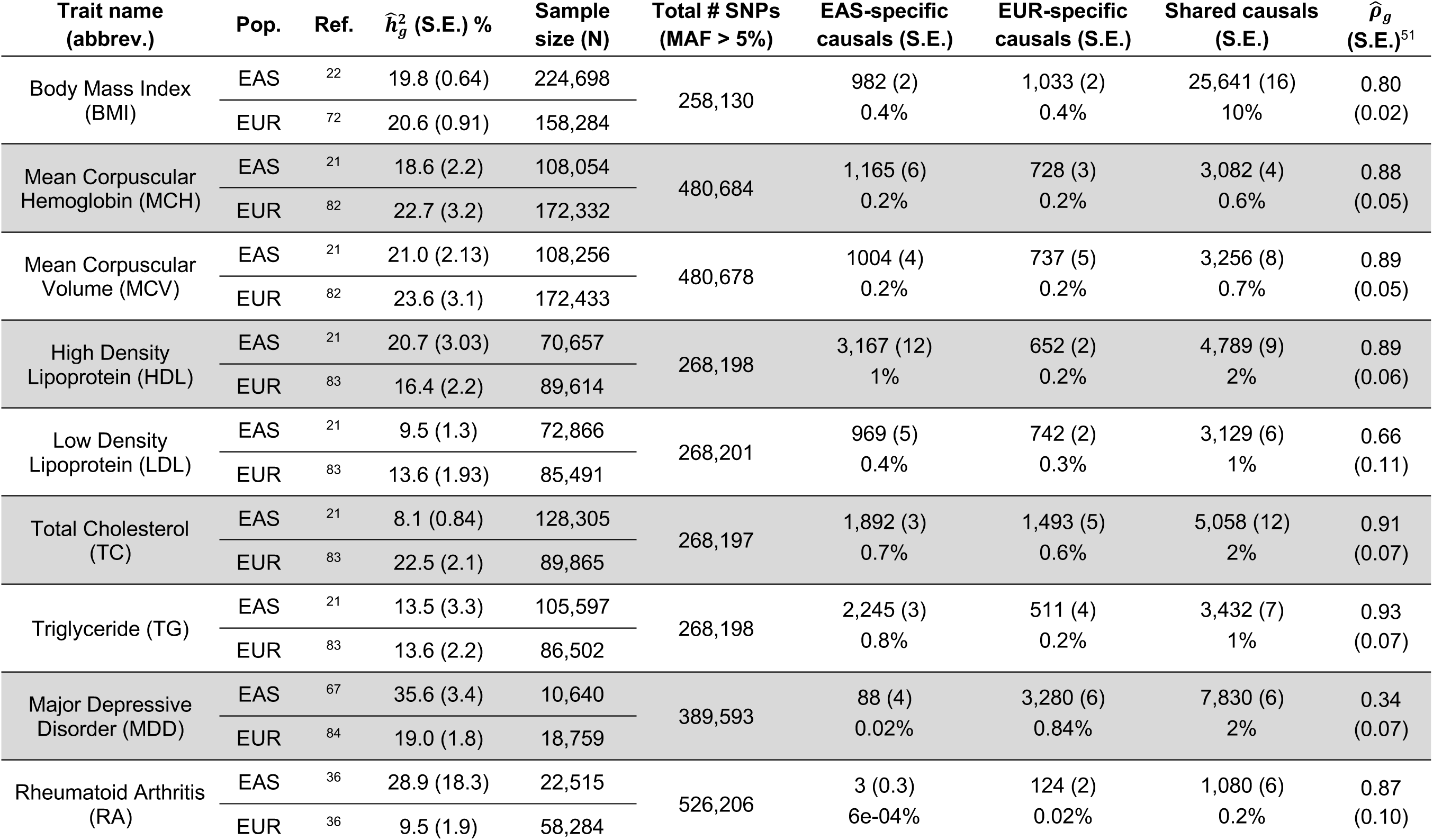
Estimated numbers and percentages of population-specific/shared common causal SNPs for 9 complex traits. We estimated genome-wide SNP-heritability using LD score regression^54^ with the intercept constrained to 1 (i.e. assuming no population stratification). Trans-ethnic genetic correlation estimates 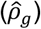 computed from a similar set of summary statistics were obtained from a previous study^51^. Standard errors of the estimated numbers of population-specific/shared causal SNPs were computed using the last 50 iterations of the EM-MCMC algorithm.

## Material and Methods

### Distribution of GWAS summary statistics in two populations

For a given complex trait, we model the causal statuses of SNP *i* in two populations as a binary vector of size two, ***C***_*i*_ *= c*_*i*1_*c*_*i*2_, where each bit, *c*_*i*1_ ∈ {0,1} and *c*_*i*2_ ∈ {0,1}, represents the causal status of SNP *i* in populations 1 and 2, respectively. ***C***_*i*_ *=* 00 indicates that SNP *i* is not causal in either population; ***C***_*i*_ *=* 01 and ***C***_*i*_ *=* 10 indicate that SNP *i* is causal only in the first and second population, respectively; and ***C***_*i*_ *=* 11 indicates that SNP *i* is causal in both populations. We assume ***C***_*i*_ follows a multivariate Bernoulli (MVB) distribution^62,63^

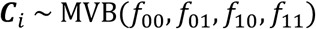

in order to facilitate optimization and interpretation (Supplemental Note). Assuming the causal status vector of a SNP is independent from those of other SNPs (***C***_*i*_ ⊥ ***C***_*j*_ for *i* ≠ *j*), the joint probability of the causal statuses of *p* SNPs is 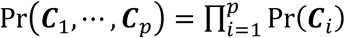.

Given two genome-wide association studies with sample sizes *n*_1_ and *n*_2_ for the first and second populations, respectively, we derive the distribution of Z-scores, ***Z***_1_ and ***Z***_2_ (both are *p* × 1 vectors), conditional on the causal status vectors for each population, ***c***_1_ *=* (*c*_11_, … *c*_*p*1_)^*T*^ and ***c***_2_ *=* (*c*_12_ … *c*_*p*2_)^*T*^. Although it is reasonable to suspect that there are nonzero cross-population correlations of effect sizes at shared causal SNPs, to facilitate inference, we impose the (potentially strong) assumption that ***Z***_1_ and ***Z***_2_ are independent given ***c***_1_ and ***c***_2_. Thus, for population *j*,

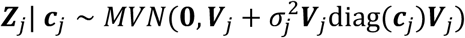

where ***V***_*j*_ is the *p* × *p* LD matrix for population *j*; diag(***c***_*j*_) is a diagonal matrix in which the *k*-th diagonal element is 1 if *c*_*kj*_ *=* 1 and 0 if *c*_*kj*_ *=* 0; and 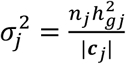, where 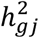 and |*c*_*j*_| are the SNP-heritability of the trait and the number of causal SNPs, respectively, in population *j* (Supplemental Note).

Finally, we derive the joint probability of ***Z***_1_ and ***Z***_2_ by integrating over all possible causal status vectors in the two populations:

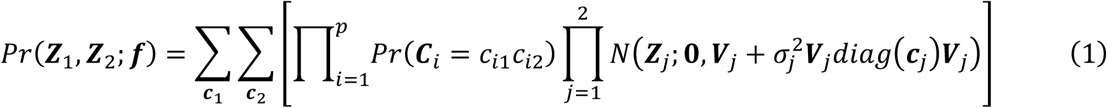

where ***f*** *=* (*f*_00_, *f*_01_, *f*_10_, *f*_11_) is the vector of parameters of the MVB distribution. In practice, we partition the genome into approximately independent regions^64^ and model the distribution of Z-scores at all regions as the product of the distribution of Z-scores in each region (Supplemental Note).

### Estimating genome-wide proportions of population-specific/shared causal SNPs

We use Expectation-Maximization (EM) coupled with Markov Chain Monte Carlo (MCMC) to maximize the likelihood function in Equation (1) over the MVB parameters ***f***. We initialize ***f*** to ***f*** *=* (0, −3.9, −3.9, 3.9) which corresponds to 2% of SNPs being causal in population 1, 2% being causal in population 2, and 2% being shared causals. In the expectation step, we approximate the surrogate function *Q*(***f***|***f***^(***t***)^) using an efficient Gibbs sampler; in the maximization step, we maximize *Q*(***f***|***f***^(***t***)^) using analytical formulae (Supplemental Note). From the estimated ***f***, denoted ***f***\*, we recover the proportions of population-specific and shared causal SNPs. For computational efficiency, we apply the EM algorithm to each chromosome in parallel and aggregate the chromosomal estimates to obtain estimates of the genome-wide proportions of population-specific/shared causal SNPs (Supplemental Note).

### Evaluating per-SNP posterior probabilities of being causal in a single or both populations

We estimate the posterior probability of each SNP to be causal in a single population (population-specific) or both populations (shared), using the estimated genome-wide proportions of population-specific and shared causal variants (obtained from ***f***\*) as prior probabilities in an empirical Bayes framework. Specifically, for each SNP *i*, we evaluate the posterior probabilities Pr(***C***_*i*_ *=* 01|***Z***_1_, ***Z***_2_; ***f***\*), Pr(***C***_*i*_ *=* 10|***Z***_1_, ***Z***_2_; ***f***\*), and Pr(***C***_*i*_ *=* 11|***Z***_1_, ***Z***_2_; ***f***\*). Since evaluating these probabilities requires integrating over the posterior probabilities of all 2^(2*p*)^ possible causal status configurations, we use a Gibbs sampler to efficiently approximate the posterior probabilities (Supplemental Note).

### Estimating the numbers of population-specific/shared causal SNPs in a region

We infer the posterior expected numbers of population-specific/shared causal SNPs in a region (e.g., an LD block or a chromosome) conditional on the Z-scores (***Z***_1_ and ***Z***_2_) by summing, across all SNPs in the region, the per-SNP posterior probabilities of being causal in a single or both populations. For example, in a region with *p* SNPs, the posterior expected number of shared causal SNPs is 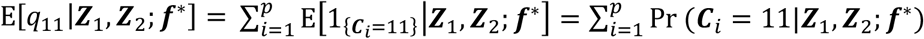. Since SNPs in a region are highly correlated, invalidating the use of jackknife to estimate standard errors, we refrain from reporting standard errors of the posterior expected regional numbers of population-specific/shared causal SNPs.

### Defining LD blocks that are approximately independent in two populations

For computational efficiency, PESCA assumes that, in both populations, a SNP in a given block is independent from all SNPS in all other blocks. This assumption requires defining blocks of SNPs that are approximately LD-independent in both populations. To this end, we first compute the “transethnic LD matrix” (***V***_*trans*_) from the East Asian- and European-ancestry LD matrices (***V***_*EAS*_ and ***V***_*EUR*_) by setting each element in the transethnic LD matrix to the larger of the East Asian-specific and European-specific pairwise LD; i.e. ***V***_*trans,ij*_ *=* ***V***_*EAS,ij*_ if |***V***_*EAS,ij*_| > |***V***_*EUR,ij*_| and ***V***_*trans,ij*_ *=* ***V***_E*UR,ij*_ if |***V***_*EUR,ij*_| > |***V***_*EAS,ij*_|. The resulting matrix ***V***_*trans*_ is block diagonal due to shared recombination hotspots in both populations; in practice, we apply this procedure to each chromosome separately to obtain 22 chromosome-wide transethnic LD matrices. We then apply LDetect^64^ to define LD blocks within the transethnic LD matrix. Applying this procedure using the 1000 Genomes Phase 3 reference panel^59^ to create the transethnic LD matrix produces 1,368 LD blocks (average length of 2-Mb) that are approximately independent in individuals of East Asian and European ancestry.

### Enrichment of population-specific/shared causal SNPs in functional annotations

We define the enrichment of population-specific/shared causal SNPs in a functional annotation as the ratio between the posterior and prior expected numbers of population-specific/shared causal SNPs. Specifically, we estimate the enrichment of population-specific/shared causal SNPs in a functional annotation *k* relative to the genome-wide background as

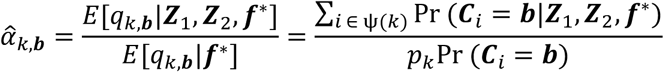

where ***b*** ∈ {01, 10, 11}, *q*_*k*,***b***_ is the number of population-specific (***b*** *=* 01 or ***b*** *=* 10) or shared (***b*** *=* 11) causal variants, ψ(*k*) is the set of SNPs in functional annotation *k*, and *p*_*k*_ is the number of SNPs in functional annotation *k*. The numerator, E[*q*_*k*,***b***_|***Z***_1_, ***Z***_2_, ***f***\*], and denominator, E[*q*_*k*,***b***_|***f***\*], represent the posterior (conditioned on Z-scores) and prior expected numbers of causal SNPs in functional annotation *k*, respectively. We estimate the standard error of 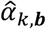 using block jackknife over 1,368 non-overlapping approximately LD-independent blocks across the entire genome. The resulting enrichment test statistics, 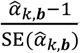, approximately follow a t-distribution with degrees of freedom equal to the number of blocks minus one^65^. Since we are interested in identifying categories of SNPs that harbor more population-specific/shared causal SNPs than expected (i.e. enrichment > 1), we report *P*-values from a one-tailed t-test where the null hypothesis is enrichment ≤ 1.

We note that our definition of enrichment of causal SNPs is related to, but conceptually different from, enrichment of SNP-heritability^13,16,66^. A positive enrichment of causal SNPs in a functional category indicates that, compared to the genome-wide background, there are more causal SNPs in that category than expected; a positive enrichment of SNP-heritability in a category indicates that the average per-SNP effect size in the category is larger than the genome-wide average per-SNP effect size.

### Simulation framework

We used real chromosome 22 genotypes of 10,000 individuals of East Asian ancestry from CONVERGE^67,68^ and 50,000 individuals of white British ancestry from the UK Biobank^69,70^ to simulate causal effects and phenotypes. First, we used PLINK^71^ (v1.9) to remove redundant SNPs in the 1000 Genomes Phase 3 reference panel^59^ such that there is no pair of SNPs with 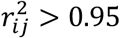 (*i* ≠ *j*) in the reference panel. We also removed strand-ambiguous SNPs and SNPs with MAF < 1% in either reference panel, resulting in a total of *M*=8,599 SNPs on chromosome 22 to use in simulations.

Given genotypes at *M* SNPs for *n*_*1*_ and *n*_*2*_ individuals in populations 1 and 2, respectively, we assume the standard linear models ***y***_**1**_ ***= X***_**1**_*β*_**1**_ + *ϵ*_**1**_ (population 1) and ***y***_**2**_ *=* ***X***_2_*β*_**2**_ + *ϵ*_**1**_ (population 2). We assume the phenotypes are standardized within each population such that E[***y***_**1**_] *=* **0**, Var[***y***_**1**_] *=* **I** and E[***y***_**2**_] *=* **0**, Var[***y***_**2**_] *=* **I**. Given ***c***_1_ and ***c***_2_, the index sets of causal SNPs in each population, the effects at the *i-*th causal SNP in each population, *β*_1*i*_ and *β*_2*i*_, are drawn from

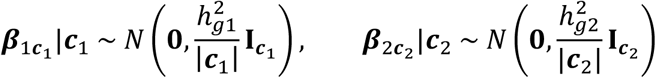

where 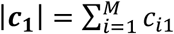 and 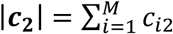 are the total numbers of causal SNPs in each population, 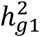 and 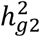 are the total SNP-heritabilities in each population, and E[*β*_1*i*_*β*_1*j*_] *=* Cov[*β*_1*i*_, *β*_1*j*_] *=* 0 and E[*β*_2*i*_ *β*_2*j*_] *=* Cov[*β*_2*i*_, *β*_2*j*_] *=* 0 for SNPs *i* ≠ *j*. The effects at non-causal SNPs are set to 0. The environmental effects for the *n*-th individual in each population are drawn i.i.d. from 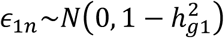 and 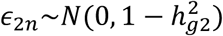.

Finally, given the real genotypes and simulated phenotypes for each population, we compute Z-scores for all SNPs in population *k* as 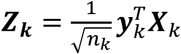.

### Application to 9 complex traits and diseases

We downloaded publicly available East Asian- and European-ancestry GWAS summary statistics for body mass index (BMI), mean corpuscular hemoglobin (MCH), mean corpuscular volume (MCV), high-density lipoprotein (HDL), low-density lipoprotein (LDL), total cholesterol (TC), triglycerides (TG), major depressive disorder (MDD), and rheumatoid arthritis (RA) from various sources (Table 1). The European-ancestry BMI GWAS is doubly corrected for genomic inflation factor^72^, which induces downward-bias in the estimated SNP-heritability; we correct this bias by re-inflating the Z-scores for this GWAS by a factor of 1.24. For all traits, we restrict to SNPs with MAF > 5% in both populations to reduce noise in the LD matrices estimated from 1000 Genomes^73^. We use PLINK^71^ (v.19) to remove redundant SNPs such that 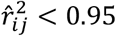 for all SNPs *i* ≠ *j* in both ancestry-matched 1000 Genomes^73^ reference panels. The resulting numbers of SNPs that were analyzed for each trait are listed in Table 1.

For each trait, we test for enrichment of population-specific/shared causal SNPs in 53 publicly available tissue-specific gene annotations^66^, each of which represents a set of genes that are “specifically expressed” in a GTEx^74^ tissue (referred to as “SEG annotations”). We set the threshold for statistical significance to *P*-value < 0.05/53 (Bonferroni correction for the number of tests performed per trait).

## Results

### Performance of PESCA in simulations

We assessed the performance of PESCA in simulations starting from real genotypes of individuals with East Asian^67,68^ (EAS) or European^69,70^ (EUR) ancestry (N_EAS_ = 10K, N_EUR_ = 50K, *M* = 8,599 SNPs) (Methods). First, we find that when in-sample LD from the GWAS is available, PESCA yields approximately unbiased estimates of the numbers of population-specific/shared causal SNPs (Figure 2, top panel). For example, in simulations where we randomly selected 50 EAS-specific, 50 EUR-specific, and 50 shared causal SNPs, we obtained estimates (and corresponding standard errors) of 37.8 (4.5) EAS-specific, 40.3 (4.9) EUR-specific, and 64.9 (6.3) shared causal SNPs, respectively. When external reference LD is used (in this case, from 1000 Genomes^73^), PESCA yields a slight upward bias (Figure 2, bottom panel); on the same simulated data, we obtained estimates of 48.0 (5.9) EAS-specific, 53.7 (7.44) EUR-specific, and 78.8 (7.6) shared causal SNPs.

**Figure 2:**
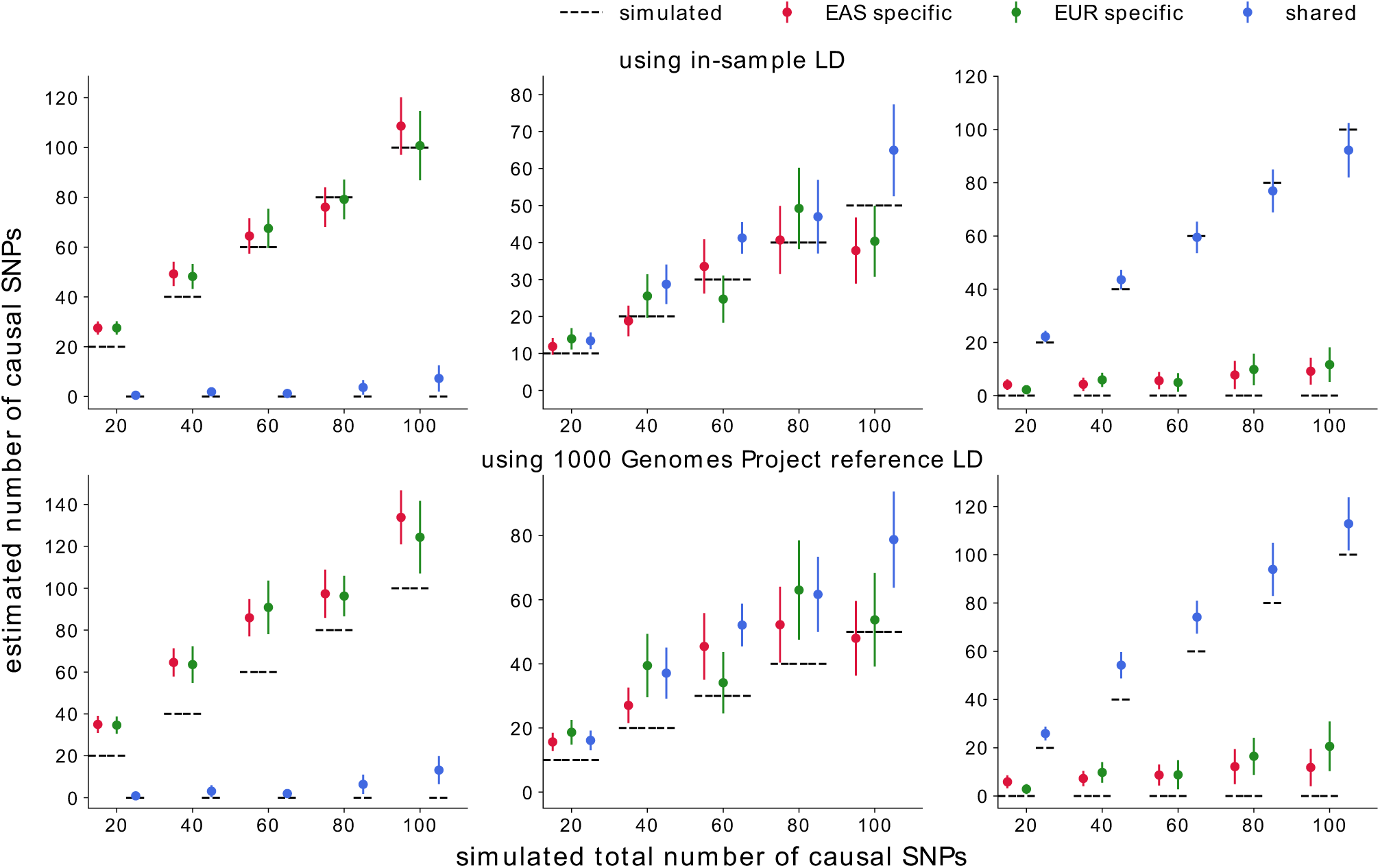
Genome-wide estimates of the numbers of population-specific/shared causal SNPs in simulations. The estimates are approximately unbiased when in-sample LD is used (top panel) and upward-biased estimates when external reference LD is used (bottom panel). For both populations, we simulate such that the product of SNP-heritability and GWAS sample size is 500. Mean and standard errors were obtained from 25 independent simulations. Error bars represent ±1.96 of the standard error.

We observe a slight decrease in accuracy as the effective sample size, the product of SNP-heritability and sample size 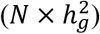, decreases (Figures S1-S5). This is expected as the likelihood of the GWAS summary statistics is a function of 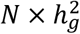 (Methods) – as the expected per-SNP variance at causal SNPs (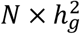 divided by the number of causal SNPs) decreases, GWAS summary statistics provide less information on the causal status of each SNP. Since it is often the case that the sample size of one GWAS is larger than that of the other, we perform simulations in which SNP-heritability is fixed to 0.05 in both populations, the EAS sample size is fixed to *N*_*EAS*_ *=* 10^4^, and the EUR sample size is varied such that the effective sample size of the EUR GWAS is 1-5x larger than that of the EAS GWAS. We find that the genome-wide estimators are relatively robust with in-sample LD; with external estimates of LD, when effective sample size differs by a factor of 2 or more, the estimator for the number of EUR-specific causal SNPs becomes less biased while the EAS-specific and shared causal estimators become increasingly inflated (Figure S6). In addition, while it seems likely that the effect sizes of shared causal SNPs would be positively correlated across populations, the PESCA model assumes zero cross-population correlation in order to facilitate inference (Methods). We therefore perform simulations under an alternative model in which EAS and EUR effect sizes at shared causal SNPs are positively correlated and find that our estimates of the genome-wide numbers of shared and population-specific causal SNPs become increasingly inflated and deflated, respectively, as the correlation increases from 0 to 1 (Figure S7).

Next, we use the estimated genome-wide proportions of population-specific/shared causal SNPs to evaluate per-SNP posterior probabilities of being causal in a single population (EAS only or EUR only) or in both populations (Methods). For each of the three causal configurations of interest (EAS only, EUR only, and shared), we observe an increase in the average correlation between the per-SNP posterior probabilities and the true causal status vector for that configuration as 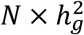 increases and as the total number of causal SNPs decreases (i.e. as per-SNP causal effect sizes increase) (Figures S8-S9). As expected, as the simulated proportion of shared causal SNPs increases, the average correlation between the posterior probabilities and true causal status vectors increases for the shared causal configuration and decreases for the population-specific causal configurations (Figures S8-S9). Since we did not have access to individual-level genotypes sampled from an ancestral group with shorter LD blocks (e.g., African-ancestry individuals), we use the EAS and EUR LD scores of each SNP as proxies for the strength of LD in the region housing the SNP to investigate the impact of population-specific LD patterns on the per-SNP posterior probabilities. Among the true causal SNPs (shared or population-specific), the posterior probabilities are relatively invariant to the magnitude of the EAS and EUR LD scores (Figure S10). In other words, under the PESCA framework, power to detect a given true causal SNP does not depend on its LD score in either population. Restricting to a set of “high-posterior SNPs” (defined here as SNPs with posterior probability greater than some threshold *t*), we investigate whether PESCA systematically misclassifies SNPs based on the magnitude of their LD scores. Again, we observe that the average EAS and EUR LD scores do not vary significantly between the true and false positive classifications (Table S1). We then assessed whether our proposed statistics for testing for enrichment of population-specific/shared causal SNPs in functional annotations (Methods) are well-calibrated under the null hypothesis of no enrichment. Overall, when both population-specific and shared causal SNPs are drawn at random, the enrichment test statistics are conservative at different levels of polygenicity and GWAS power 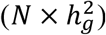, irrespective of whether in-sample LD or external reference LD is used (Figures S11-S16).

Finally, we evaluated the computational efficiency of each stage of inference. In the first stage of inference – estimating genome-wide proportions of population-specific/shared causal SNPs – the maximization step of the EM algorithm uses Gibbs sampling to efficiently sample from the posterior of the causal status vectors (Supplemental Note). We set both the number of burn-in iterations and the number of samples to 5,000 for the MCMC within the maximization step and found that the overall EM typically converged within 200 iterations (Figures S17-S19). Run-time per EM-iteration increases with the number of causal SNPs (Figure S20); for example, in simulations with a total of 8,589 SNPs, when the maximum number of EM iterations was set to 200, PESCA took an average of 90 minutes to obtain estimates in simulations with 20 randomly selected causal variants and 360 minutes in simulations with 100 randomly selected causal SNPs. This is expected because the likelihood function being maximized is proportional to the Bayes factor of only the causal SNPs (Supplemental Note). In the second stage of inference – evaluating posterior probabilities for each SNP – we set both the number of burn-in iterations and the number of samples to 5,000 for the MCMC and, to ensure stable estimates of the posterior probability, we report the average posterior probability from 20 iterations of the Gibbs sampling procedure (Supplemental Note). The average run-time was 5 minutes in simulations with 20 causal variants and 28 minutes in simulations with 100 causal variants (Figure S20). We note that both stages of inference can be parallelized to decrease run time.

### Expected genome-wide proportions of shared causal SNPs for 9 complex traits

We obtained publicly available GWAS summary statistics for 9 (non-independent) complex traits and diseases in individuals of EAS and EUR ancestry (average N_EAS_ = 94,621, N_EUR_ = 103,507) (Table 1) and applied PESCA to estimate the genome-wide proportions of population-specific/shared common causal SNPs (Methods). To ensure convergence, we applied 750 EM iterations for each trait (Figures S21-S23). Across the 9 traits, the estimated proportions of common causal SNPs in each population (the sum of the numbers of population-specific and shared causal SNPs) are consistent with previously reported estimates of polygenicity in single populations^7,8,55,75,76^. For example, we estimate that approximately 10% of common SNPs have nonzero effects on BMI in both EAS and EUR and that 2-3% have nonzero effects on the lipids traits (Table 1). The low estimates for major depressive disorder and rheumatoid arthritis may be explained in part by their small GWAS sample sizes. While there is heterogeneity in the estimated proportions of shared causal SNPs across the 9 traits, we find that most common causal SNPs are shared between the populations, consistent with findings from previous studies^33^. For example, for BMI, we estimate that approximately 96% of common causal SNPs in each population are also causal in the other; for total cholesterol (TC), we estimate that 73% of common causal SNPs in EAS and 77% of those in EUR are shared by both populations (Table 1).

### High-posterior SNPs are distributed nearly uniformly across the genome

We define 1,368 regions that are approximately LD-independent in both populations and estimate the posterior expected numbers of population-specific/shared causal SNPs in each region (Methods). For all 9 traits, high-posterior SNPs for both the population-specific and shared causal configurations are spread nearly uniformly across the genome (Figure 3, Figures S24-S31). For example, mean corpuscular hemoglobin (MCH) harbored, on average, 0.68 (S.D. 0.42) EAS-specific, 0.53 (S.D. 0.40) EUR-specific, and 2.19 (S.D. 1.46) shared high-posterior SNPs per region (Figure 3, Figure S29). Aggregating posterior probabilities by chromosome, we find that the posterior expected numbers of EAS-specific, EUR-specific, and shared causal SNPs per chromosome are highly correlated with chromosome length (Figures S32-S34), recapitulating previous findings based on regional SNP-heritability^55,60^.

**Figure 3:**
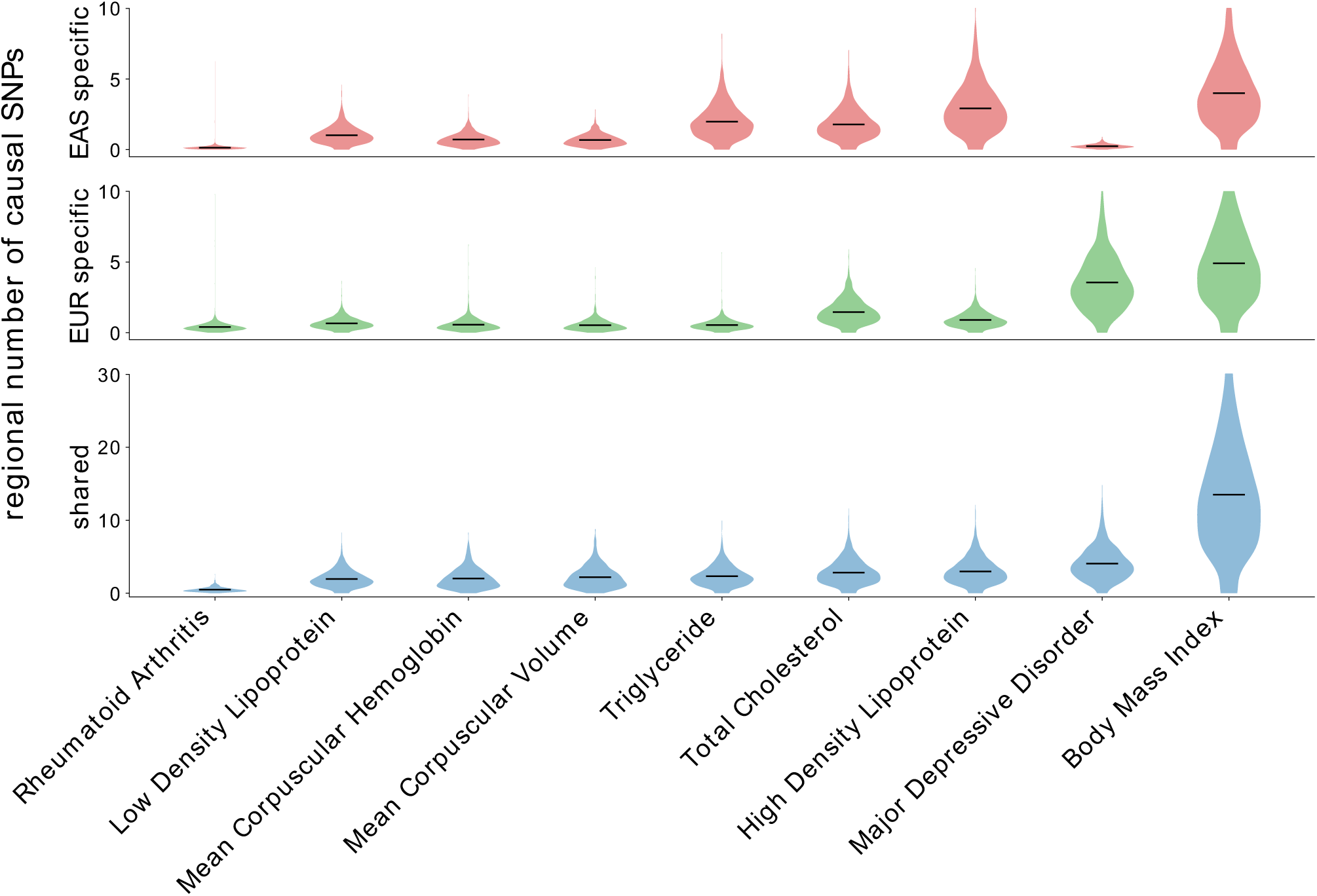
Distributions of the numbers of population-specific/shared causal SNPs across 1,368 regions that are approximately independent in both EAS and EUR. Each violin plot represents the distribution of the posterior expected number of population-specific or shared causal SNPs per region; details on how the regions were defined can be found in the Methods. For a single region, the posterior expected number of SNPs in a given causal configuration is estimated by summing, across all SNPs in the region, the per-SNP posterior probabilities of having that causal configuration (Methods). The dark lines mark the means of the distributions. The traits are sorted on the x-axis by the average number of shared high-posterior SNPs per region.

### Distributions of high-posterior SNPs across GWAS risk regions

We aggregate per-SNP posterior probabilities within GWAS risk regions that are EAS-specific, EUR-specific, or shared by both populations and find that most GWAS risk regions harbor two or more shared high-posterior SNPs (Figure 4, Figures S35-S39), concordant with previous findings on allelic heterogeneity of complex traits^55,77,78^. On average across the 9 traits, we observe a 2.8x enrichment of shared high-posterior SNPs in population-specific GWAS risk regions relative to the genome-wide background. For example, for mean corpuscular hemoglobin (MCH), the EAS-specific and EUR-specific GWAS risk regions harbor an average of 3.0 (S.D. 1.7) and 3.3 (S.D. 1.5) shared high-posterior SNPs per region, respectively, whereas the average number of shared high-posterior SNPs per region across all regions is 2.0 (S.D. 1.3) (Figure 4). While BMI, the blood traits (MCH and MCV), and rheumatoid arthritis have similar numbers of EAS-specific and EUR-specific high-posterior SNPs in their population-specific GWAS risk regions, the lipids traits (HDL, LDL, total cholesterol and triglycerides) have significantly more EAS-specific high-posterior SNPs in all GWAS risk regions (Figure 4, Figures S35-S39).

**Figure 4:**
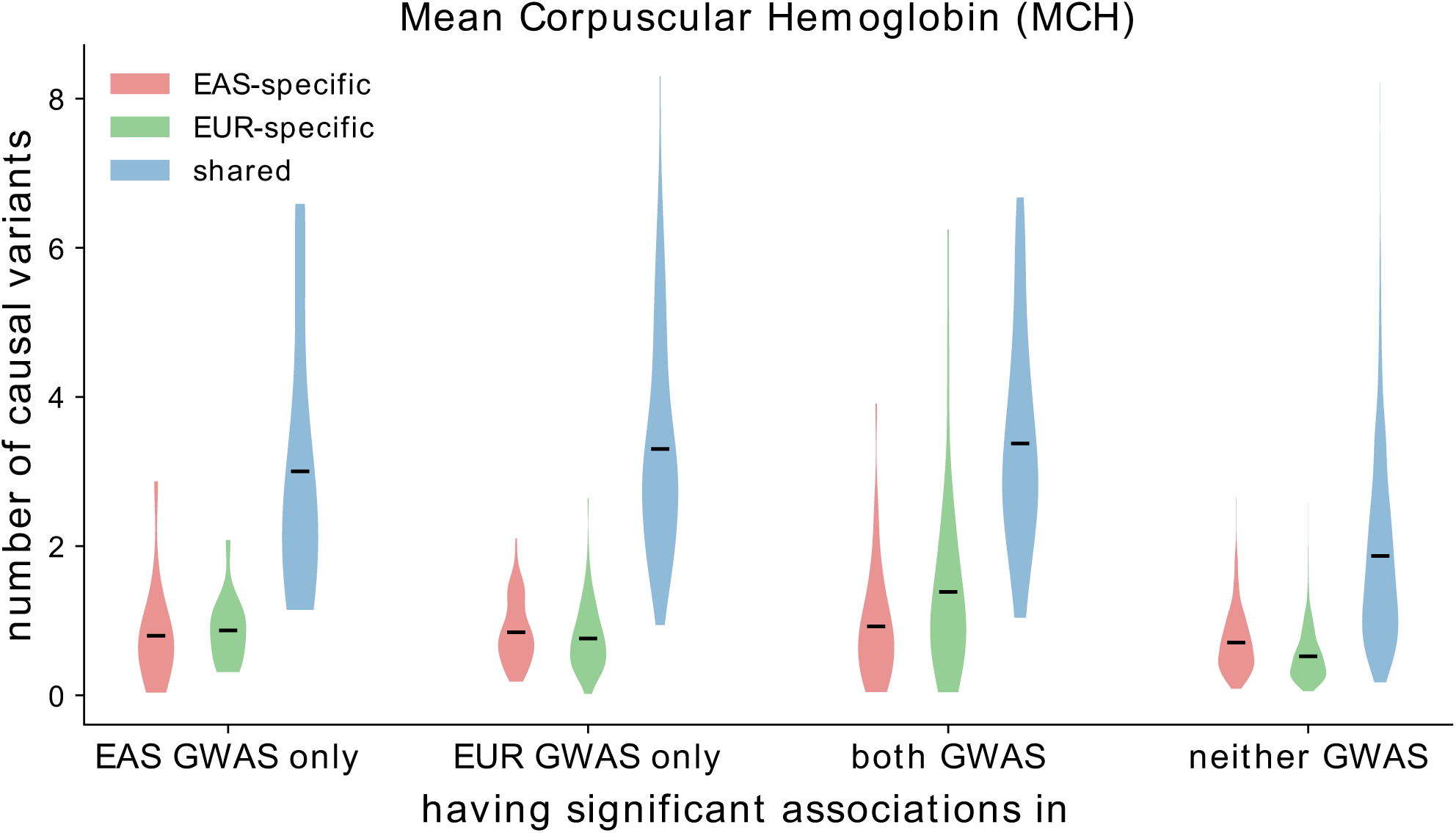
Distributions of the numbers of population-specific/shared causal variants at GWAS risk regions for mean corpuscular hemoglobin (MCH). Each violin plot represents the distribution of the posterior expected number of population-specific (red/green) or shared (blue) causal SNPs at regions with significant associations (*p*_*GWAS*_ < 5 × 10^−8^) in EAS GWAS only, EUR GWAS only, both EAS and EUR, and neither GWAS. The dark lines mark the means of the distributions.

For each causal configuration (EAS-specific, EUR-specific, or shared), we examine the effect sizes of high-posterior SNPs (posterior probability > 0.8) in EAS and EUR (Figure 5). Across the 9 traits, the majority of EAS-specific high-posterior SNPs are nominally significant (*p*_*GWAS*_ 5 × 10^−6^) either in the EAS GWAS only or in both GWASs. While five EUR-specific high-posterior SNPs are nominally significant in only the EAS GWAS, the majority are nominally significant either in the EUR GWAS only or in both GWASs. We observe strong correlations between the effect sizes in EAS and EUR for all three sets of high-posterior SNPs (Pearson *r*^2^ of 0.79 [EAS-specific], 0.73 [EUR-specific], and 0.80 [shared]) that are driven by SNPs that are nominally significant in both GWASs (Figure 5). Taken together, these results suggest that most population-specific GWAS risk regions harbor shared causal variants that are undetected in the other population due to heterogeneity in LD structures, allele frequencies, and/or GWAS sample sizes^33^.

**Figure 5:**
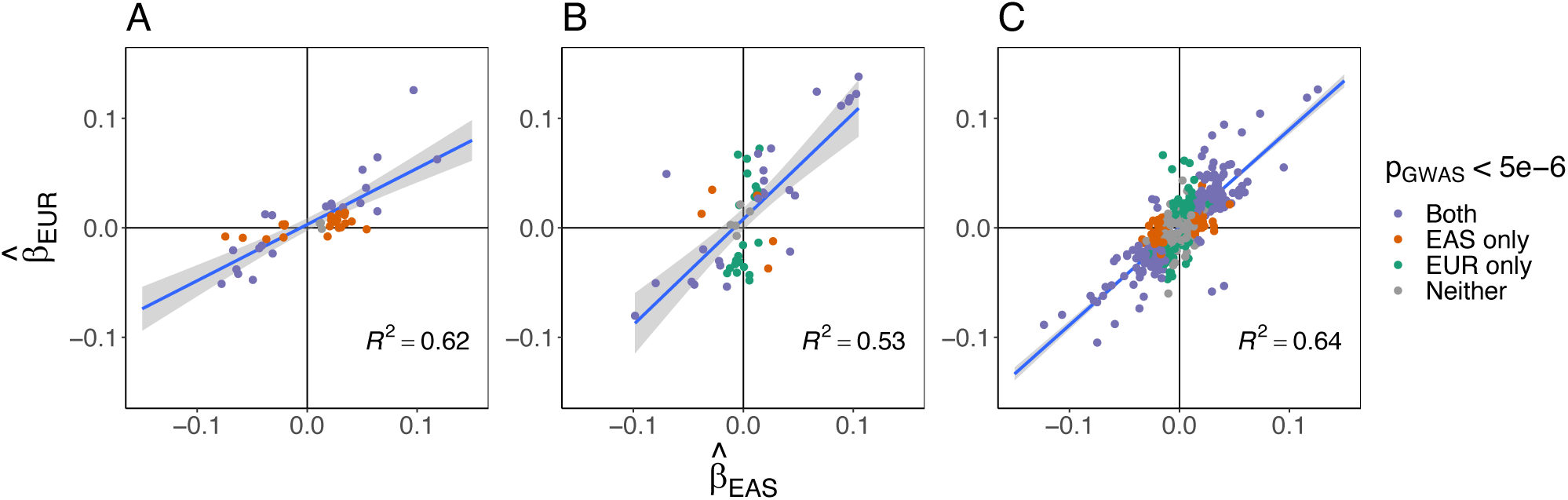
Marginal regression coefficients of high-posterior SNPs for 9 complex traits. Each plot corresponds to one of the three causal configurations of interest: EAS-specific (A), EUR-specific (B), and shared (C). Each point represents a SNP with posterior probability > 0.8 for a single trait. The x-axis and y-axis mark the marginal regression coefficients in the EAS-ancestry GWAS and EUR-ancestry GWAS, respectively. The colors indicate whether the SNP is nominally significant (*p*_*GWAS*_ < 5 × 10^−6^) in both GWASs (purple), the EAS GWAS only (orange), the EUR GWAS only (green), or in neither GWAS (gray). The gray band marks the 95% confidence interval of the regression line.

### Enrichment of high-posterior SNPs near genes expressed in trait-relevant tissues

Motivated by recent work that found enrichment of SNP-heritability in regions near genes that are “specifically expressed” in trait-relevant tissues and cell types (referred to as “SEG annotations”), we tested for enrichments of population-specific and shared causal SNPs in the same 53 tissue-specific SEG annotations^61^. For a given causal configuration, the enrichment of causal SNPs in an annotation is defined as the ratio between the posterior and prior expected numbers of causal SNPs in the annotation (Methods). For 8 of the 9 traits, we find significant enrichment of shared high-posterior SNPs in at least one SEG annotation (*P*-value < 0.05/53 to correct for 53 tests per trait) (Figures S40-S44). All SEG annotations with significant enrichments of population-specific high-posterior SNPs are also enriched with shared high-posterior SNPs for the same trait, providing additional evidence that many signatures of population-specific genetic architecture are induced by population-specific LD and allele frequencies rather than distinct genetic etiologies. We do not find enrichment of any high-posterior SNPs in any SEG annotation for major depressive disorder (MDD) (Figure S44), which could be due to low GWAS sample sizes (Table 1). Finally, for each SEG annotation, we obtain a meta-analyzed transethnic SNP-heritability enrichment by computing the inverse-variance weighted average of the EAS and EUR SNP-heritability enrichments (which are obtained separately using stratified LD score regression^13,16^). We observe a strong correlation between the meta-analyzed SNP-heritability enrichments and the enrichments of shared high-posterior SNPs (Figure 6), suggesting that SNP-heritability enrichments are largely driven by many low-effect SNPs rather than a small number of high-effect SNPs.

**Figure 6:**
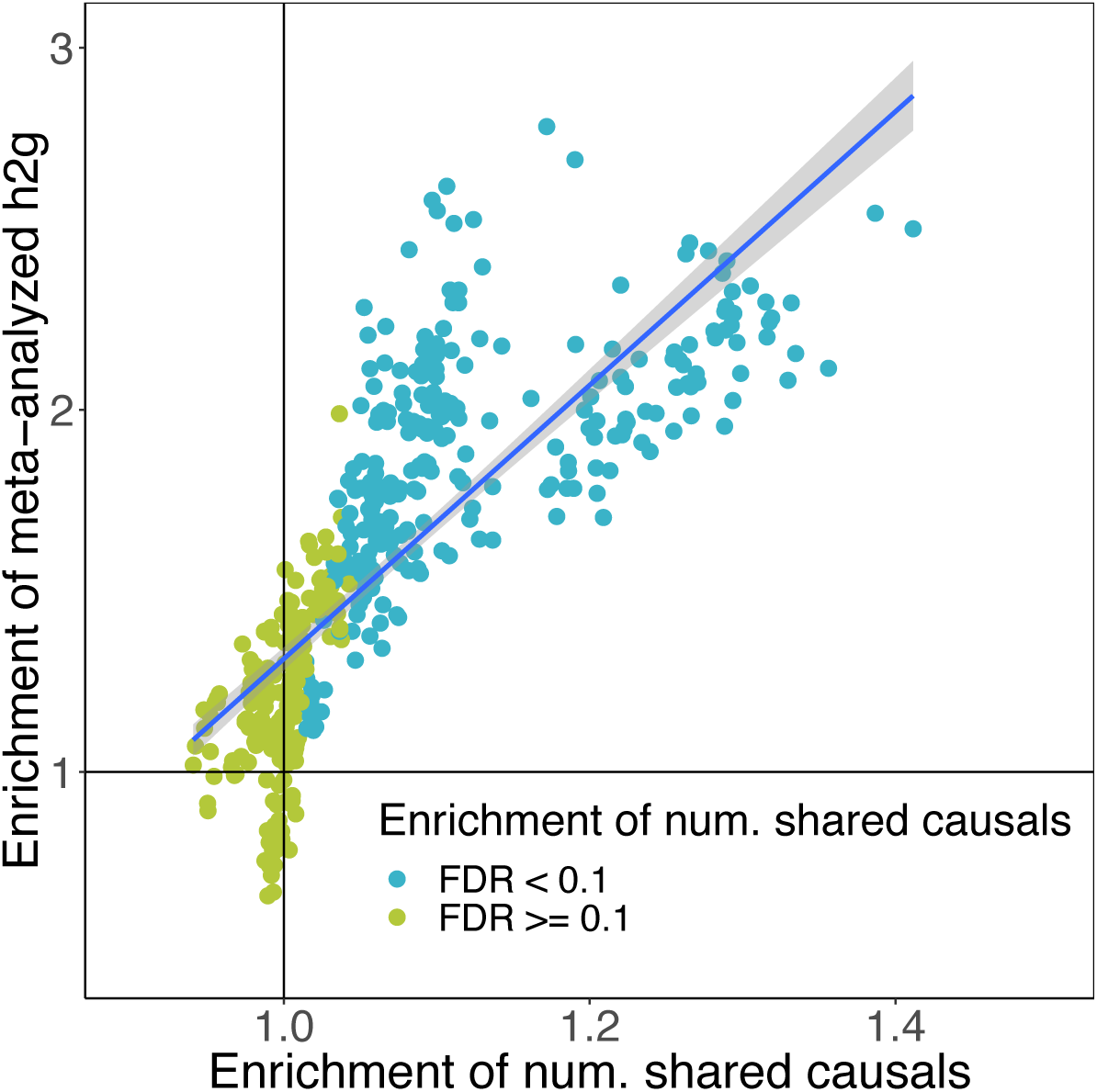
Enrichments of shared high-posterior SNPs in 53 tissue-specific functional categories are highly correlated with SNP-heritability enrichments. Each point represents a trait-tissue pair; each tissue-specific functional category represents a set of genes that are “specifically expressed” in one of 53 GTEx tissues (53 SEG annotations). The x-axis is the enrichment of shared high-posterior SNPs in the SEG annotation obtained from PESCA. The y-axis is the meta-analyzed transethnic SNP-heritability explained by the SEG annotation, defined as the inverse-variance weighted average of the EAS and EUR SNP-heritability enrichments (obtained separately using stratified LD score regression). The points are colored by whether the trait has a statistically significant enrichment of shared high-posterior SNPs in the corresponding SEG annotation (FDR < 0.1). Enrichment estimates and standard errors for each trait-tissue pair can be found in Figures S40-S44.

## Discussion

We have presented PESCA, a method for estimating the genome-wide proportions of SNPs with nonzero effects in a single population (population-specific) or in two populations (shared) from GWAS summary statistics and estimates of LD. We applied PESCA to EAS and EUR GWAS summary statistics for 9 complex traits and find that, while the lipids traits have significantly more EAS-specific common causal SNPs compared to the remaining traits, the majority of common causal SNPs are shared by both populations. Regions that harbor statistically significant GWAS associations for one population are enriched with SNPs with high-posterior probability of being causal in both populations; moreover, high-posterior SNPs (posterior probability > 0.8 for any causal configuration) have highly correlated effect sizes in EAS and EUR, recapitulating results of previous studies^33^. For all traits except MDD, we identify tissue-specific SEG annotations^66^ enriched with shared high-posterior SNPs and observe that all SEG annotations enriched with population-specific high-posterior SNPs are a subset of those enriched with shared high-posterior SNPs. Taken together, our results indicate that most population-specific GWAS risk regions contain shared common causal SNPs that are undetected in the second population due to differences in LD or allele frequencies. This suggests that localizing shared components of genetic architecture and explicitly correcting for population-specific LD and allele frequencies may help improve transferability of results from well-powered European-ancestry studies to other understudied populations. Based on the simulation results in Figure S1 (in which 100% of causal SNPs are shared) and our estimates of SNP-heritability for the traits in Table 1, we recommend applying PESCA to summary statistics for which the *effective per-SNP sample size*, 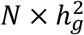 divided by the number of causal SNPs, is at least 3 for both GWASs. For a typical quantitative trait (e.g., Table 1), this corresponds to a total effective sample size of approximately 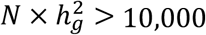.

We conclude by discussing the caveats and limitations of our analyses. First, the estimated proportions of causal SNPs must be interpreted with caution as they can be influenced by gene-environment interactions. For example, if a SNP has a nonzero effect on a trait only in the presence of environmental factors that are specific to EAS-ancestry individuals, PESCA will interpret that SNP as an EAS-specific causal SNP even though it would have a nonzero effect in Europeans in the presence of the same environmental factors.

Second, we chose to analyze a set of traits that were present in both the UK Biobank and Biobank Japan and for which GWAS summary statistics were publicly available. Since most publicly available summary statistics of large-scale GWAS are meta-analyses of smaller studies, in-sample LD is often unavailable. While PESCA with in-sample LD is relatively robust to differential GWAS power, with external LD, performance decreases when the GWAS effective sample sizes differ by more than a factor of 2x. We note, however, that for the real traits analyzed in this work, effective sample size differs by a maximum factor of 2x (mean corpuscular hemoglobin; Table 1). Additionally, PESCA currently cannot be applied to admixed populations if in-sample LD is unavailable. An extension of PESCA to properly account for external/noisy estimates of LD would thus increase its utility; we defer a thorough investigation of this to future work. In parallel, in light of ongoing efforts at several institutions to establish biobanks^69,70,79–81^, we believe that well-powered GWASs (with in-sample LD) will become increasingly available for diverse and admixed populations. Another challenge is that many publicly available summary statistics were computed from fixed-effect meta-analyses or linear mixed models. Since the PESCA model is defined with respect to GWAS marginal effects estimated by ordinary least squares (OLS) regression, it is unclear whether PESCA is sensitive to non-OLS association statistics, which have different statistical properties; we defer a thorough investigation of this to future work.

Third, we restricted our analyses to SNPs with MAF > 5% in both populations to reduce noise in the LD matrices estimated from external reference panels. Consequently, the estimates we report in this work do not capture effects of low frequency or rare variants that are not well-tagged by common SNPs. Furthermore, since most common variants are shared across continental populations and rarer variants tend to localize among closely related populations^73^, our study design undersamples population-specific causal variants. We note, however, that lower MAF thresholds can be used if in-sample LD is available. We also note that for the purpose of improving transferability of polygenic risk scores (PRS) across populations, prediction accuracy depends largely on the accuracy of the PRS weights at common SNPs (the average per-SNP contribution to total SNP-heritability is larger for common SNPs than for low frequency or rare variants^11^).

Finally, PESCA can be sensitive to model misspecification. For computational efficiency, PESCA relies on having regions that are approximately LD-independent in both populations; if there is LD leakage between regions, the estimated proportions of causal SNPs will be biased. We therefore recommend defining LD blocks for each pair of populations one analyzes. Similarly, to facilitate inference, PESCA does not explicitly model cross-population correlations of effect sizes at shared causal variants; we conjecture that modeling these correlations can further improve performance.

## Acknowledgements

We are grateful to Alkes L. Price and Steven Gazal for helpful discussions that greatly improved the quality of this manuscript. We also thank Na Cai, Sriram Sankararaman, Jonathan Flint, and the UK Biobank (application #33297) for providing resources that made this work possible. This work was funded in part by the National Institutes of Health (NIH) under awards R01HG009120, R01MH115676, U01CA194393, T32NS048004, T32MH073526, and T32HG002536.

## Declaration of Interests

The authors declare no competing interests.

## Web Resources

GIANT consortium GWAS summary statistics: http://portals.broadinstitute.org/collaboration/giant

Biobank Japan GWAS summary statistics: http://jenger.riken.jp/en/result

GWAS summary statistics for hematological traits: http://www.bloodcellgenetics.org

LD score regression: https://github.com/bulik/ldsc

PLINK 1.9: https://www.cog-genomics.org/plink/1.9/

Popcorn: https://github.com/brielin/Popcorn

PESCA: https://github.com/huwenboshi/pesca

Specifically expressed genes: https://data.broadinstitute.org/alkesgroup/LDSCORE/LDSC_SEG_ldscores

