## Supplemental Material for "Localizing components of shared transethnic genetic architecture of complex traits from GWAS summary data"

[1]Bioinformatics Interdepartmental Program, University of California, Los Angeles, CA
[2]Department of Epidemiology, Harvard T.H. Chan School of Public Health, Boston, MA
[3]Program in Medical and Population Genetics, Broad Institute of MIT and Harvard, Cambridge, MA
[4]Department of Computer Science, University of California, Los Angeles, CA
[5]Department of Human Genetics, David Geffen School of Medicine, University of California, Los Angeles, CA
[6]Center for Genetic Epidemiology, Department of Preventive Medicine, Keck School of Medicine, University of Southern California, Los Angeles, CA
[7]Department of Biological Sciences, Florida International University, Miami, FL
[8]Department of Biomedical Engineering, Boston University, Boston, MA
[9]Department of Pathology and Laboratory Medicine, David Geffen School of Medicine, University of California, Los Angeles, CA
[10]Department of Computational Medicine, David Geffen School of Medicine, University of California, Los Angeles, CA
[*]These authors contributed equally to this work

#### 1 Supplemental Figures

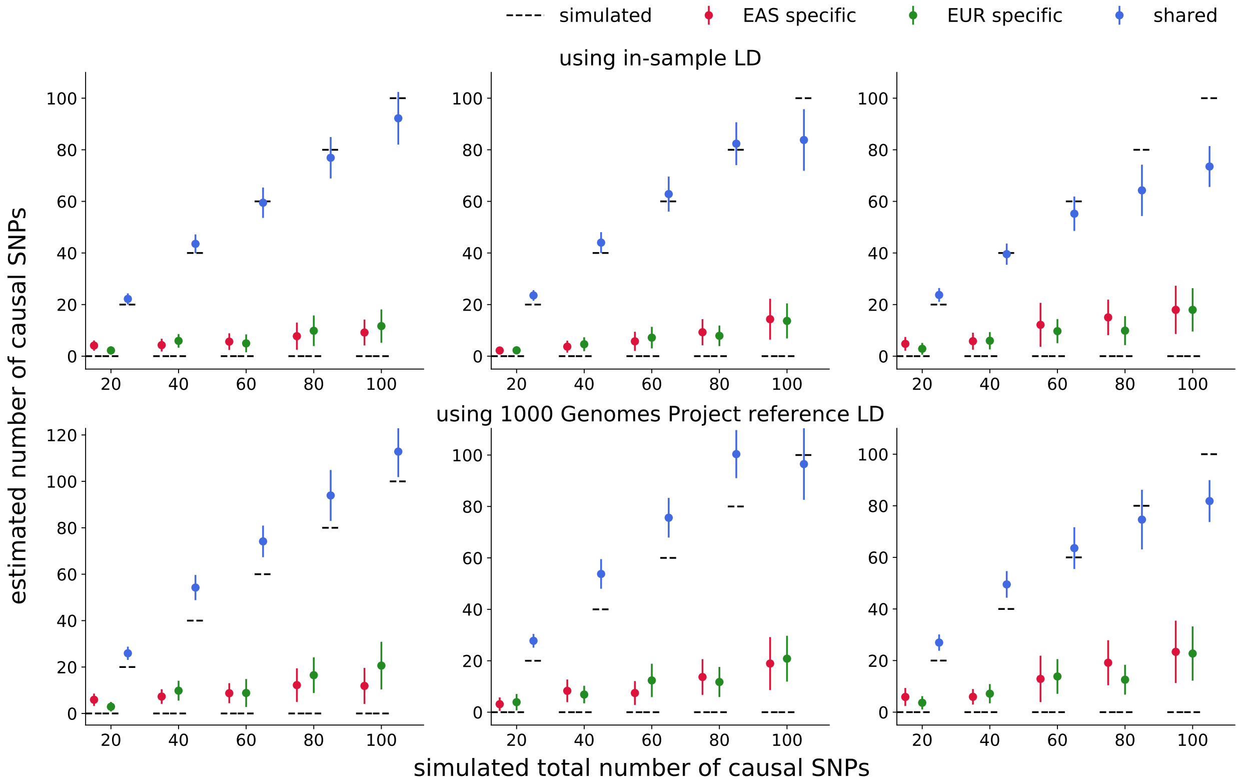

Figure S1: **PESCA estimators for the genome-wide numbers of population-specific/shared causal SNPs when 100% of causal variants are shared.** We simulated 20 to 100 causal variants per population (x-axis), all of which were shared by both populations. We set the product of SNP-heritability and sample size of the GWAS to 500 (left column), 375 (middle column), and 250 (right column), which correspond to per-SNP effective sample sizes (N x per-snp variance) that decrease from 25 to 5 (left), 18.75 to 3.75 (middle), and 12.5 to 2.5 (right). Each dot represents the mean across 25 simulations and error bars represent $\pm$1.96 s.e.m.

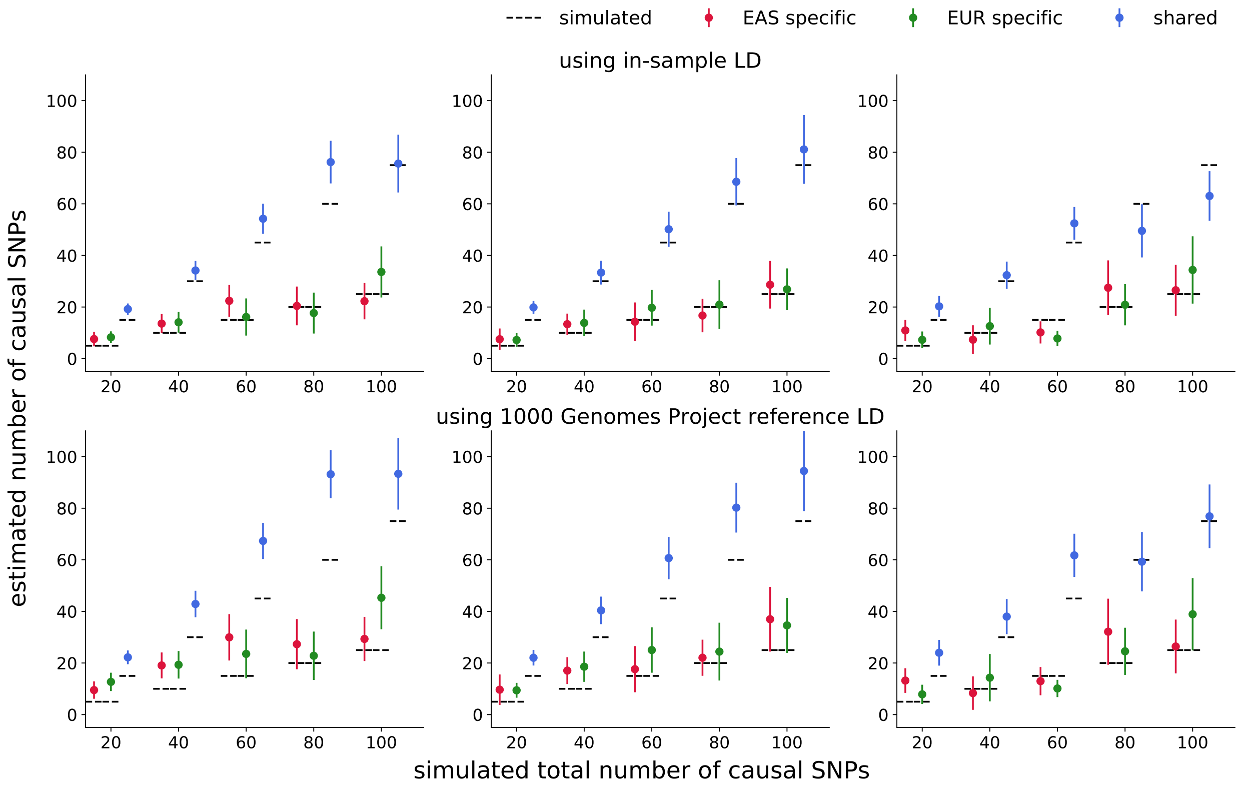

Figure S2: **PESCA estimators for the genome-wide numbers of population-specific/shared causal SNPs when 75% of causal variants are shared.** We simulated 20 to 100 causal variants per population (x-axis), 75% of which were shared; the remaining 25% were population-specific. We set the product of SNP-heritability and sample size of the GWAS to 500 (left column), 375 (middle column), and 250 (right column). Each dot represents the mean across 25 simulations and error bars represent $\pm 1.96$ s.e.m.

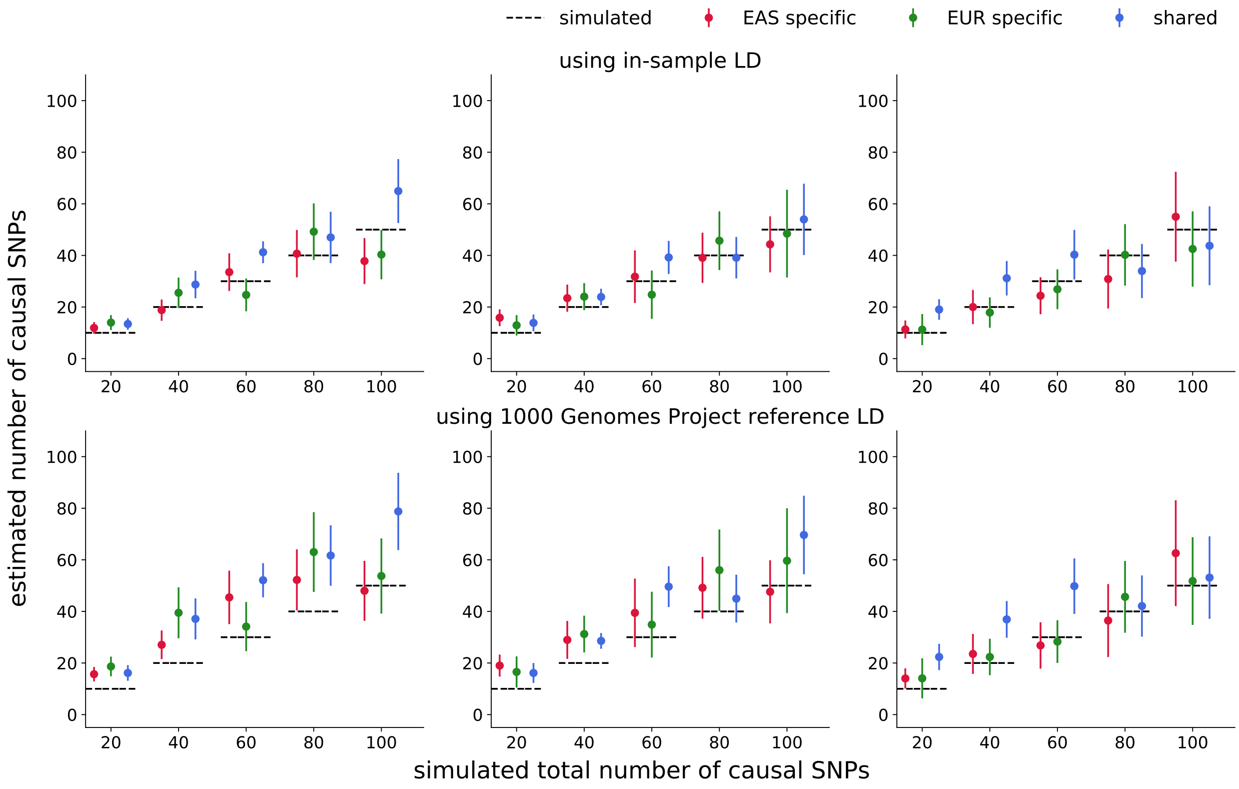

Figure S3: **PESCA estimators for the genome-wide numbers of population-specific/shared causal SNPs when 50% of causal variants are shared.** We simulated 20 to 100 causal variants per population (x-axis), 50% of which were shared; the remaining 50% were population-specific. We set the product of SNP-heritability and sample size of the GWAS to 500 (left column), 375 (middle column), and 250 (right column). Each dot represents the mean across 25 simulations and error bars represent $\pm 1.96$ s.e.m.

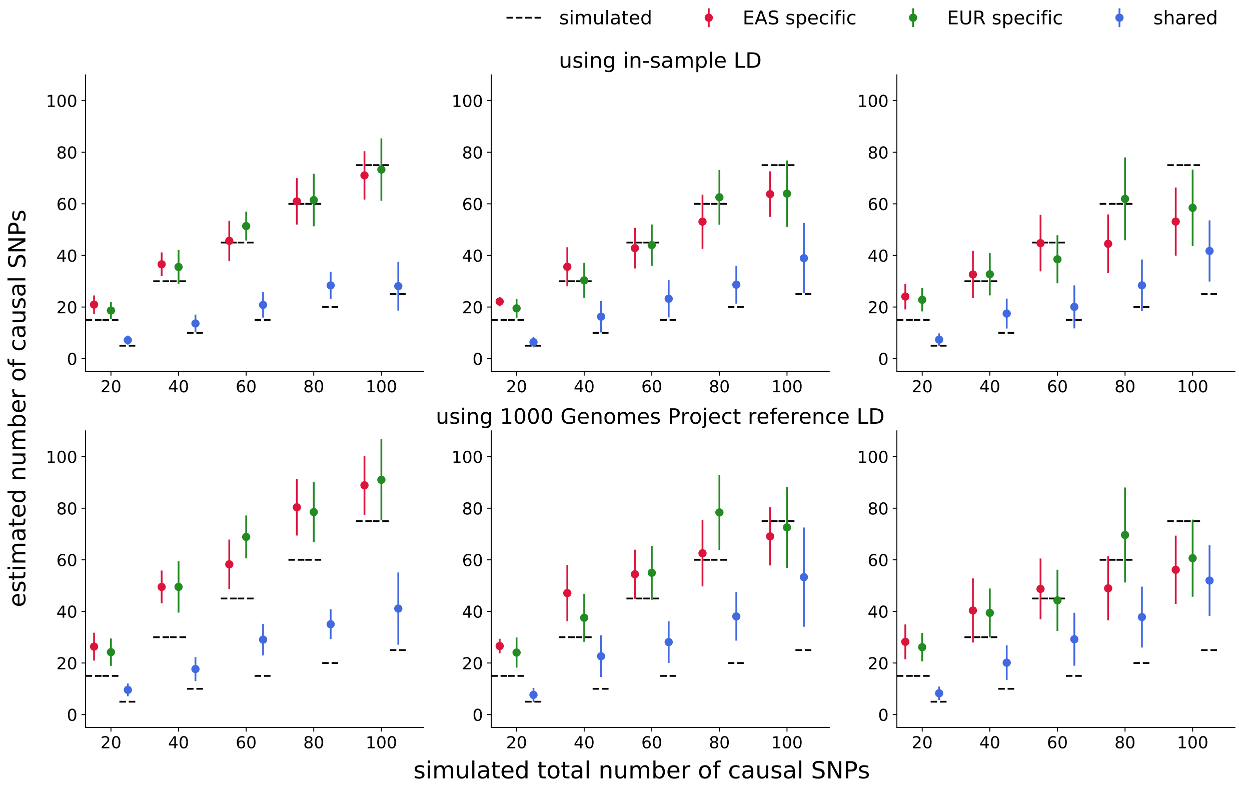

Figure S4: **PESCA estimators for the genome-wide numbers of population-specific/shared causal SNPs when 25% of causal variants are shared.** We simulated 20 to 100 causal variants per population (x-axis), 25% of which were shared; the remaining 75% were population-specific. We set the product of SNP-heritability and sample size of the GWAS to 500 (left column), 375 (middle column), and 250 (right column). Each dot represents the mean across 25 simulations and error bars represent $\pm 1.96$ s.e.m.

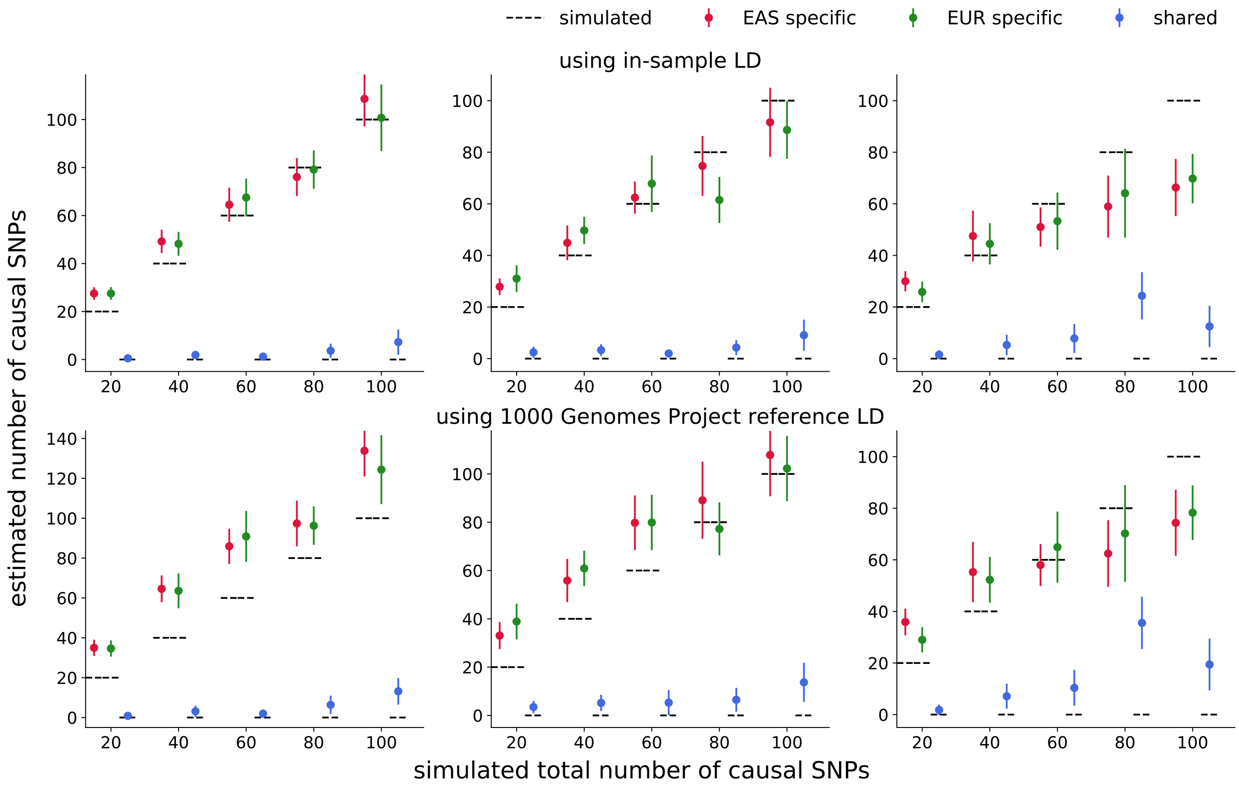

Figure S5: **PESCA estimators for the genome-wide numbers of population-specific/shared causal SNPs when 0% of causal variants are shared.** We simulated 20 to 100 causal variants per population (x-axis), all of which were population-specific. We set the product of SNP-heritability and sample size of the GWAS to 500 (left column), 375 (middle column), and 250 (right column). Each dot represents the mean across 25 simulations and error bars represent $\pm 1.96$ s.e.m.

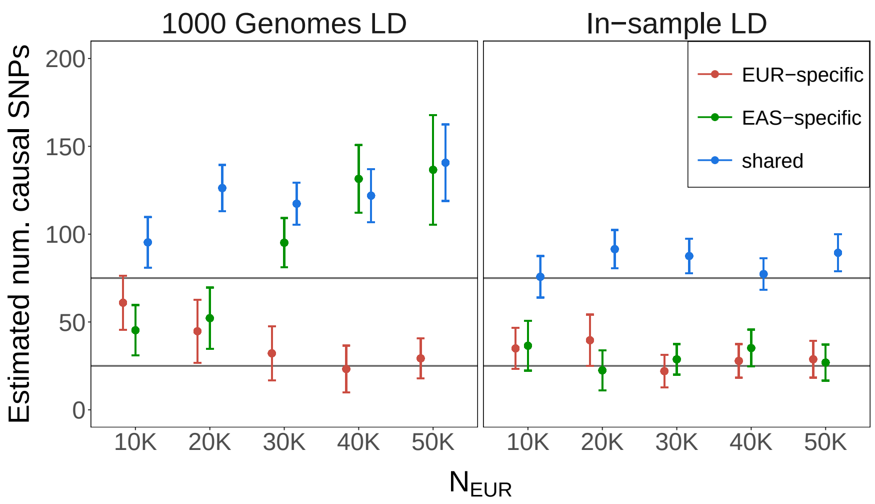

Figure S6: **Effect of differential effective sample size on PESCA estimates of the genome-wide numbers of population-specific/shared causal SNPs.** Total SNP-heritability was fixed to $h_g^2 = 0.05$ for both populations. $N_{EAS} = 10^4$ in all simulations; $N_{EUR}$ was varied from $1 \times 10^4$ to $5 \times 10^4$ (x-axis). Horizontal lines mark the number of shared (75), EAS-specific (25), and EUR-specific (25) causal SNPs. Each dot represents the mean across 25 simulations and error bars represent $\pm$1.96 s.e.m. The colors correspond to the estimators for the numbers of population-specific (red and green) and shared (blue) causal variants.

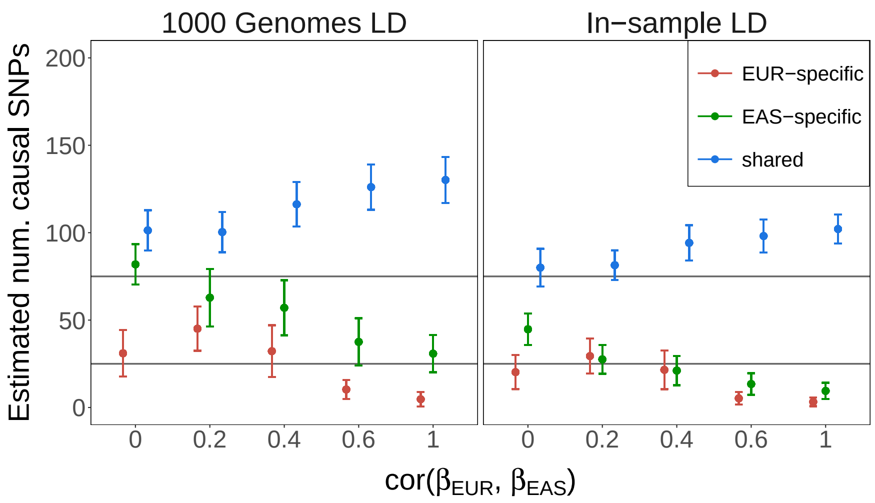

Figure S7: **Effect of cross-population correlation of causal effects on PESCA estimates of the genome-wide numbers of population-specific/shared causal SNPs.** Total SNP-heritability was fixed to $h_g^2 = 0.05$ for both populations. $N_{EAS} = 1 \times 10^4$ and $N_{EUR} = 2 \times 10^4$ in all simulations. Horizontal lines mark the number of shared (75), EAS-specific (25), and EUR-specific (25) causal SNPs. The correlation of effect sizes at causal SNPs was varied from 0 to 1 (x-axis). Each dot represents the mean across 25 simulations and error bars represent $\pm$1.96 s.e.m. The colors correspond to the estimators for the numbers of population-specific (red and green) and shared (blue) causal variants.

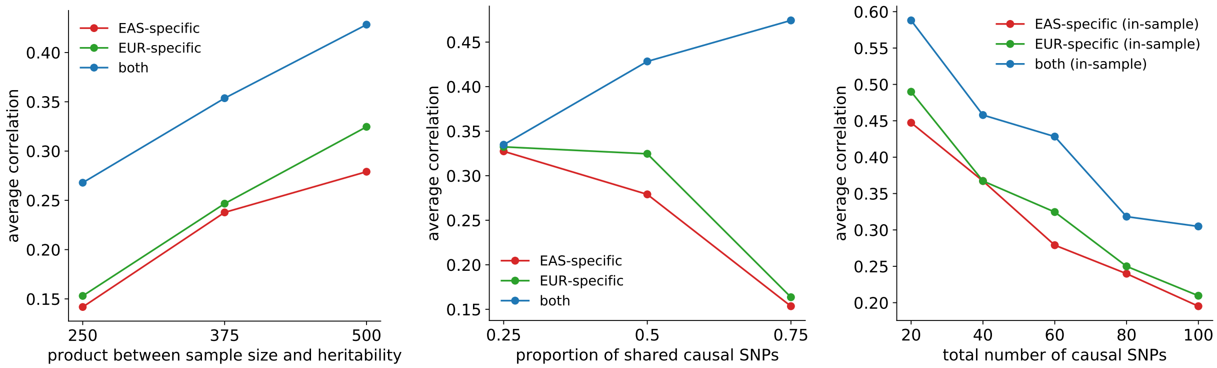

Figure S8: **Accuracy of PESCA posterior probabilities in simulations using in-sample LD.** Each point represents the average correlation (across 25 simulation replicates) between the vector of per-SNP posterior probabilities of causality and the vector of simulated causal statuses for one of the possible causal configurations (EAS-specific, EUR-specific, or both) as a function of $N \times h_g^2$ (left), the total number of causal SNPs in both populations (middle), and the proportion of shared causal SNPs (right). The correlations are calculated from a set of SNPs with MAF $>$ 5% in both populations that satisfy $r_{ij}^2 < 0.95$ for all pairs of SNPs ($i \neq j$) in both populations (Methods).

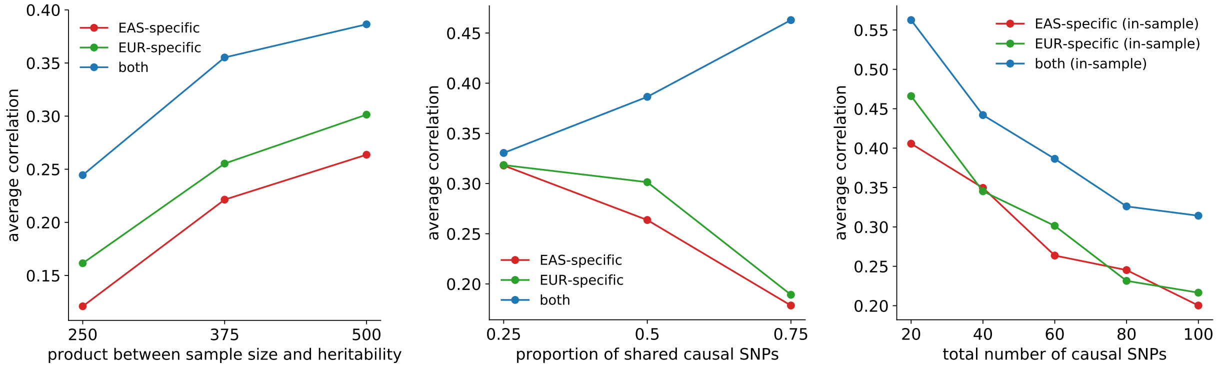

Figure S9: **Accuracy of PESCA posterior probabilities in simulations using external reference panel LD (1000 Genomes).** Each point represents the average correlation (across 25 simulation replicates) between the vector of per-SNP posterior probabilities of causality and the vector of simulated causal statuses for one of the possible causal configurations (EAS-specific, EUR-specific, or both) as a function of $N \times h_g^2$ (left), the total number of causal SNPs in both populations (middle), and the proportion of shared causal SNPs (right). The correlations are calculated from a set of SNPs with MAF $>$ 5% in both populations that satisfy $r_{ij}^2 < 0.95$ for all pairs of SNPs ($i \neq j$) in both populations (Methods).

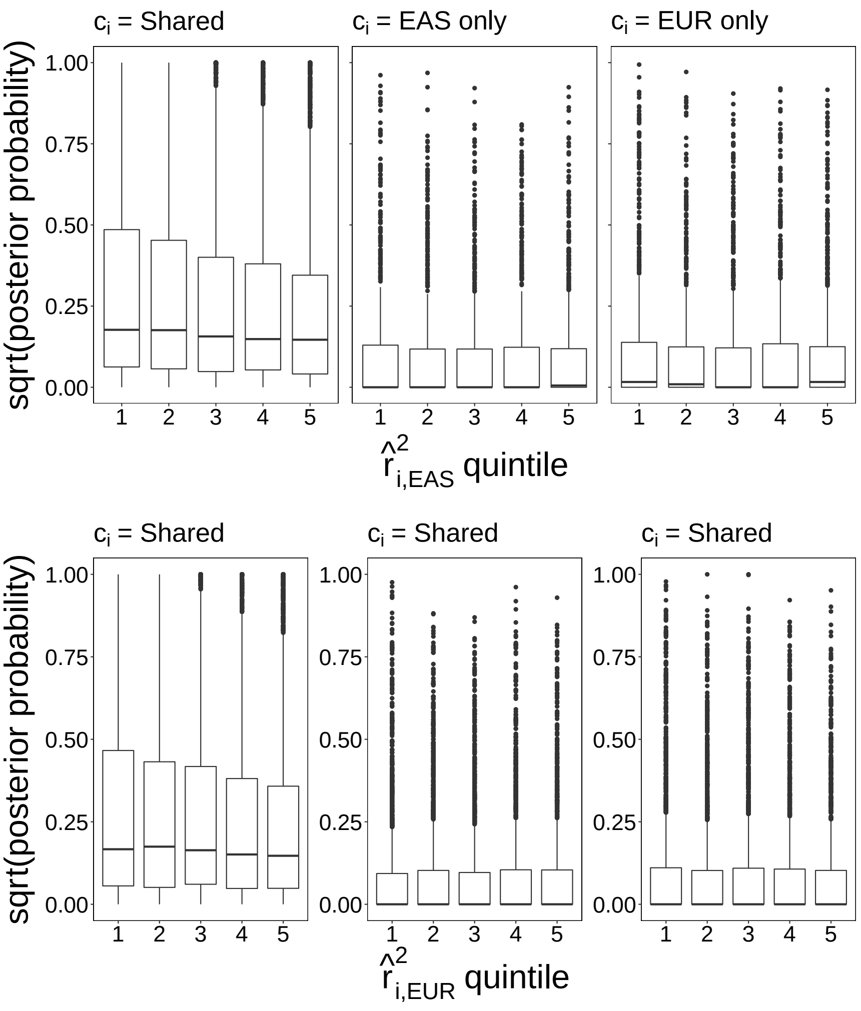

Figure S10: **Posterior probabilities of true causal SNPs with respect to LD score quintiles**. Total SNP-heritability was fixed to $h_g^2 = 0.05$ for both populations and $N_{EAS} = N_{EUR} = 10^4$. In each of the 200 simulation replicates, 75 shared (left panel), 25 EAS-specific (middle panel), and 25 EUR-specific (right panel) causal SNPs were drawn at random from 8,599 SNPs on chromosome 22. Each point in each boxplot represents a single true causal SNP. Each boxplot shows the distribution of per-SNP posterior probabilities for the corresponding correct causal configuration with respect to LD score quintiles in EAS (top) or EUR (bottom). We plot the square root of the posteriors to facilitate visualization.

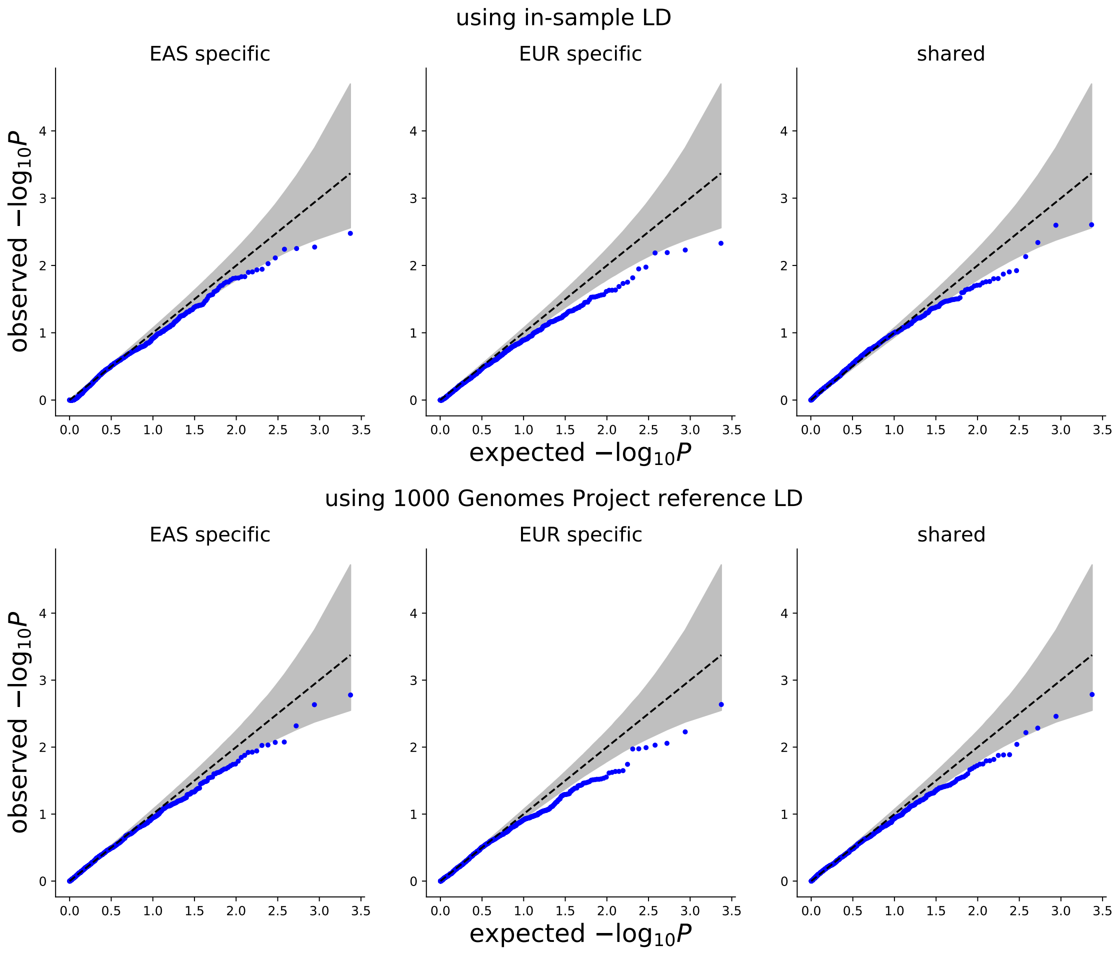

Figure S11: **Q-Q plot of p-values of the enrichments of population-specific/shared causal variants in SEG annotations[1] obtained using in-sample LD (top row) or ancestry-matched 1000 Genomes LD (bottom row).** We computed p-values from the enrichment test statistics of SEG annotations in 53 GTEx tissues from 25 null simulations, where we drew 25 EAS-specific, 25 EUR-specific, and 75 shared causal variants at random. In all simulations, we set $N \times h_g^2 = 500$ in both populations. Columns correspond to enrichment test statistics for the number of EAS-specific (left), EUR-specific (middle), or shared (right) causal variants.

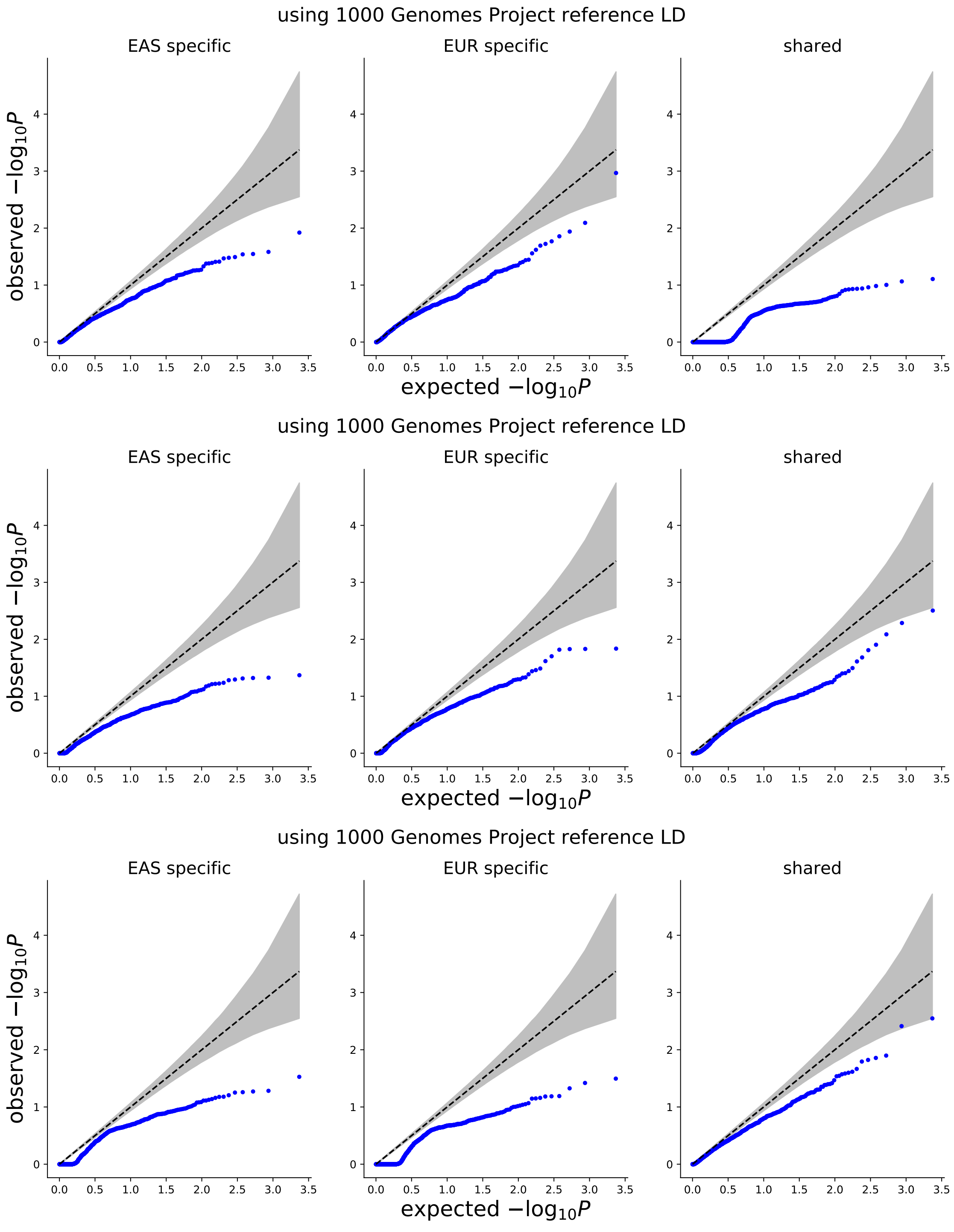

Figure S12: **Q-Q plot of p-values of the enrichments of population-specific/shared causal variants in SEG annotations[1] (20 causal variants per population).** We computed p-values for SEG annotations across 53 GTEx tissues from 25 null simulations, where we drew 20 causal variants at random for each population. In all simulations, we set $N \times h_g^2 = 500$ in both populations. The top, middle, and bottom rows represent results from simulations where 0% (top), 50% (middle), and 100% (bottom) of the causal SNPs were shared. All results were obtained using 1000 Genomes Project reference LD.

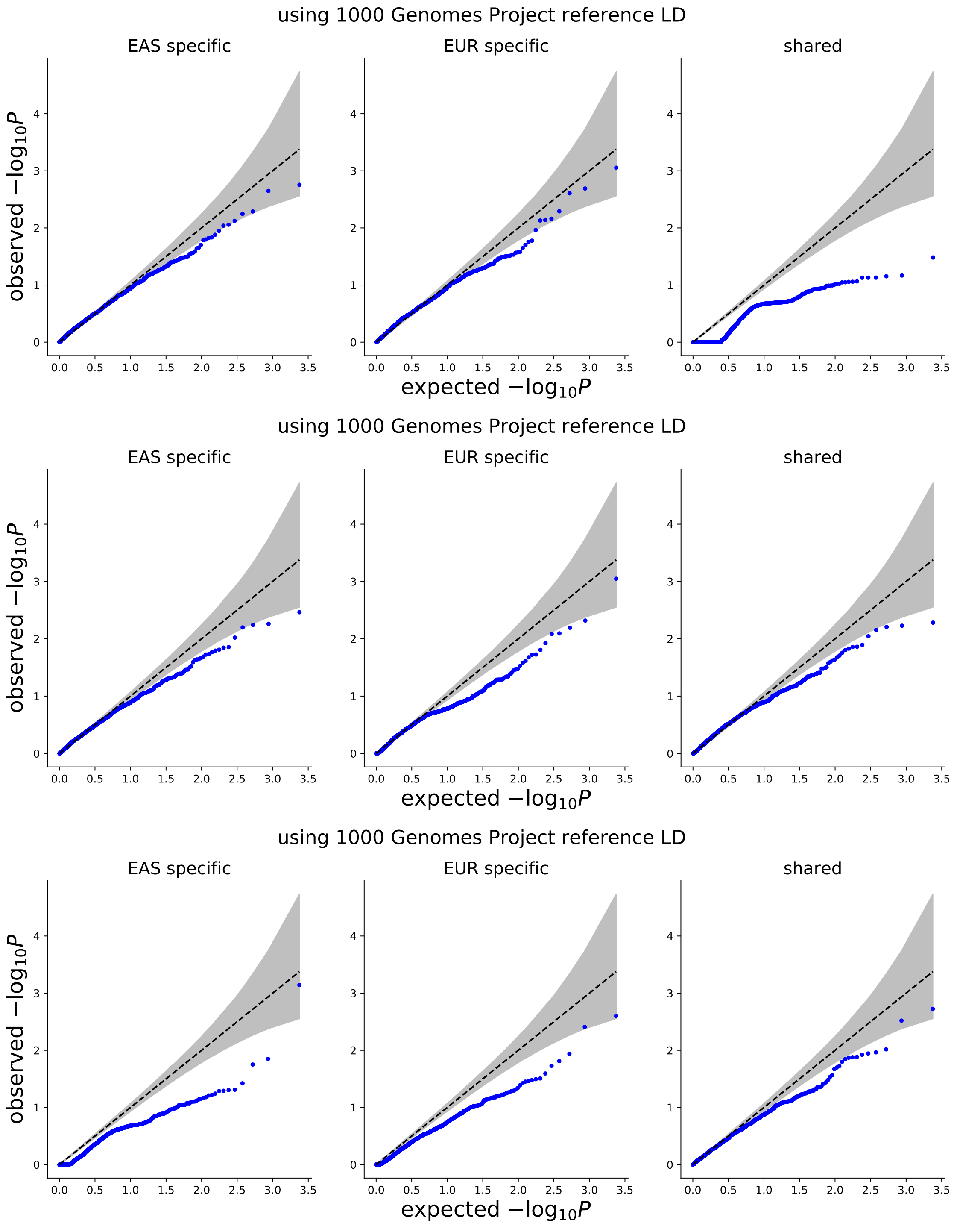

Figure S13: **Q-Q plot of p-values of the enrichments of population-specific/shared causal variants in SEG annotations[1] (60 causal variants per population).** We computed p-values for SEG annotations across 53 GTEx tissues from 25 null simulations, where we drew 60 causal variants at random for each population. In all simulations, we set $N \times h_g^2 = 500$ in both populations. The top, middle, and bottom rows represent results from simulations where 0% (top), 50% (middle), and 100% (bottom) of the causal SNPs were shared. All results were obtained using 1000 Genomes Project reference LD.

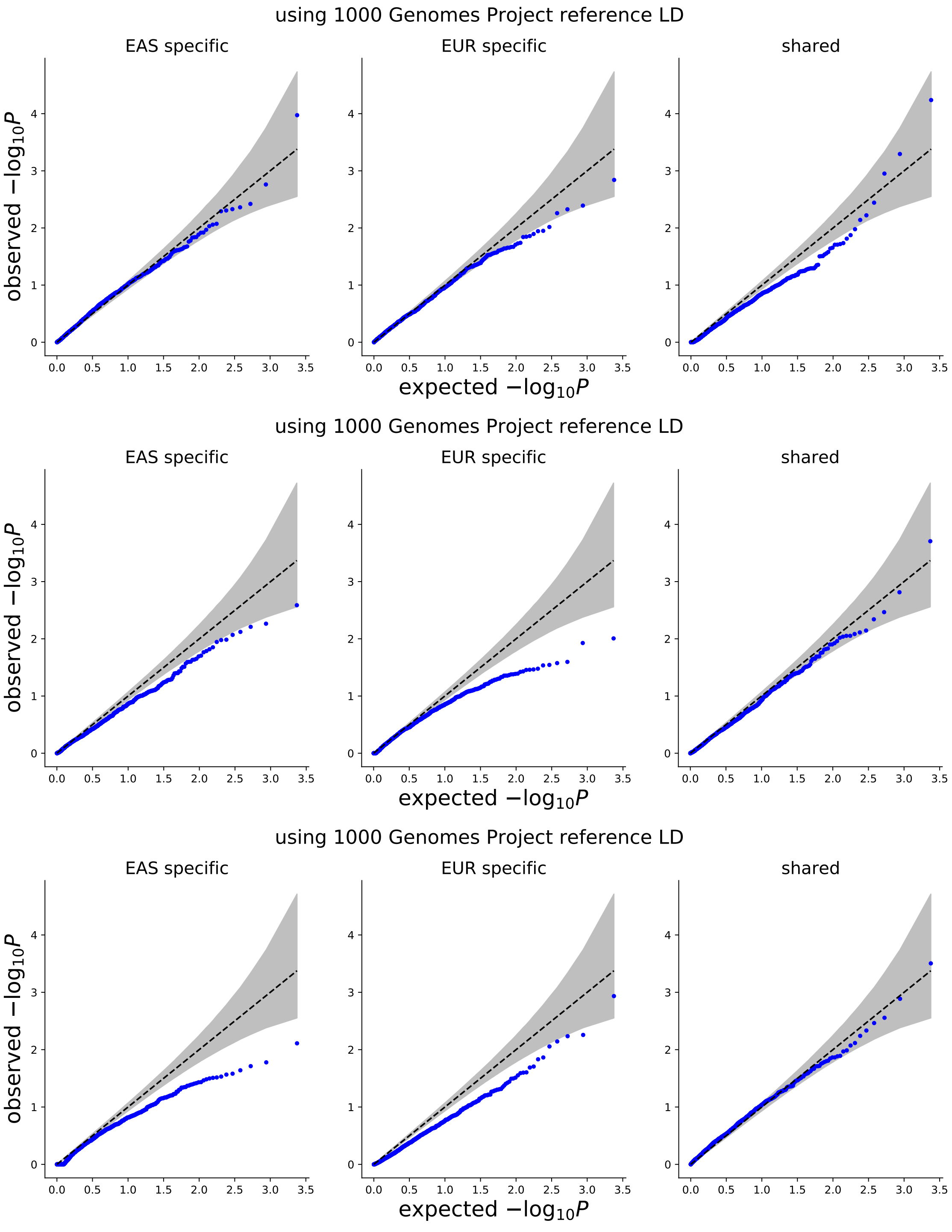

Figure S14: **Q-Q plot of p-values of the enrichments of population-specific/shared causal variants in SEG annotations[1] (100 causal variants per population).** We computed p-values for SEG annotations across 53 GTEx tissues from 25 null simulations, where we drew 100 causal variants at random for each population. In all simulations, we set $N \times h_g^2 = 500$ in both populations. The top, middle, and bottom rows represent results from simulations where 0% (top), 50% (middle), and 100% (bottom) of the causal SNPs were shared. All results were obtained using 1000 Genomes Project reference LD.

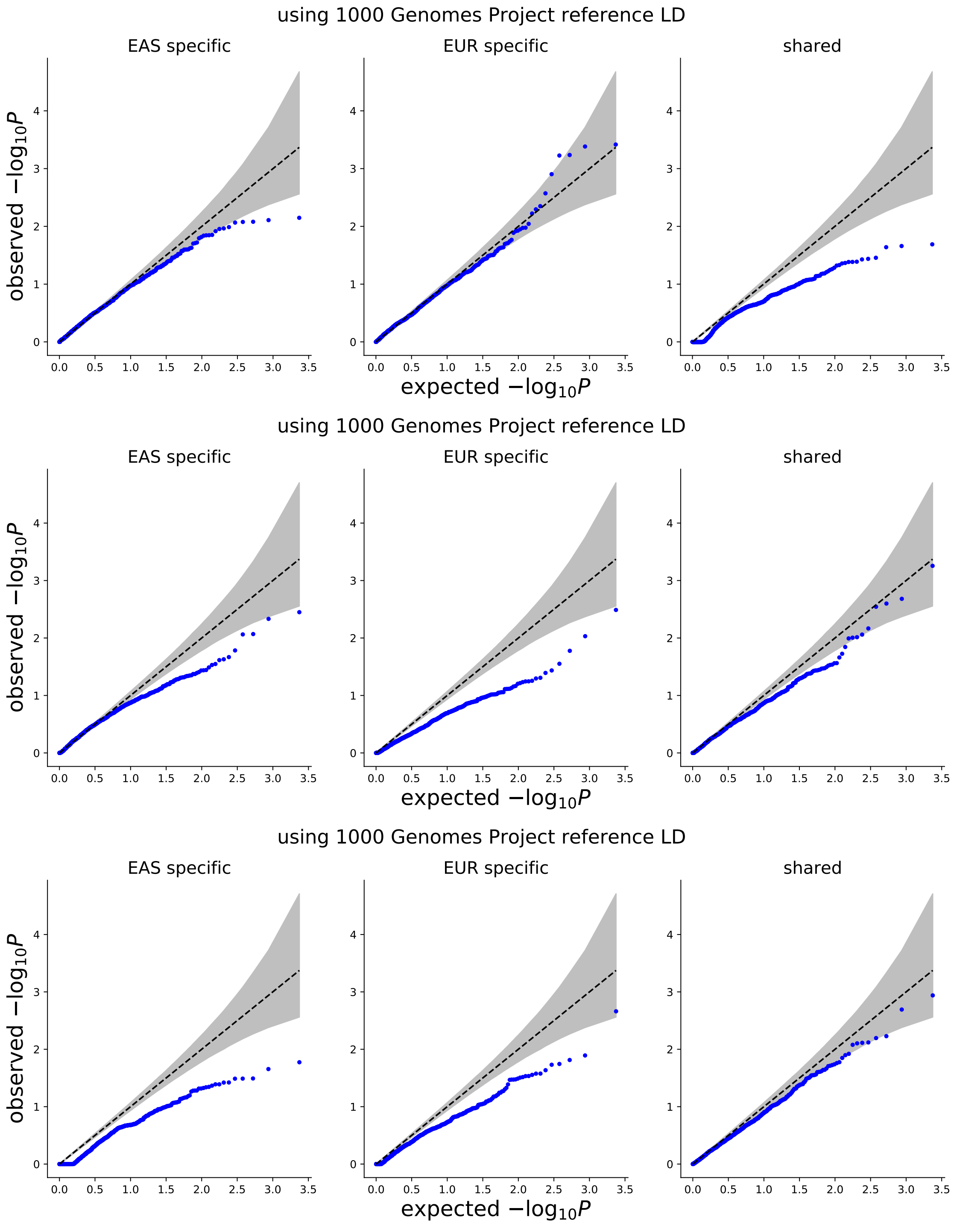

Figure S15: **Q-Q plot of p-values of the enrichments of population-specific/shared causal variants in SEG annotations**[1] **($N \times h_g^2 = 375$, 60 causal variants per population).** We computed p-values for SEG annotations across 53 GTEx tissues from 25 null simulations, where we drew 60 causal variants at random for each population. In all simulations, we set $N \times h_g^2 = 375$ in both populations. The top, middle, and bottom rows represent results from simulations where 0% (left), 50% (middle), and 100% (right) of the causal SNPs were shared. All results were obtained using 1000 Genomes Project reference LD.

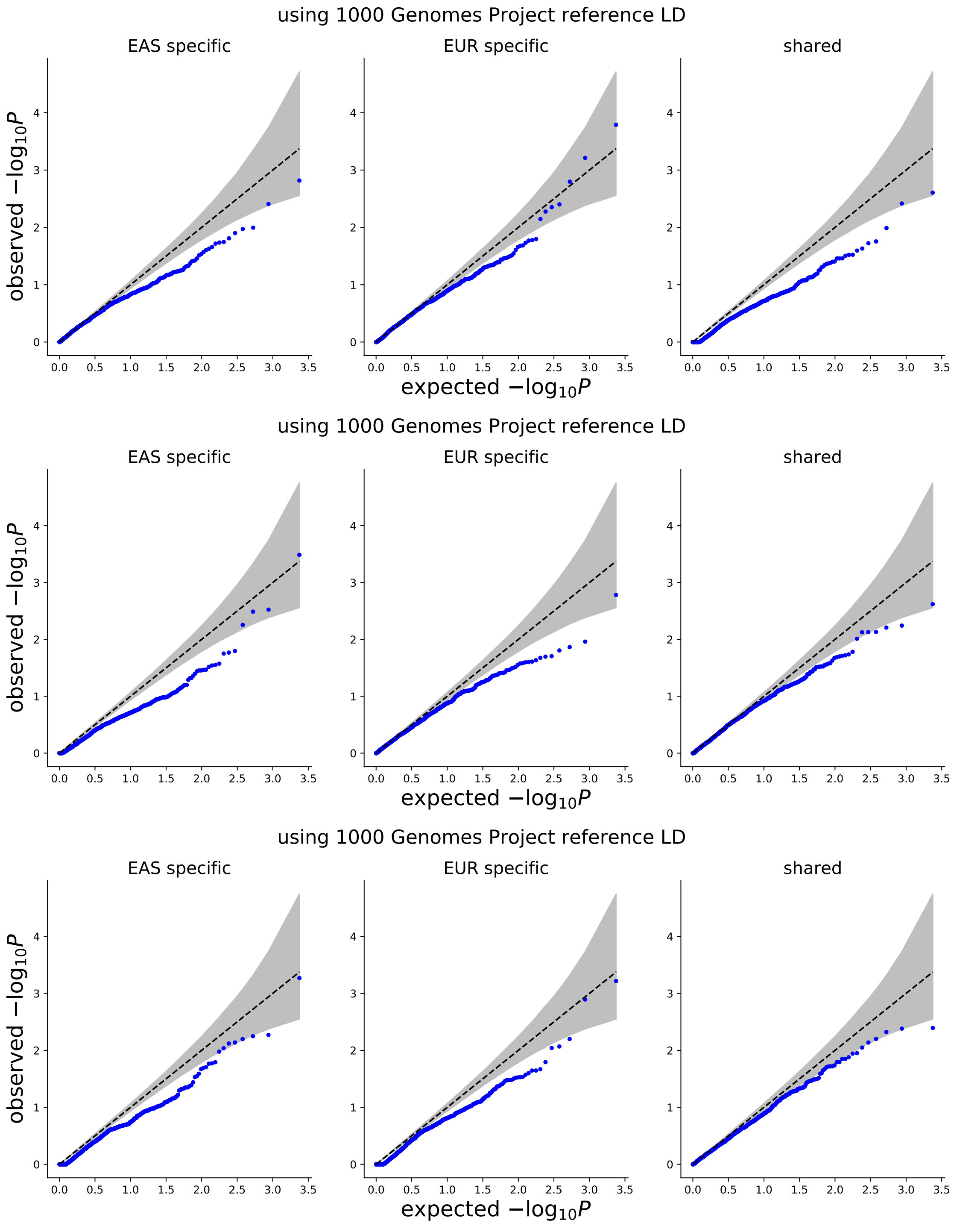

Figure S16: **Q-Q plot of p-values of the enrichments of population-specific/shared causal variants in SEG annotations**[1] **($N \times h_g^2 = 250$, 60 causal variants per population)** We computed p-values for SEG annotations across 53 GTEx tissues from 25 null simulations, where we drew 60 causal variants at random for each population. In all simulations, we set $N \times h_g^2 = 250$ in both populations. The top, middle, and bottom rows represent results from simulations where 0% (left), 50% (middle), and 100% (right) of the causal SNPs were shared. All results were obtained using 1000 Genomes Project reference LD.

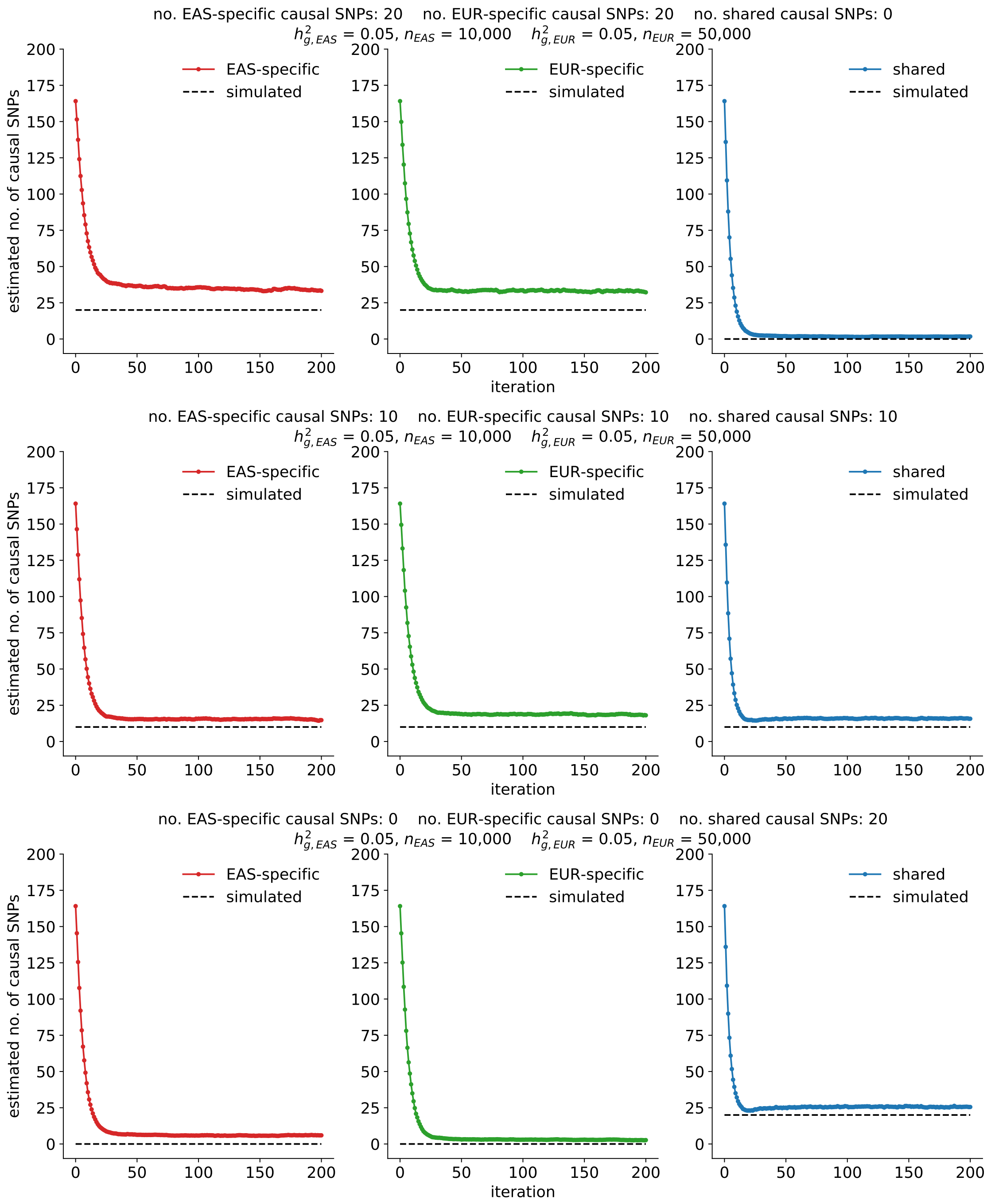

Figure S17: **Estimated numbers of population-specific/shared causal SNPs across iterations of the EM algorithm (20 causal SNPs per population).** We randomly selected 20 causal SNPs on chr22 (out of 8,599) in both populations where either 0% (top), 50% (middle) or 100% (bottom) were shared causal SNPs. $N \times h_g^2 = 500$ for both populations. Each curve represents the average estimate across 25 simulations.

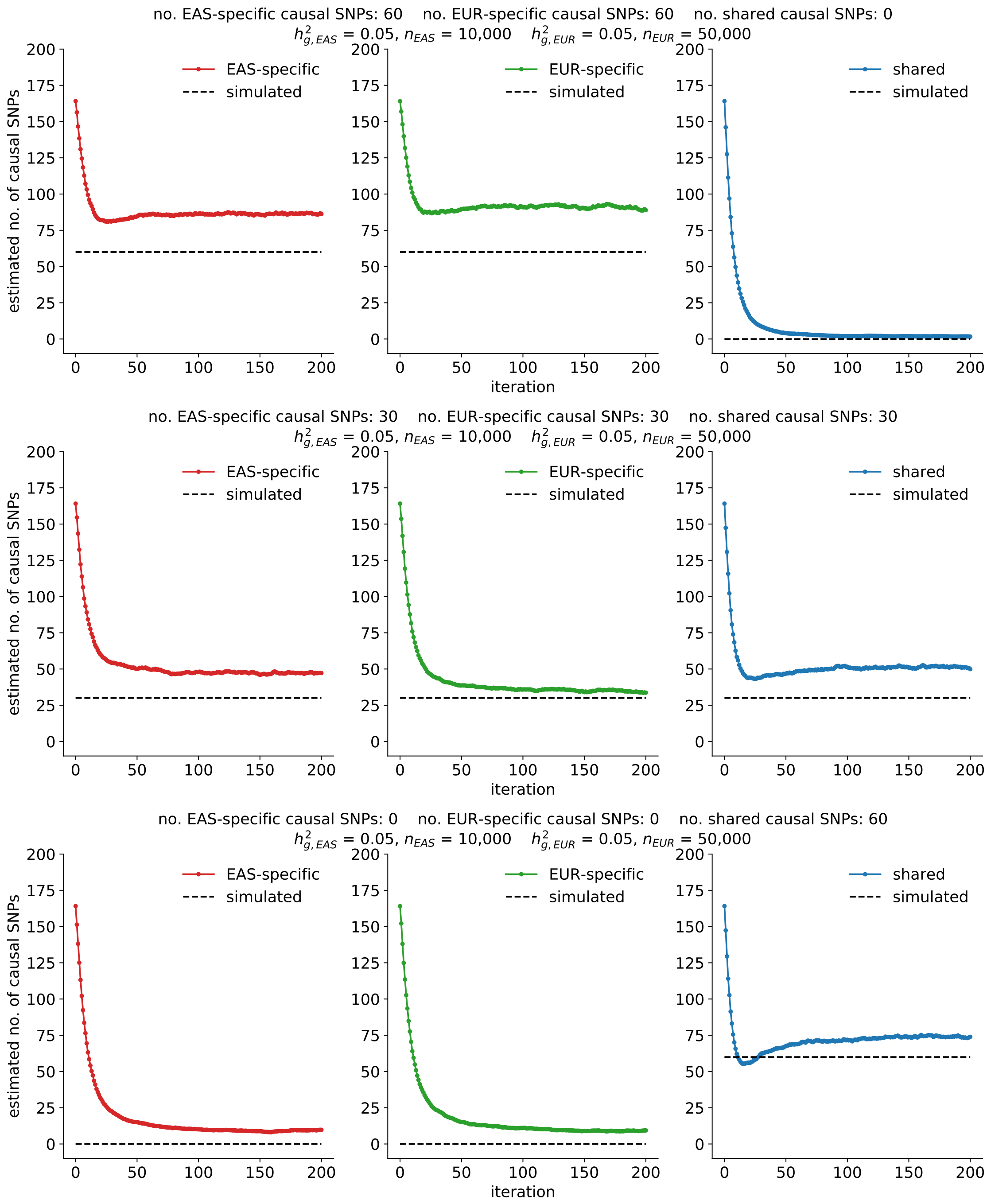

Figure S18: **Estimated number of population-specific and shared causal variants across iterations of the EM algorithm (60 causal SNPs per population).** We randomly selected 60 causal SNPs (out of 8,599) in both populations where either 0% (top), 50% (middle) or 100% (bottom) were shared causal SNPs. $N \times h_g^2 = 500$ for both populations. Each curve represents the average across 25 simulations.

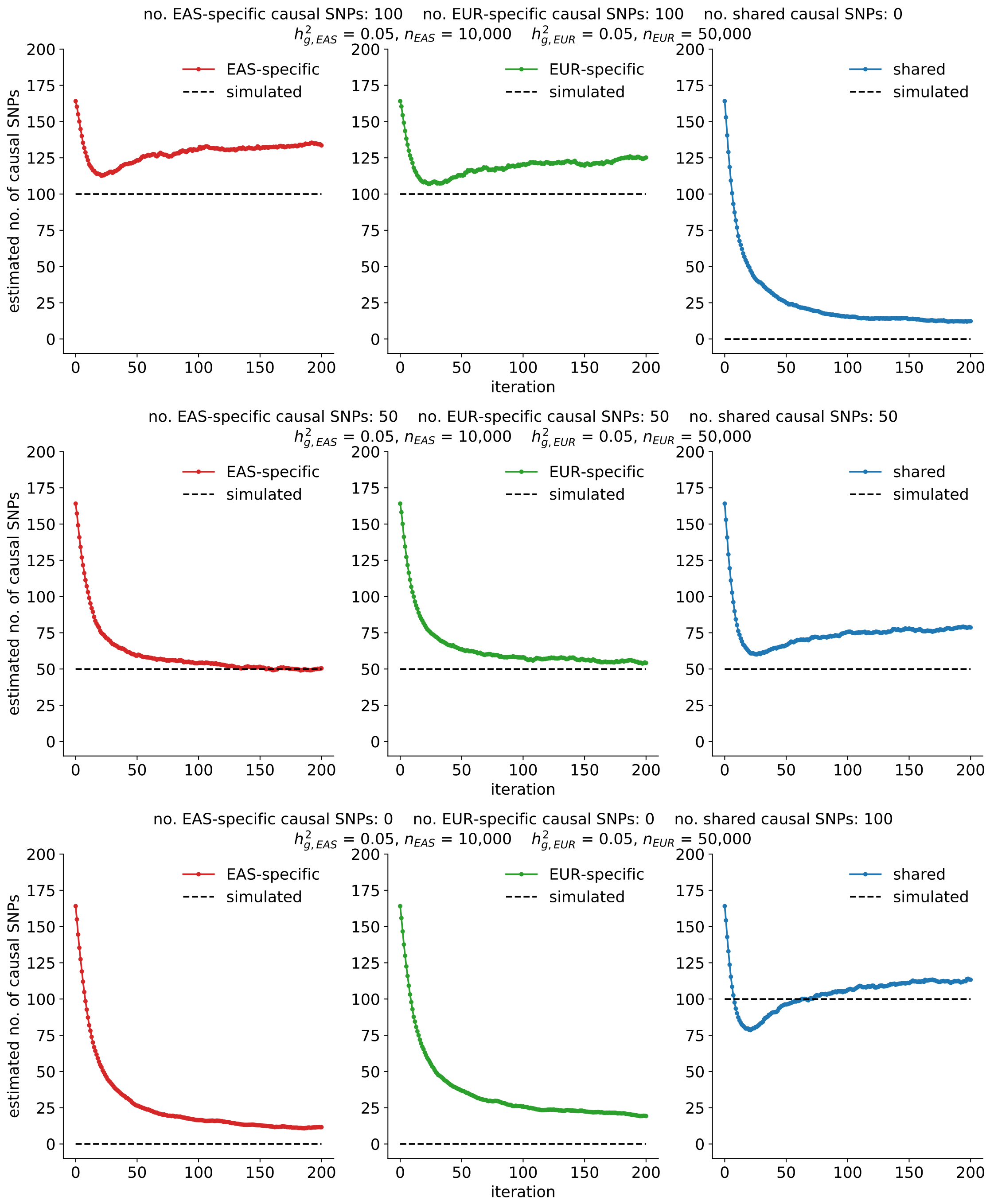

Figure S19: **Estimated number of population-specific and shared causal variants across iterations of the EM algorithm (100 causal SNPs per population).** We randomly selected 100 causal SNPs (out of 8,599) in both populations where either 0% (top), 50% (middle) or 100% (bottom) were shared causal SNPs. $N \times h_g^2 = 500$ for both populations. Each curve represents the average across 25 simulations.

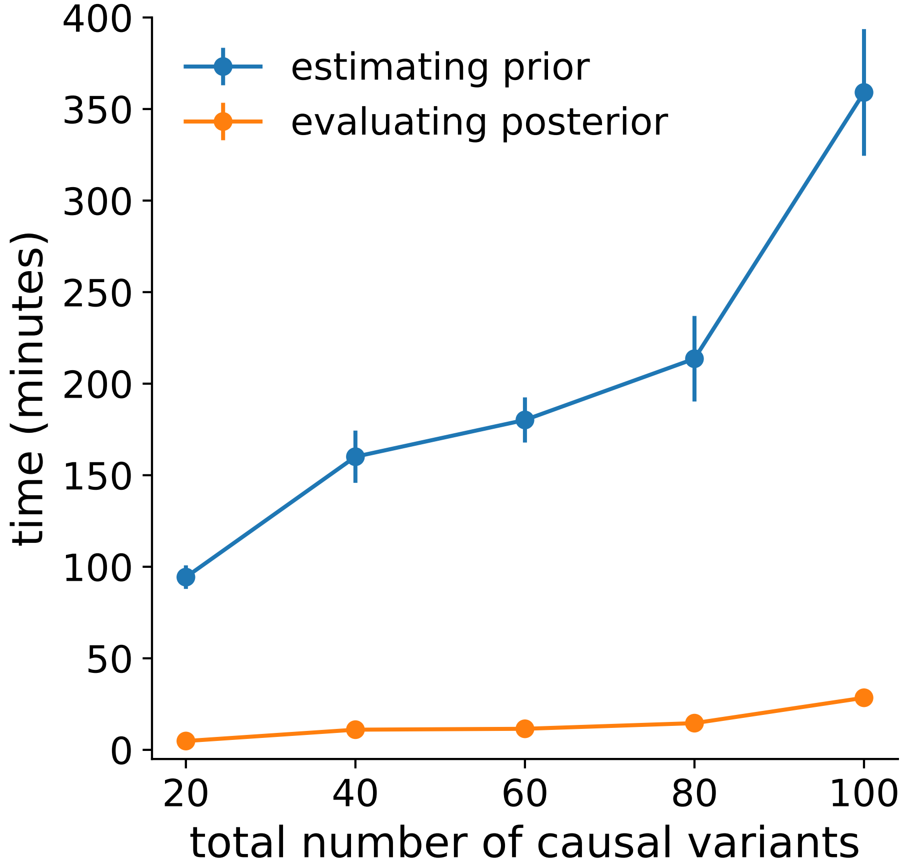

Figure S20: **Average run-times for estimating the prior (MVB parameters) and evaluating the per-SNP posterior probabilities of being causal in one or both populations.** Each dot represents the average run-time across 25 simulations; the total number of causal variants per population is specified on the x-axis. $N \times h_g^2 = 500$ for both populations. Error bars represent $\pm$1.96 s.e.m.

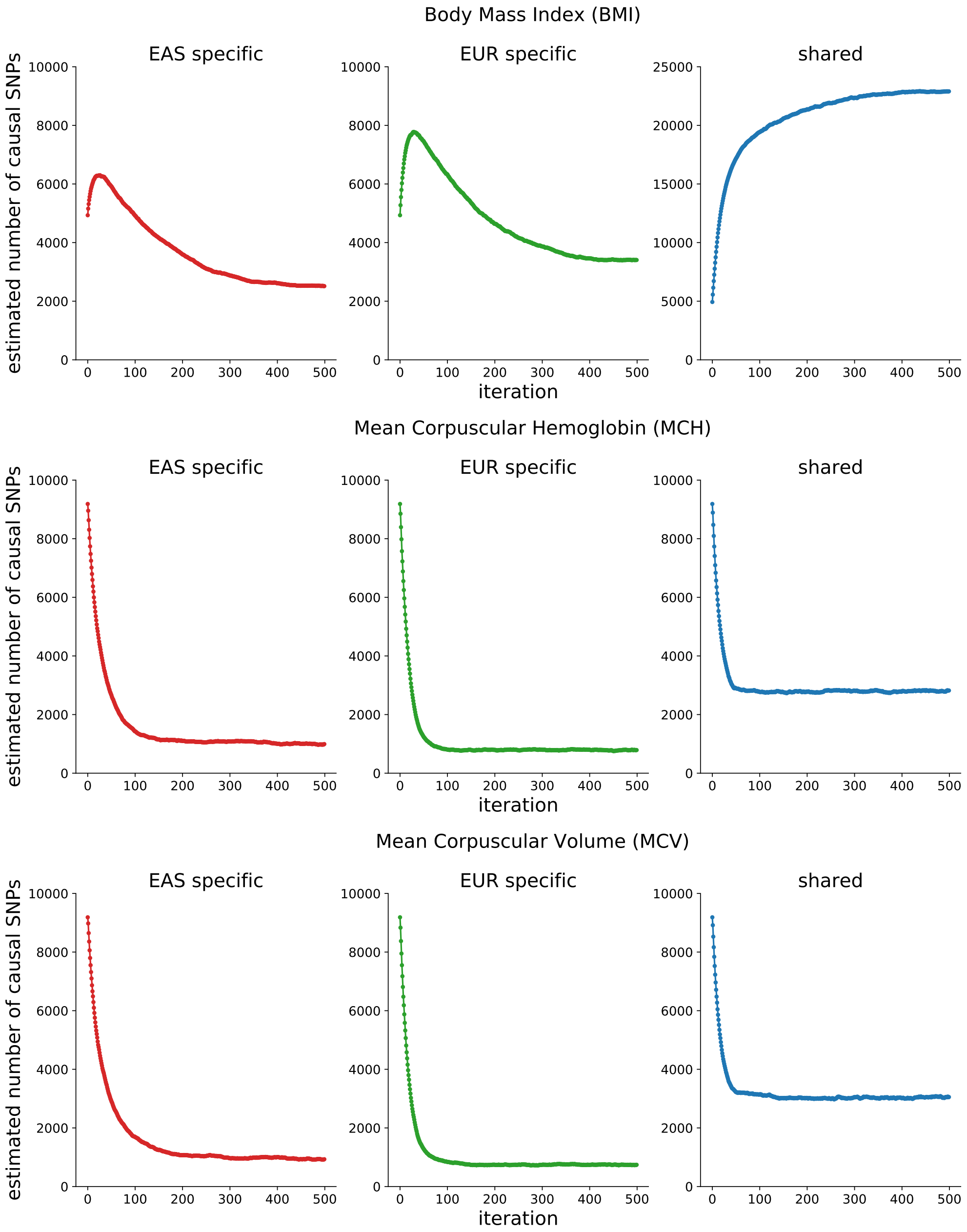

Figure S21: **Estimated numbers of population-specific/shared causal variants across EM iterations for BMI, MCH, and MCV.**

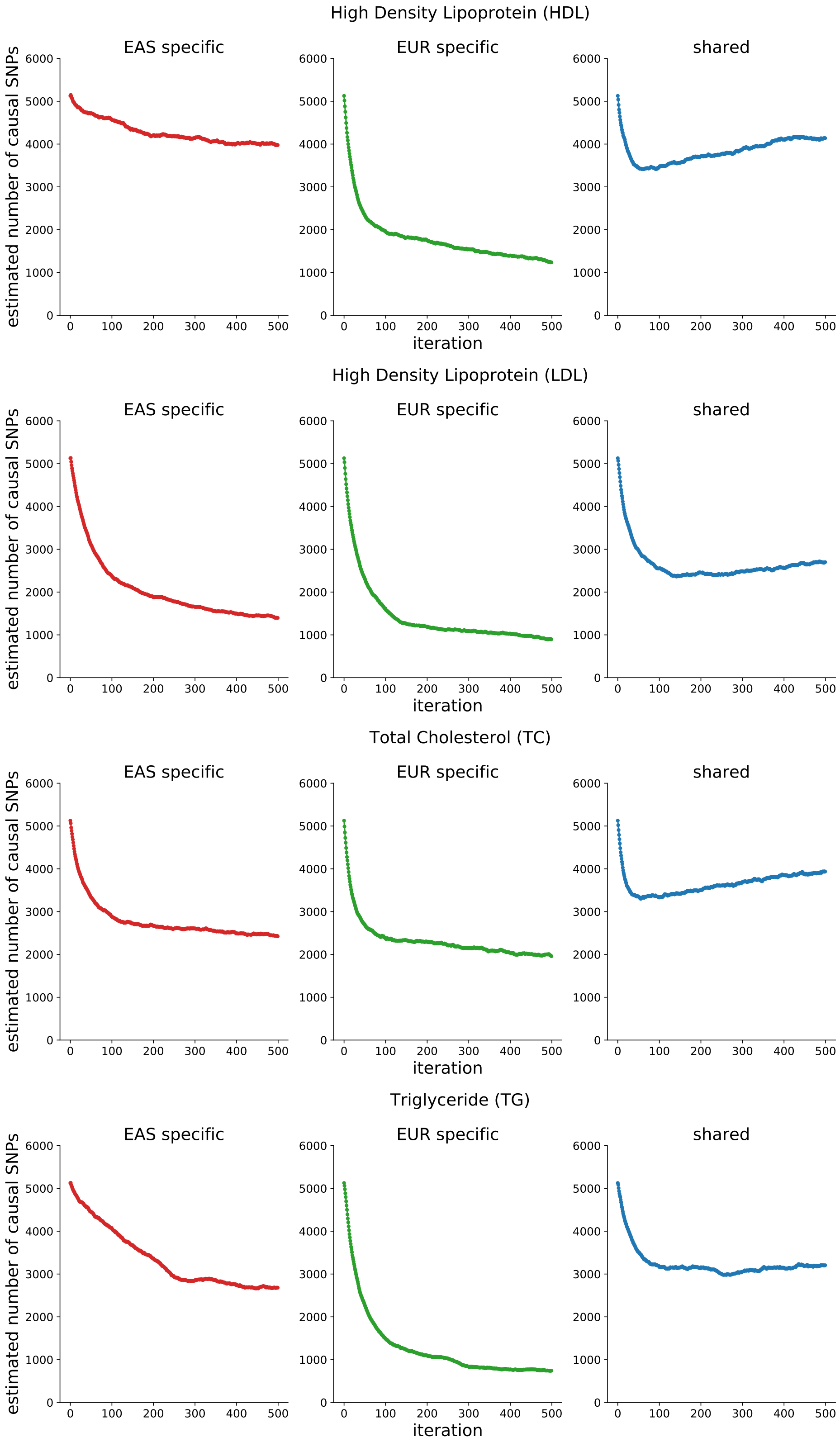

Figure S22: **Estimated numbers of population-specific/shared causal variants across EM iterations for HDL, LDL, TC, and TG.**

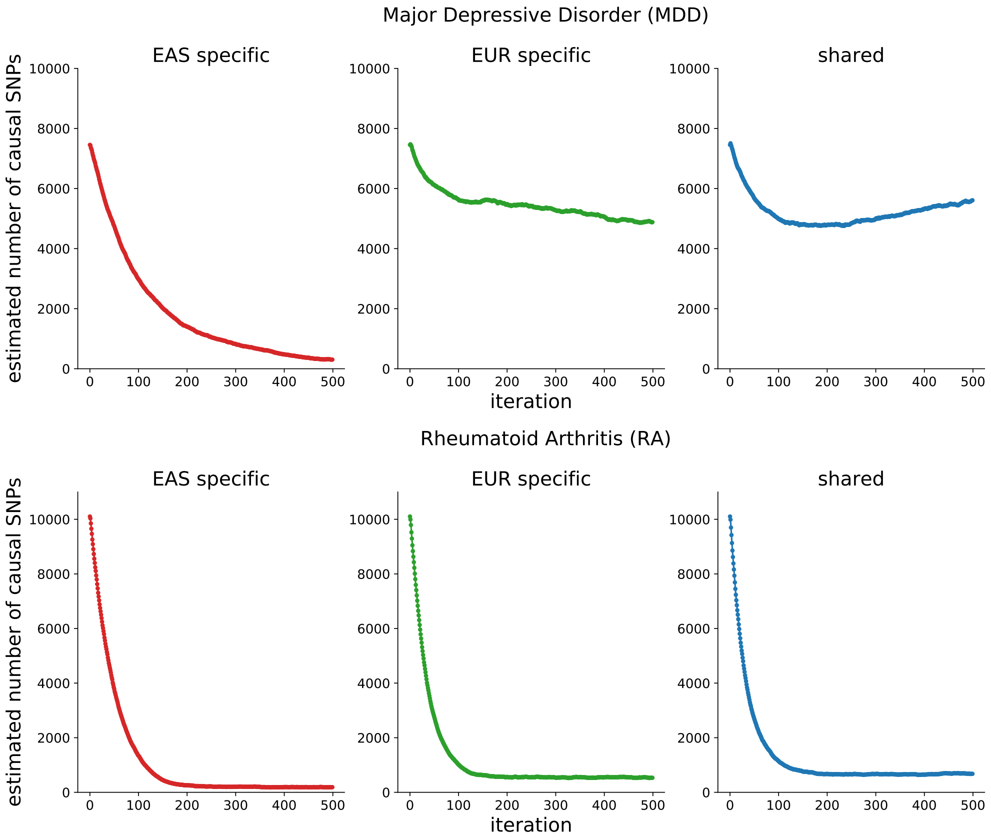

Figure S23: **Estimated numbers of population-specific/shared causal variants across EM iterations for MDD and RA.**

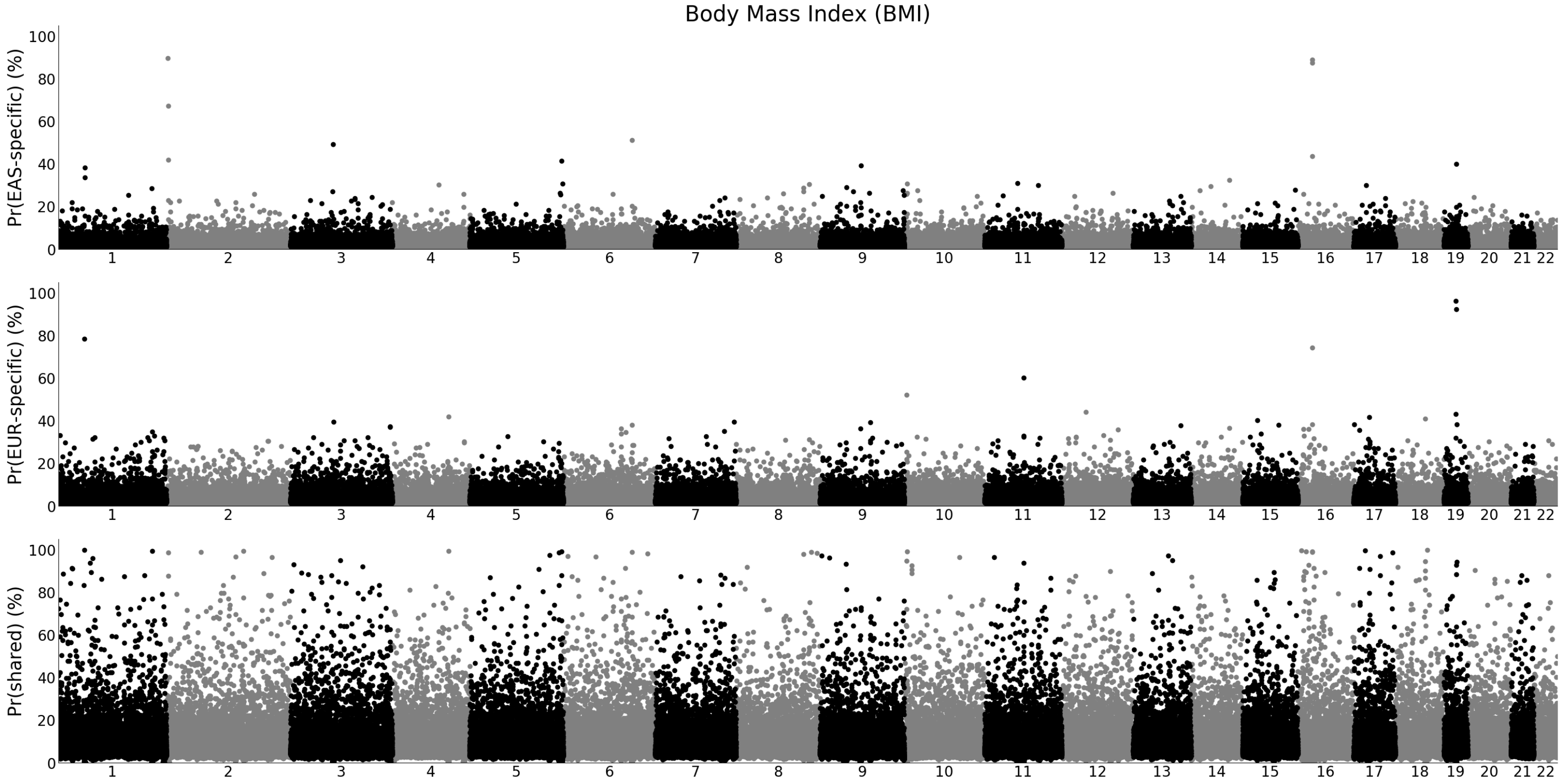

Figure S24: **Manhattan-style plots for posterior probability of each SNP to population-specific or shared for BMI.**

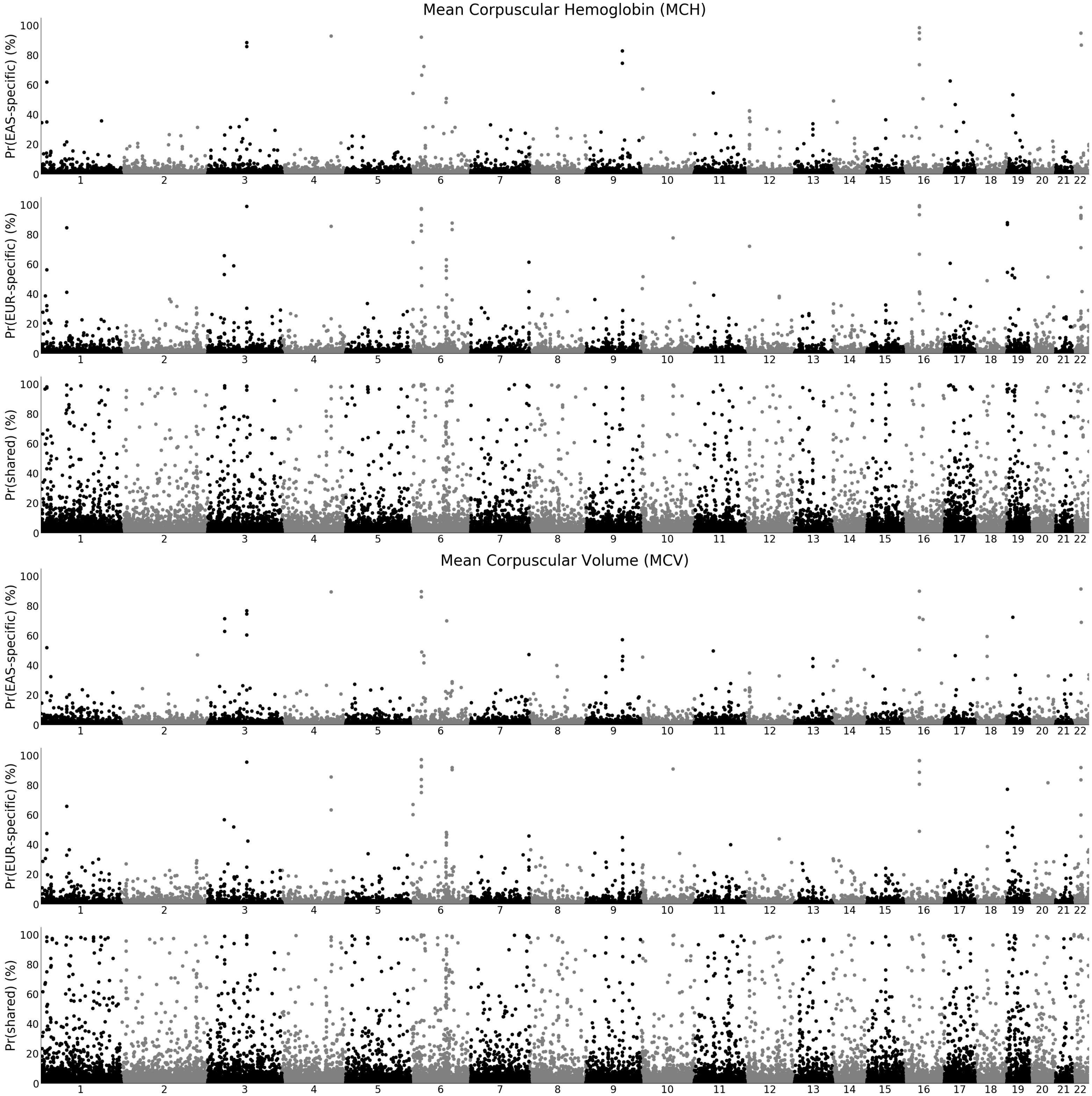

Figure S25: **Manhattan-style plots for posterior probability of each SNP to population-specific or shared for MCH and MCV.**

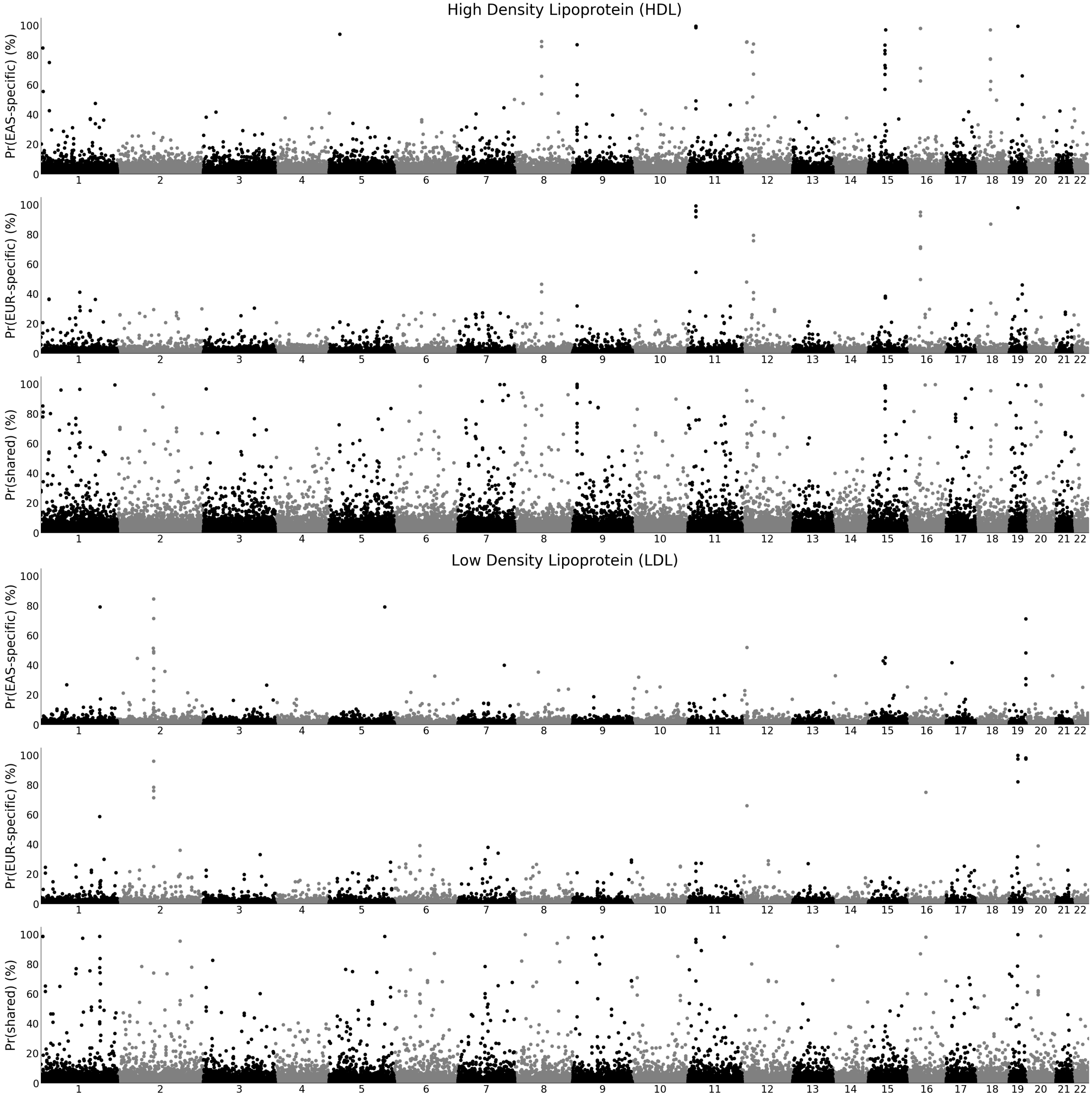

Figure S26: **Manhattan-style plots for posterior probability of each SNP to population-specific or shared for HDL and LDL.**

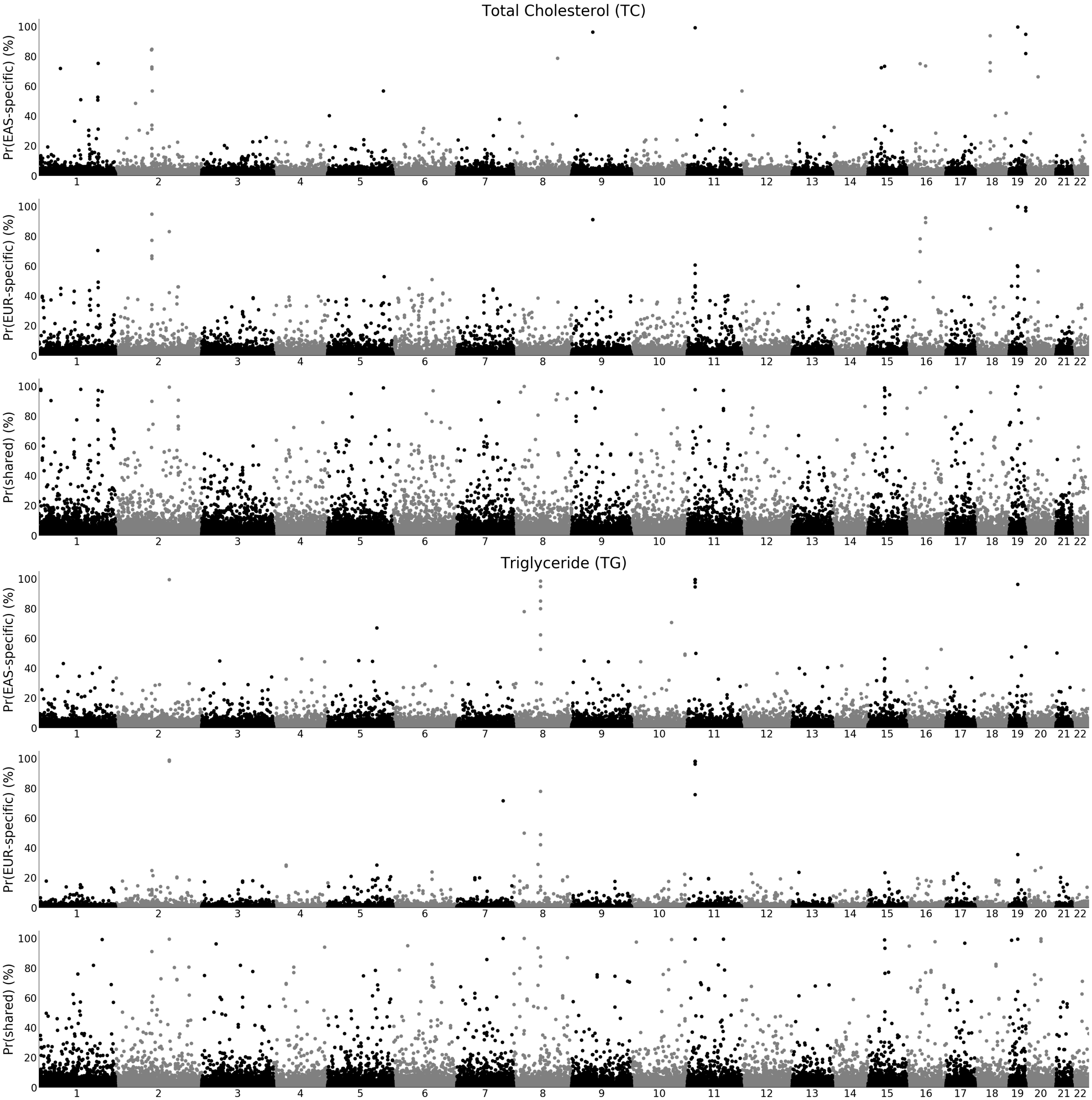

Figure S27: **Manhattan-style plots for posterior probability of each SNP to population-specific or shared for TC and TG.**

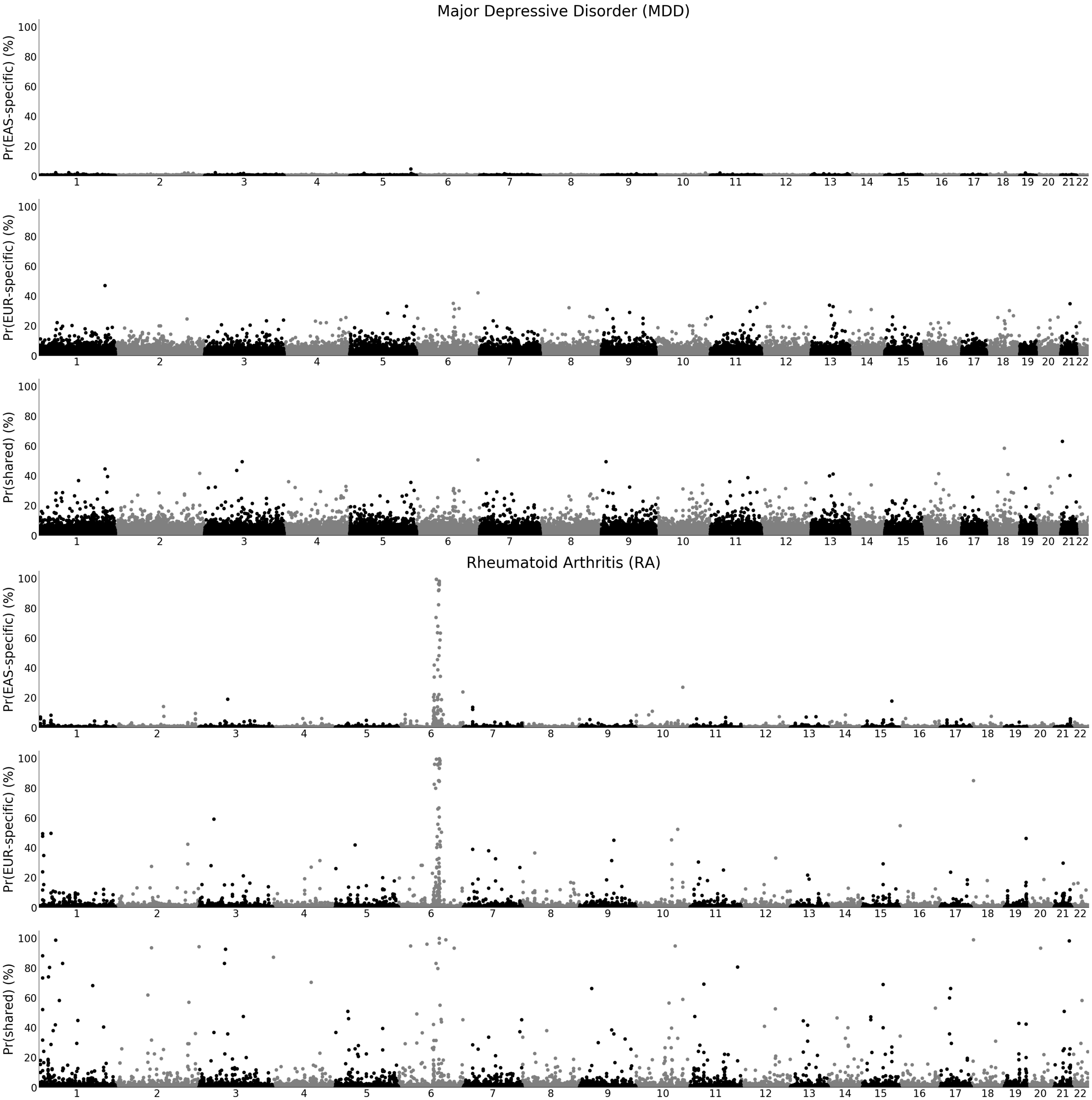

Figure S28: **Manhattan-style plots for posterior probability of each SNP to population-specific or shared for MDD and RA.**

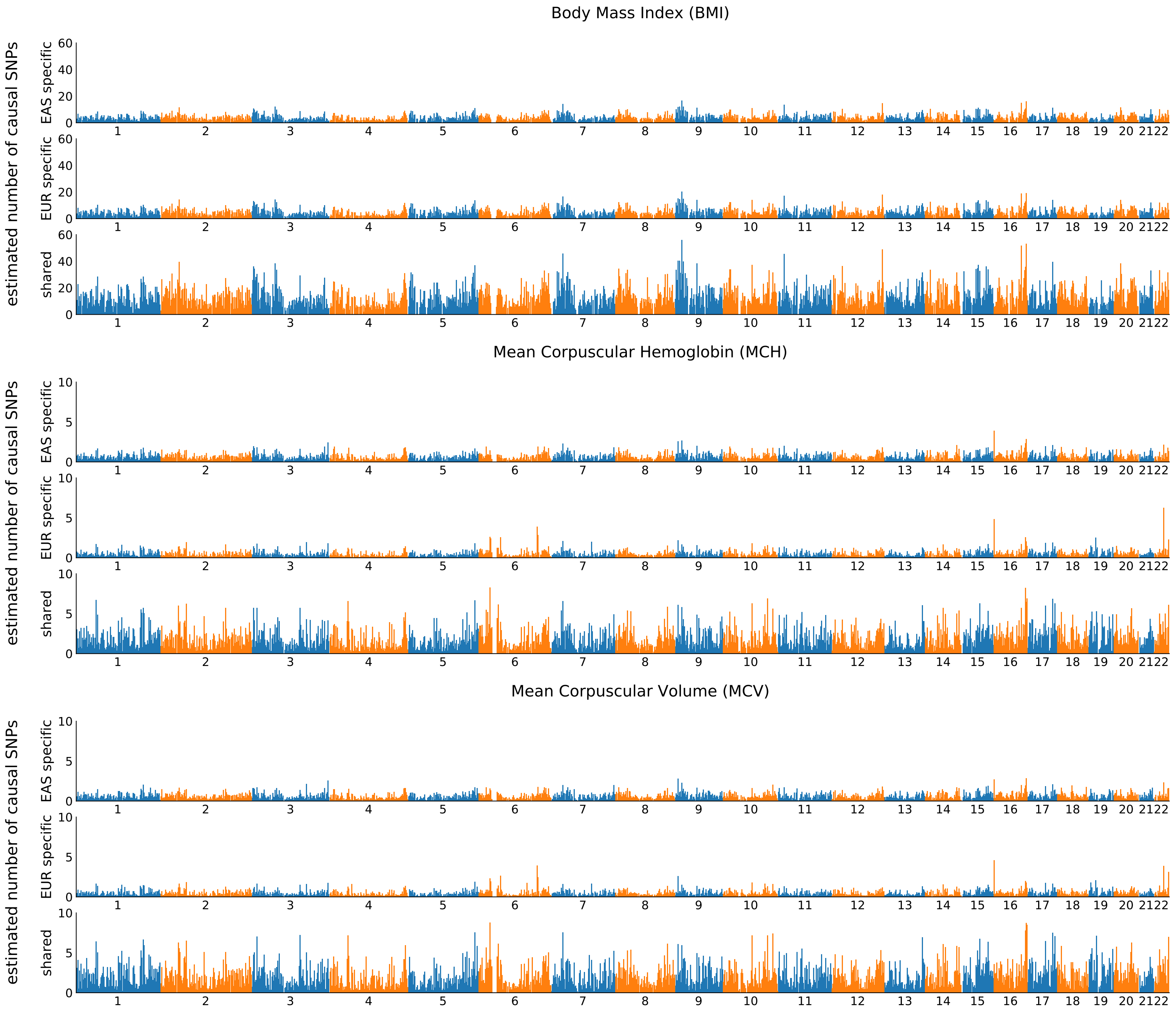

Figure S29: **Regional number of causal variants for BMI, MCH, and MCV.**

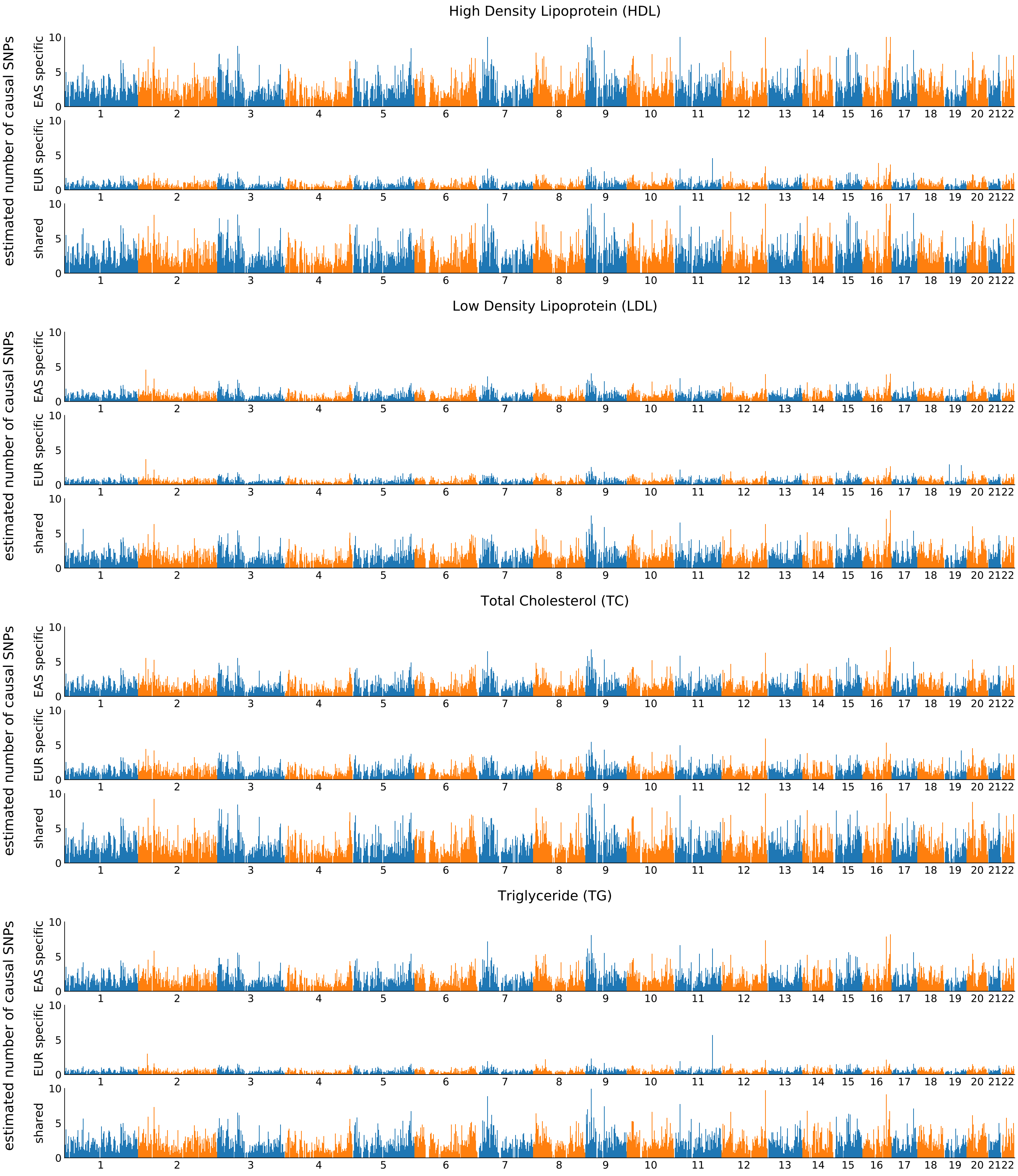

Figure S30: **Regional number of causal variants for HDL, LDL, TC, and TG.**

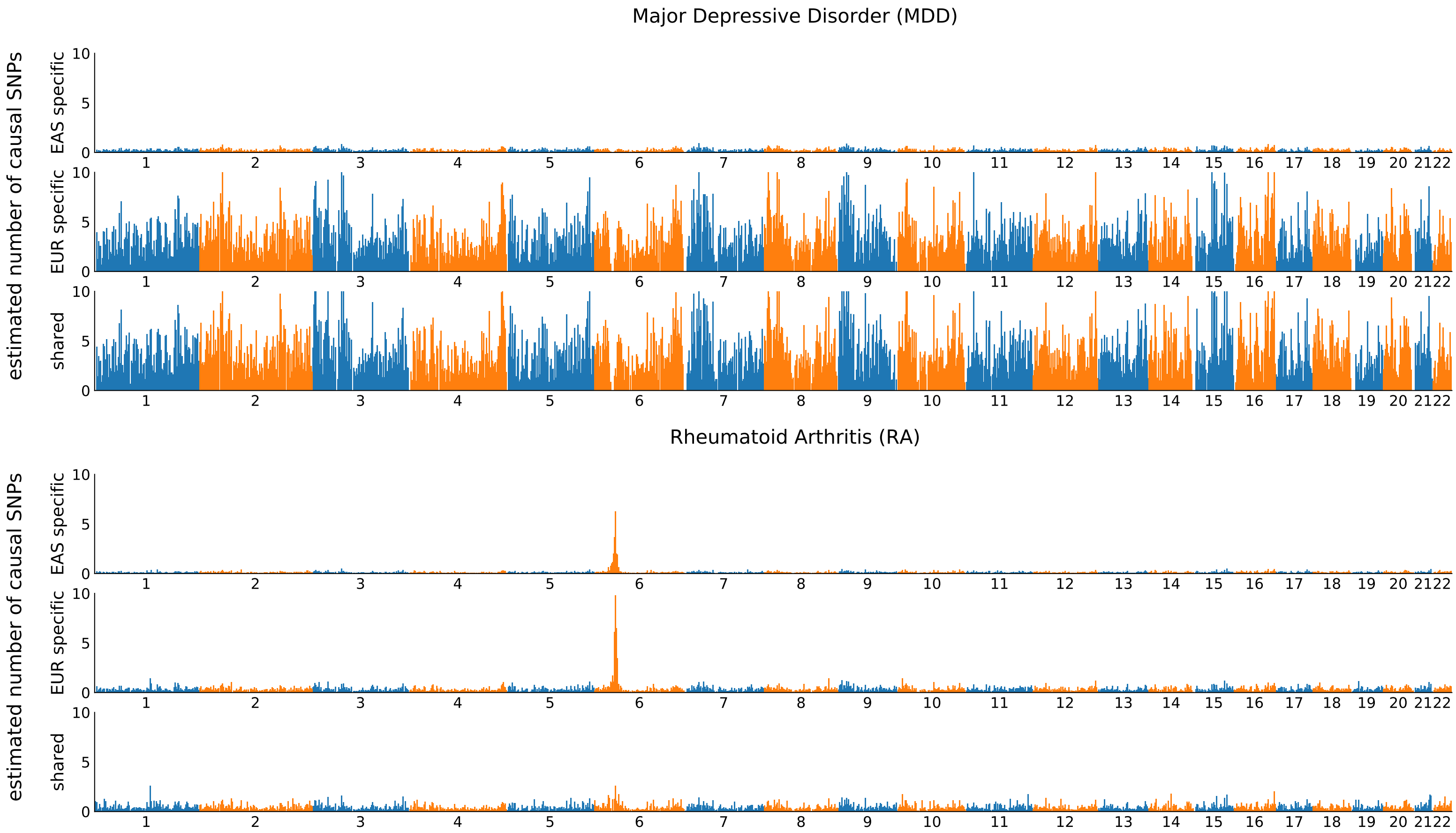

Figure S31: **Regional number of causal variants for MDD and RA.**

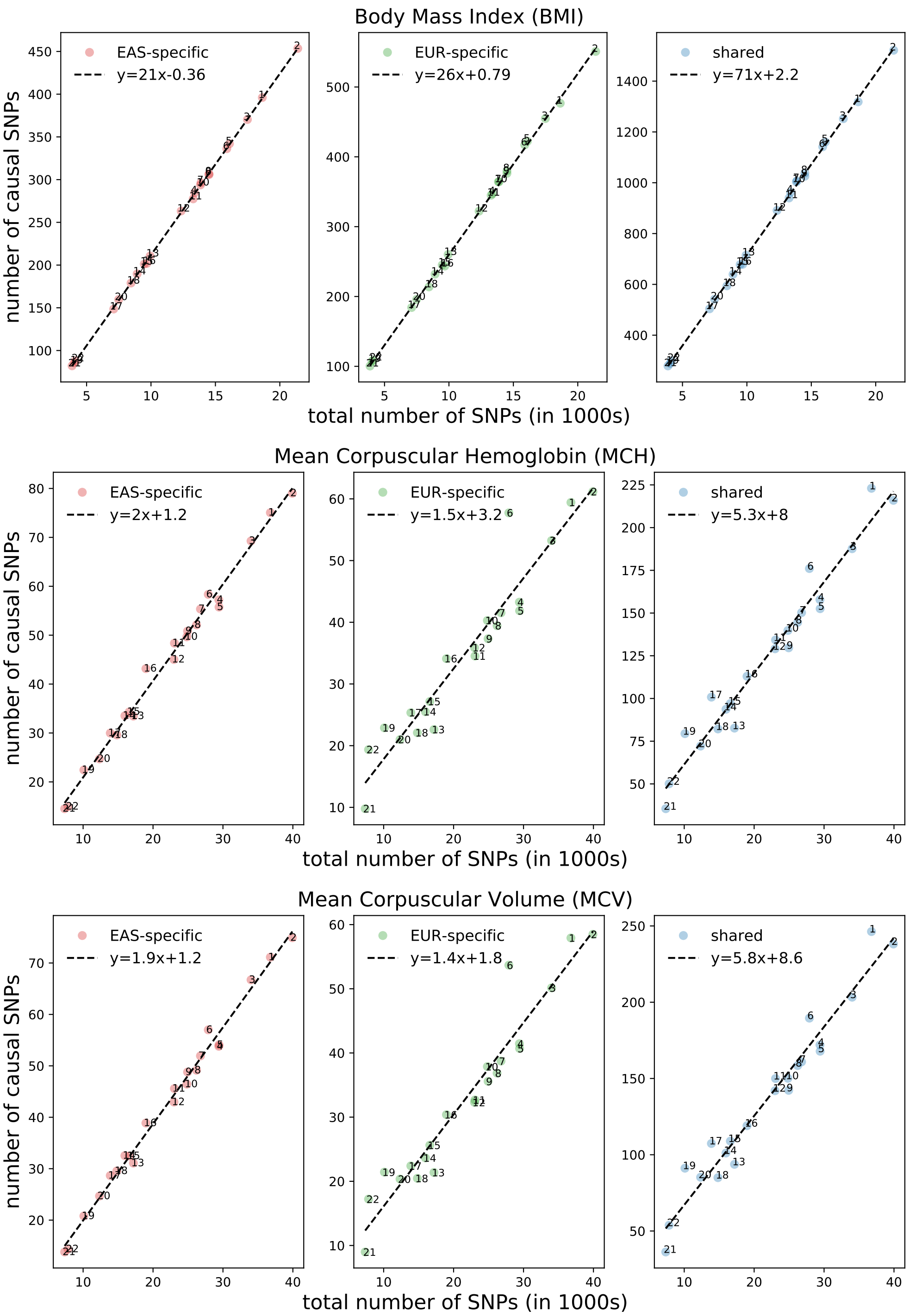

Figure S32: **Chromosomal number of causal variants for BMI, MCH, and MCV.**

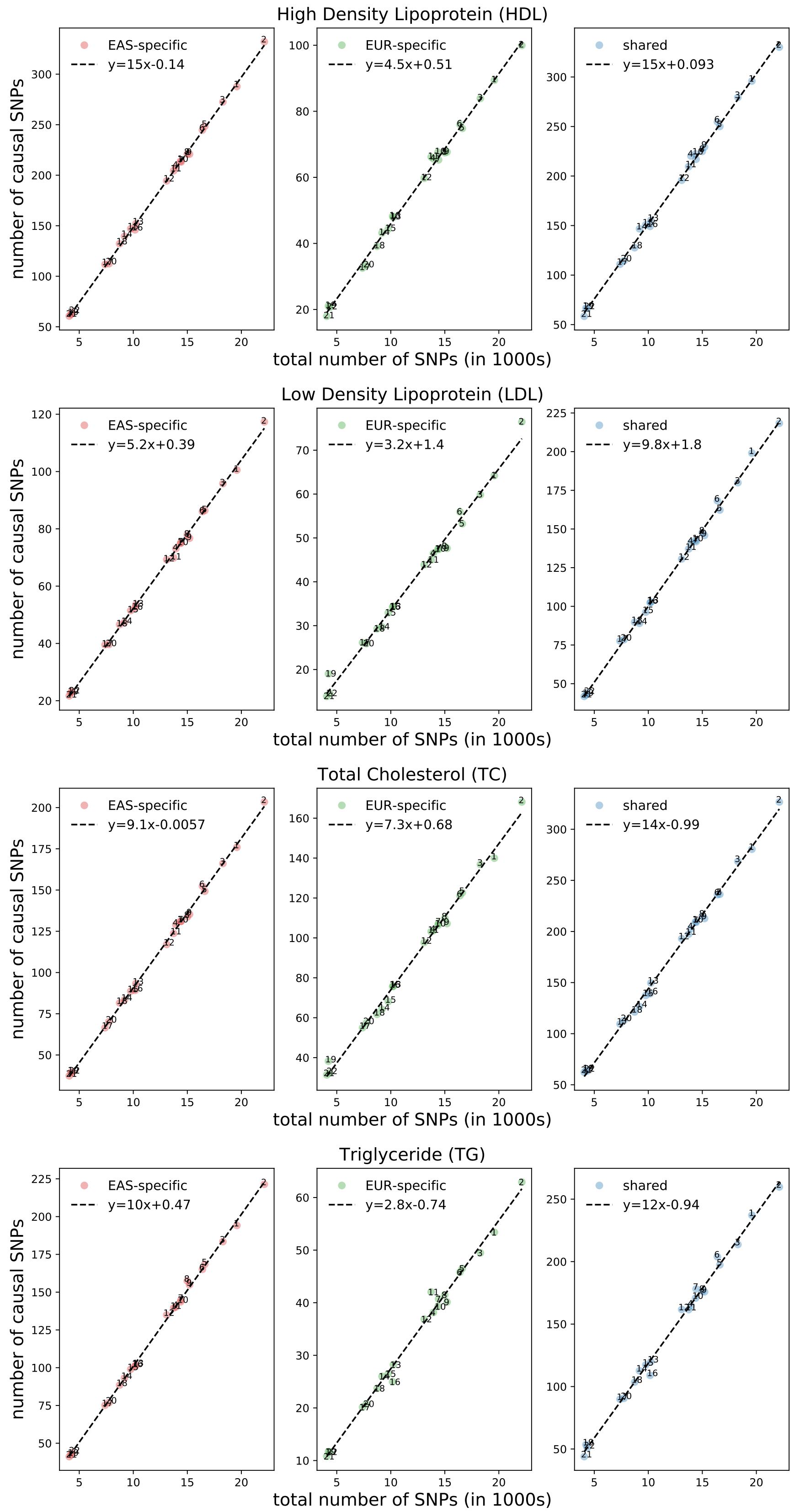

Figure S33: **Chromosomal number of causal variants for HDL, LDL, TC, and TG.**

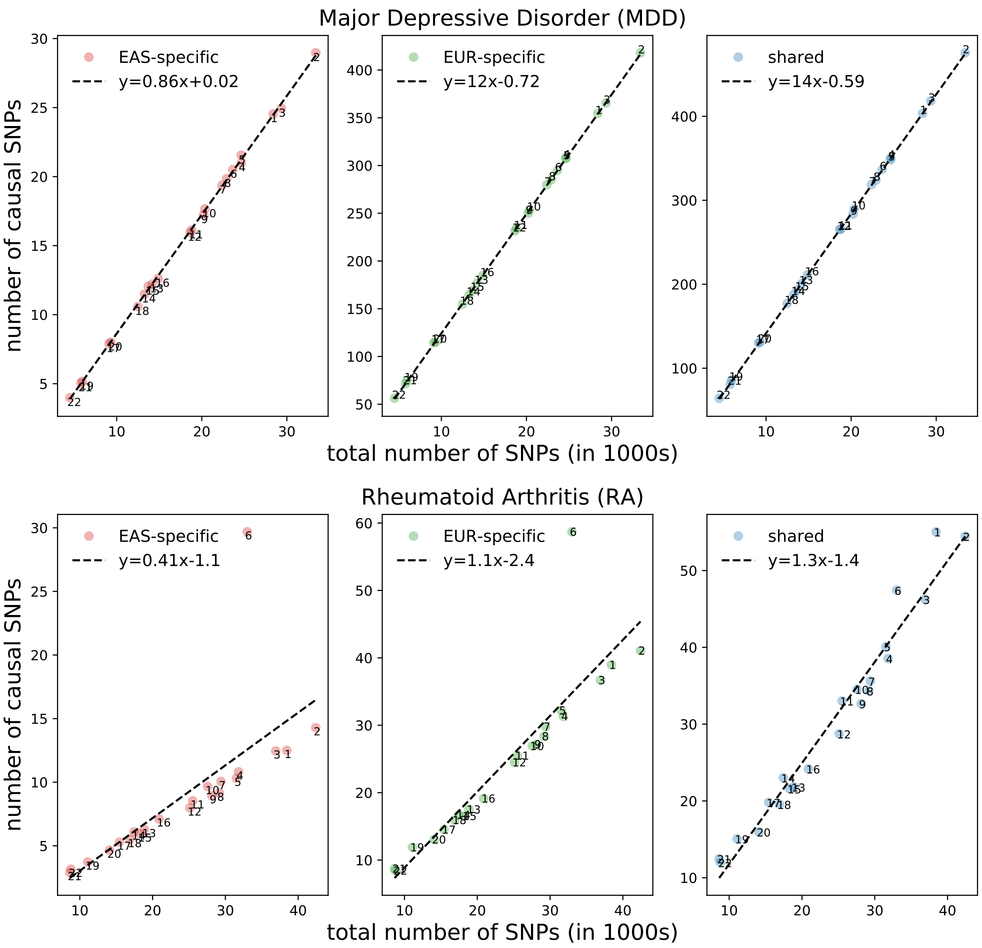

Figure S34: **Chromosomal number of causal variants for MDD and RA.**

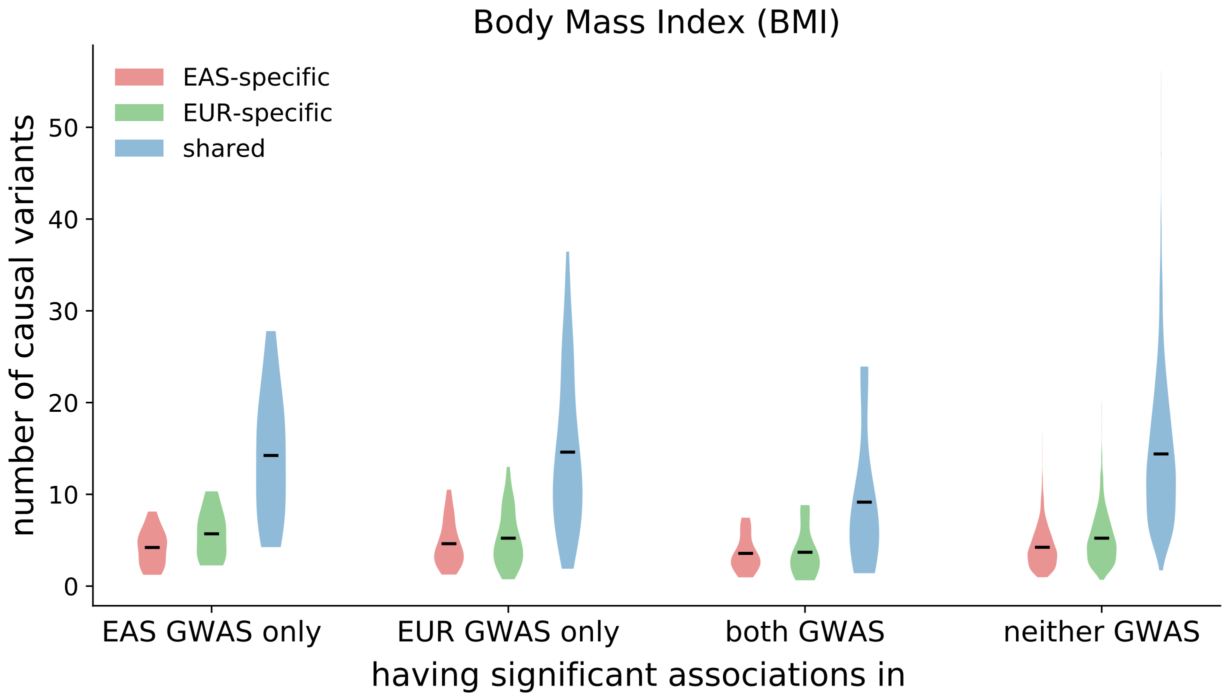

Figure S35: **Distribution of regional number of causal variants at GWAS risk regions.** Each violin plot shows the distribution of population-specific or shared causal variants at regions harboring significant associations ($p < 5 \times 10^{-5}$) in the East Asian GWAS only, in the European GWAS only, in both GWASs, and in neither GWAS. The dark line represents the mean of the distribution.

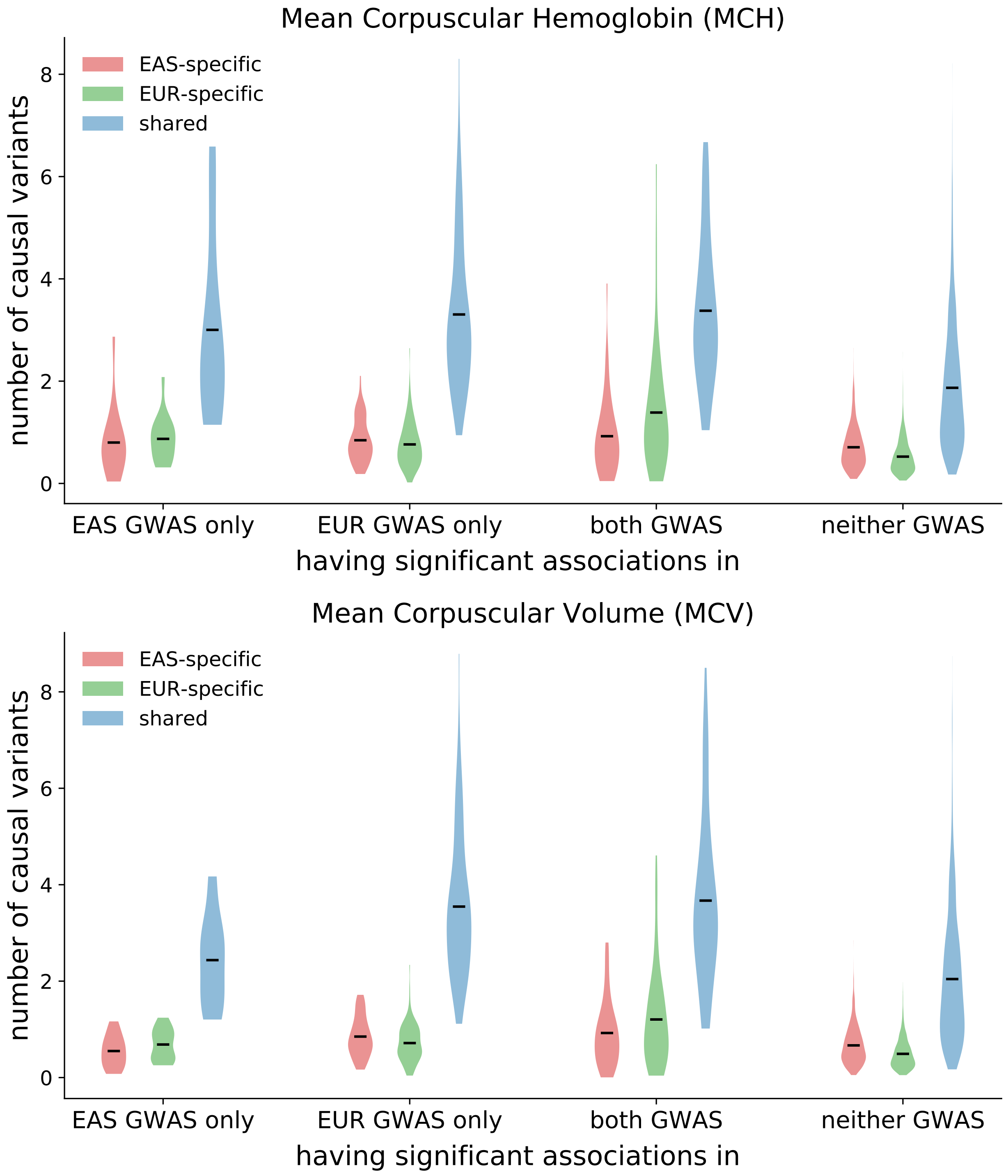

Figure S36: **Distribution of regional number of causal variants at GWAS risk regions.** Each violin plot shows the distribution of population-specific or shared causal variants at regions harboring significant associations ($p < 5 \times 10^{-5}$) in the East Asian GWAS only, in the European GWAS only, in both GWASs, and in neither GWAS. The dark line represents the mean of the distribution.

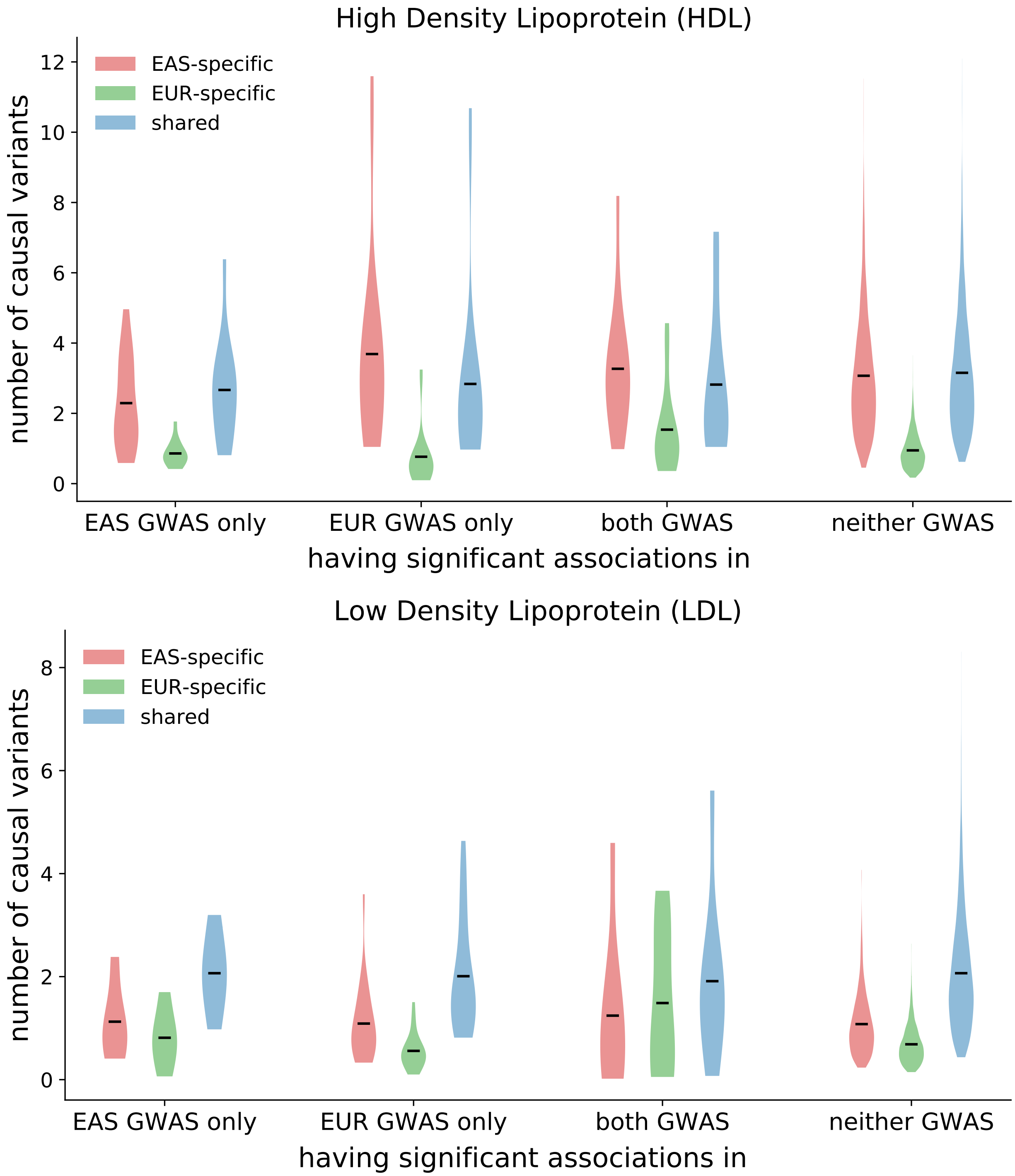

Figure S37: **Distribution of regional number of causal variants at GWAS risk regions.** Each violin plot shows the distribution of population-specific or shared causal variants at regions harboring significant associations ($p < 5 \times 10^{-5}$) in the East Asian GWAS only, in the European GWAS only, in both GWASs, and in neither GWAS. The dark line represents the mean of the distribution.

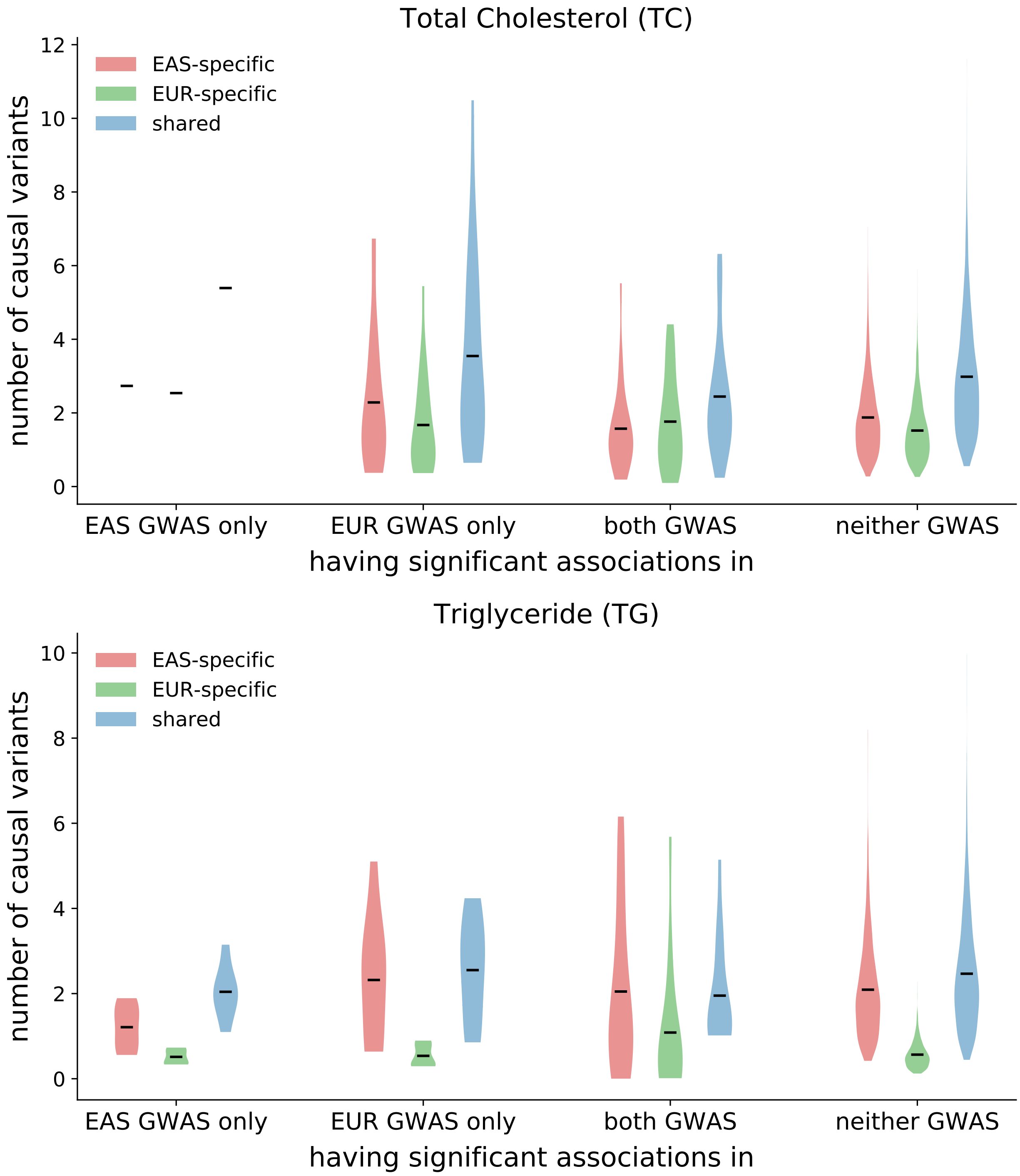

Figure S38: **Distribution of regional number of causal variants at GWAS risk regions.** Each violin plot shows the distribution of population-specific or shared causal variants at regions harboring significant associations ($p < 5 \times 10^{-5}$) in the East Asian GWAS only, in the European GWAS only, in both GWASs, and in neither GWAS. The dark line represents the mean of the distribution.

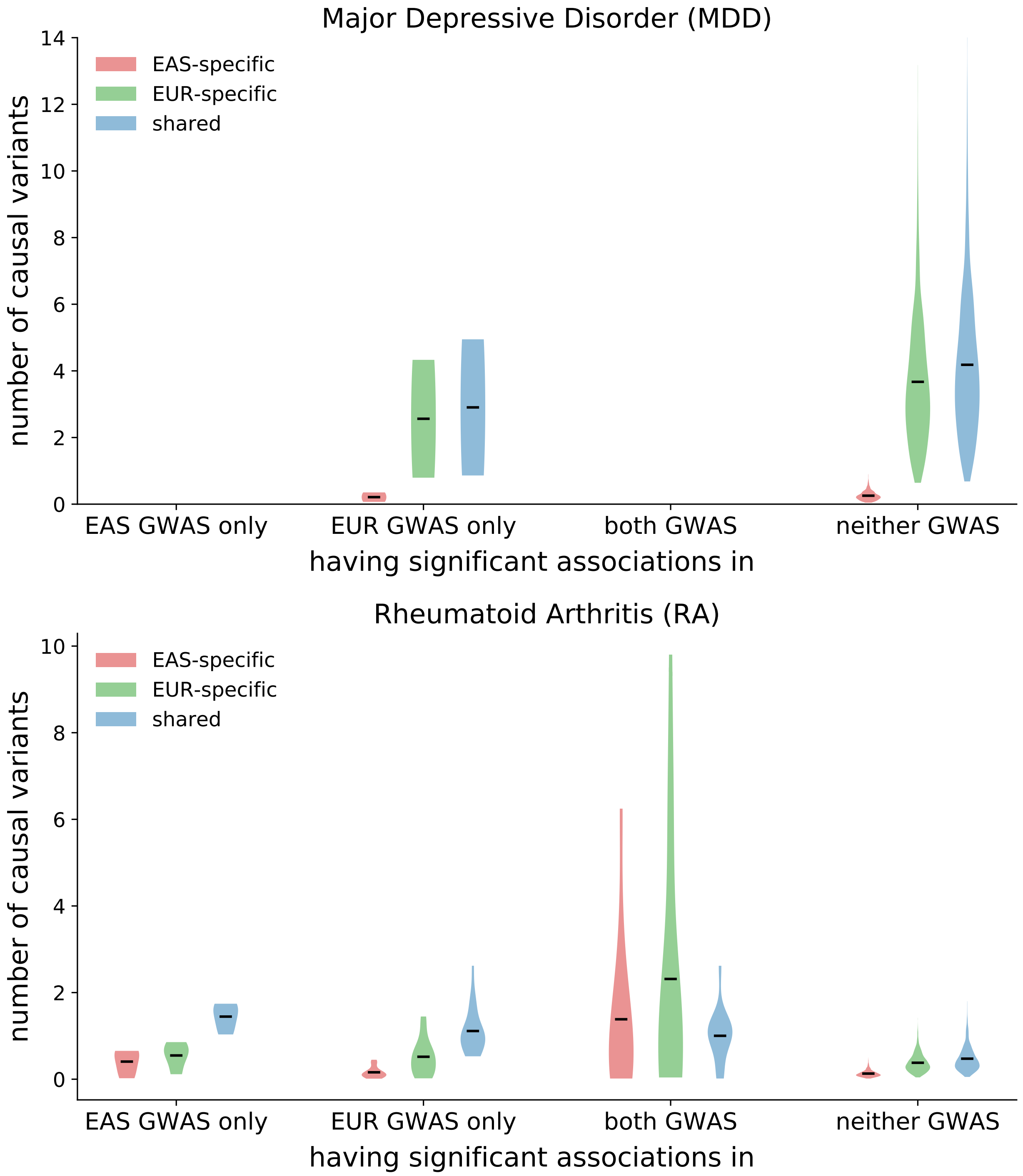

Figure S39: **Distribution of regional number of causal variants at GWAS risk regions.** Each violin plot shows the distribution of population-specific or shared causal variants at regions harboring significant associations ($p < 5 \times 10^{-5}$) in the East Asian GWAS only, in the European GWAS only, in both GWASs, and in neither GWAS. The dark line represents the mean of the distribution.

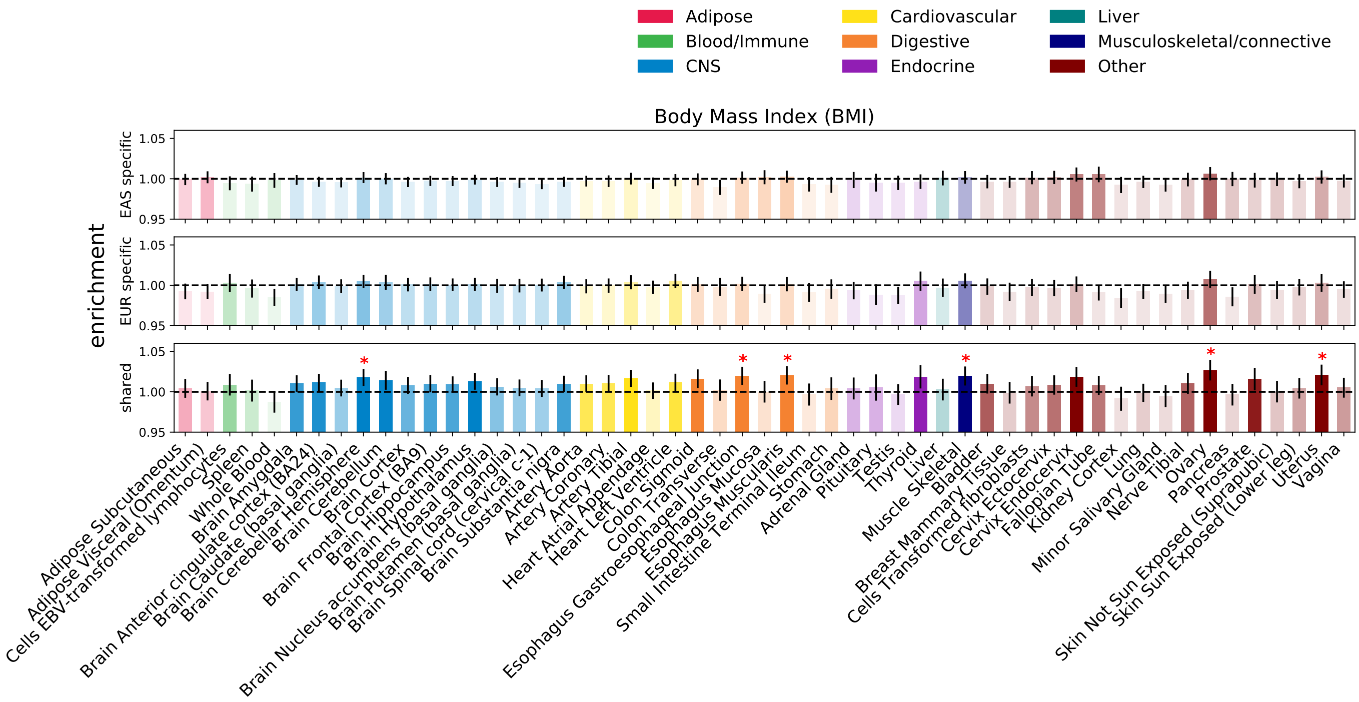

Figure S40: **Enrichment of population-specific and shared causal variants in specifically expressed genes annotation across 53 GTEx tissues.** Error bars represent 1.96 times the standard error on each side. The darker the color, the more significant an enrichment is. We mark enrichment with p-value less than $0.05/53$ with a star.

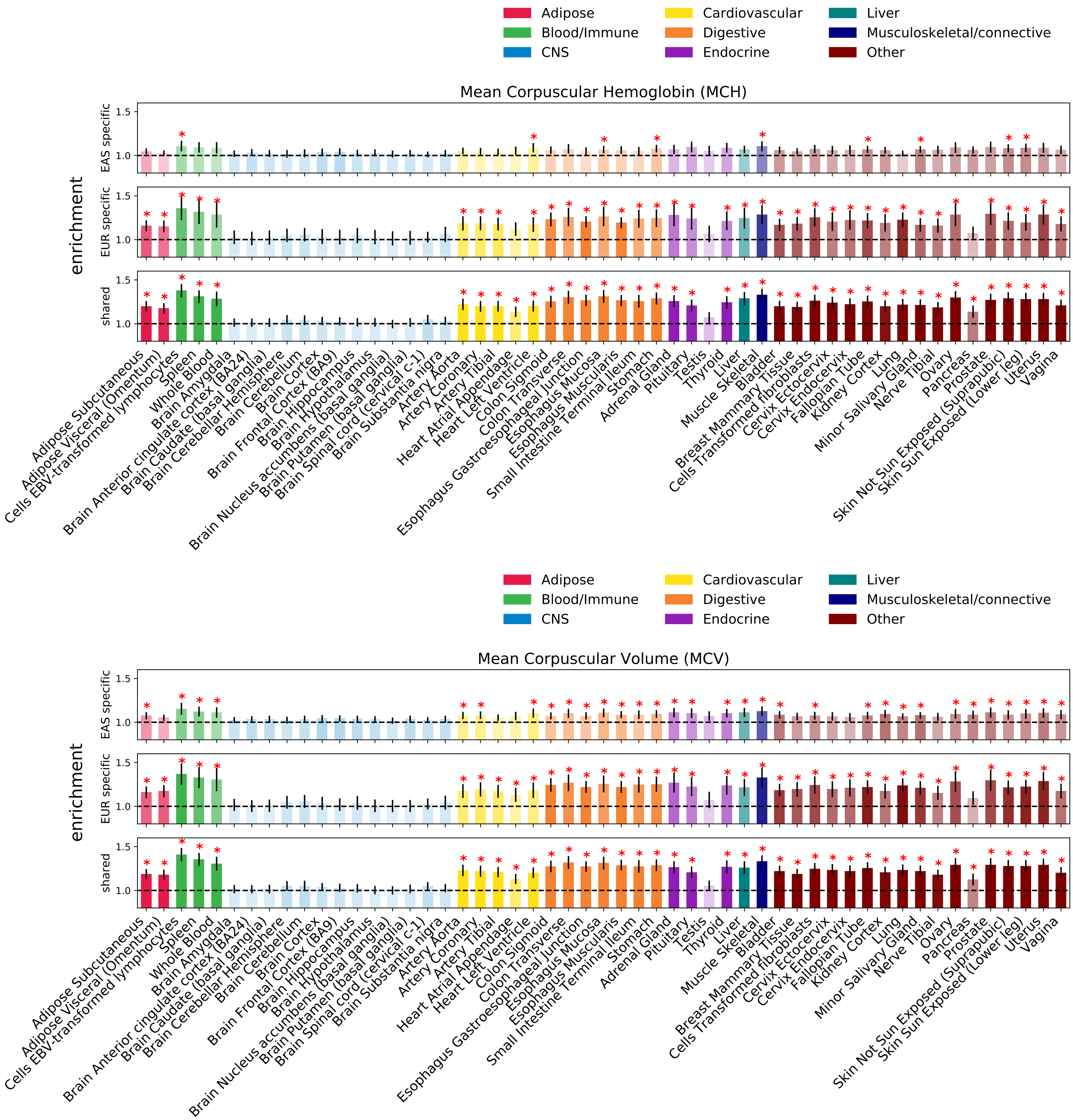

Figure S41: **Enrichment of population-specific and shared causal variants in specifically expressed genes annotation across 53 GTEx tissues.** Error bars represent 1.96 times the standard error on each side. The darker the color, the more significant an enrichment is. We mark enrichment with p-value less than $0.05/53$ with a star.

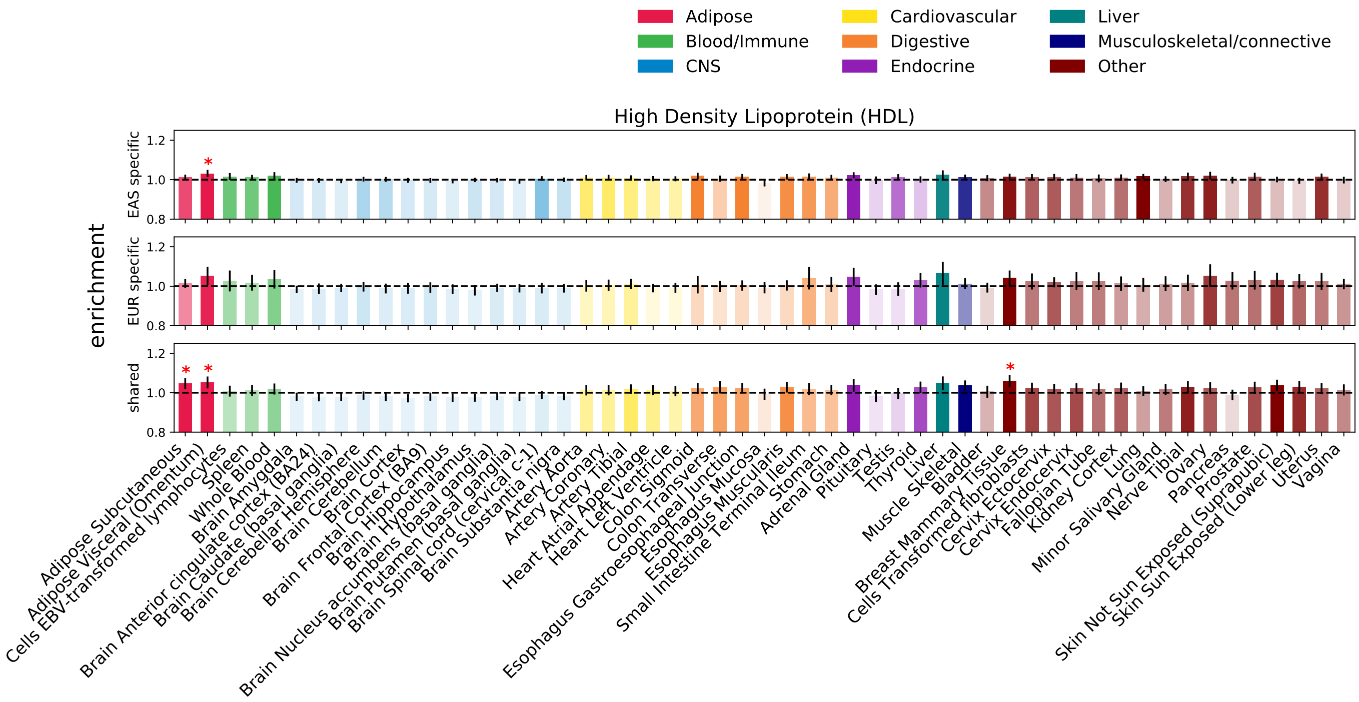

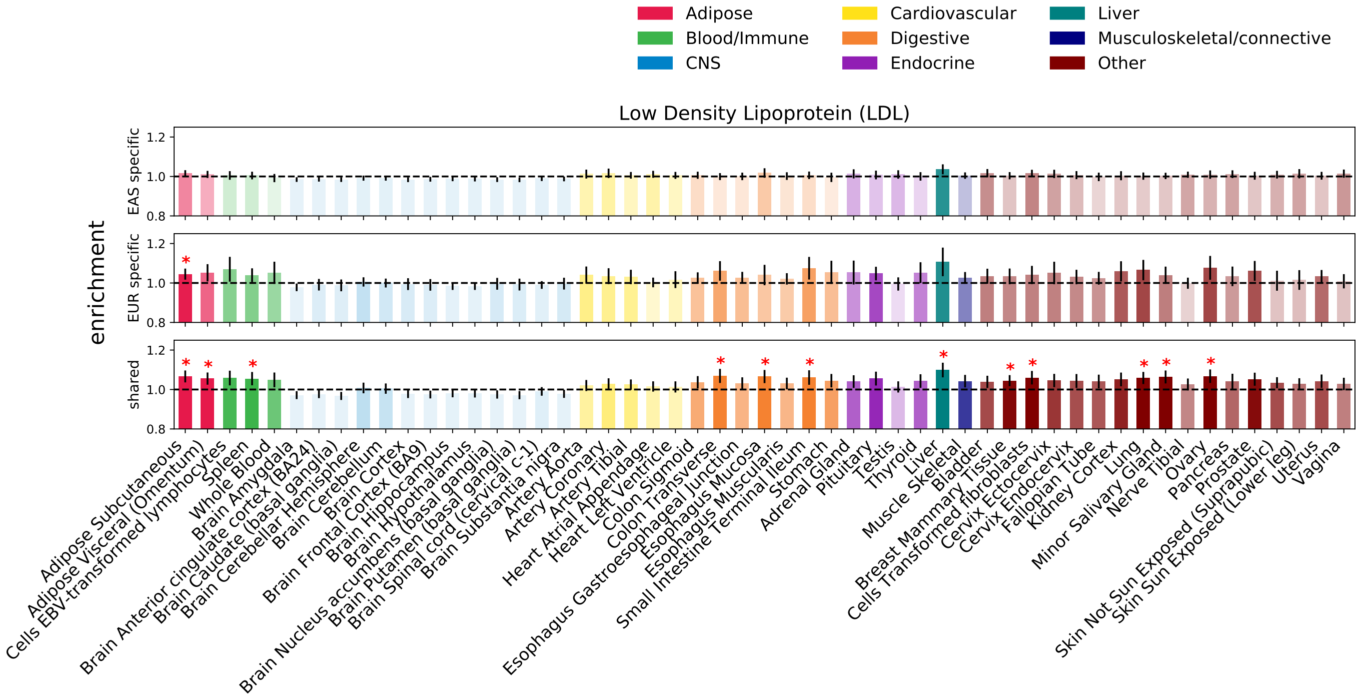

Figure S42: **Enrichment of population-specific and shared causal variants in specifically expressed genes annotation across 53 GTEx tissues.** Error bars represent 1.96 times the standard error on each side. The darker the color, the more significant an enrichment is. We mark enrichment with p-value less than $0.05/53$ with a star.

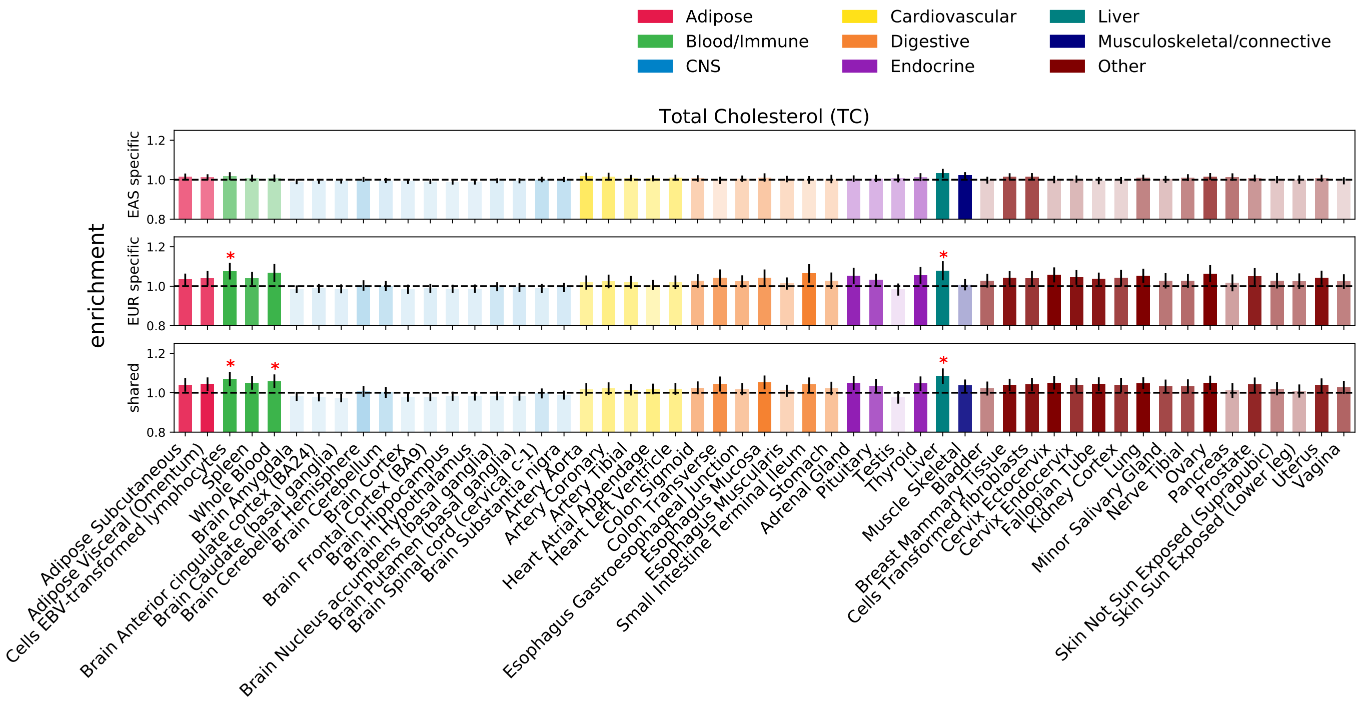

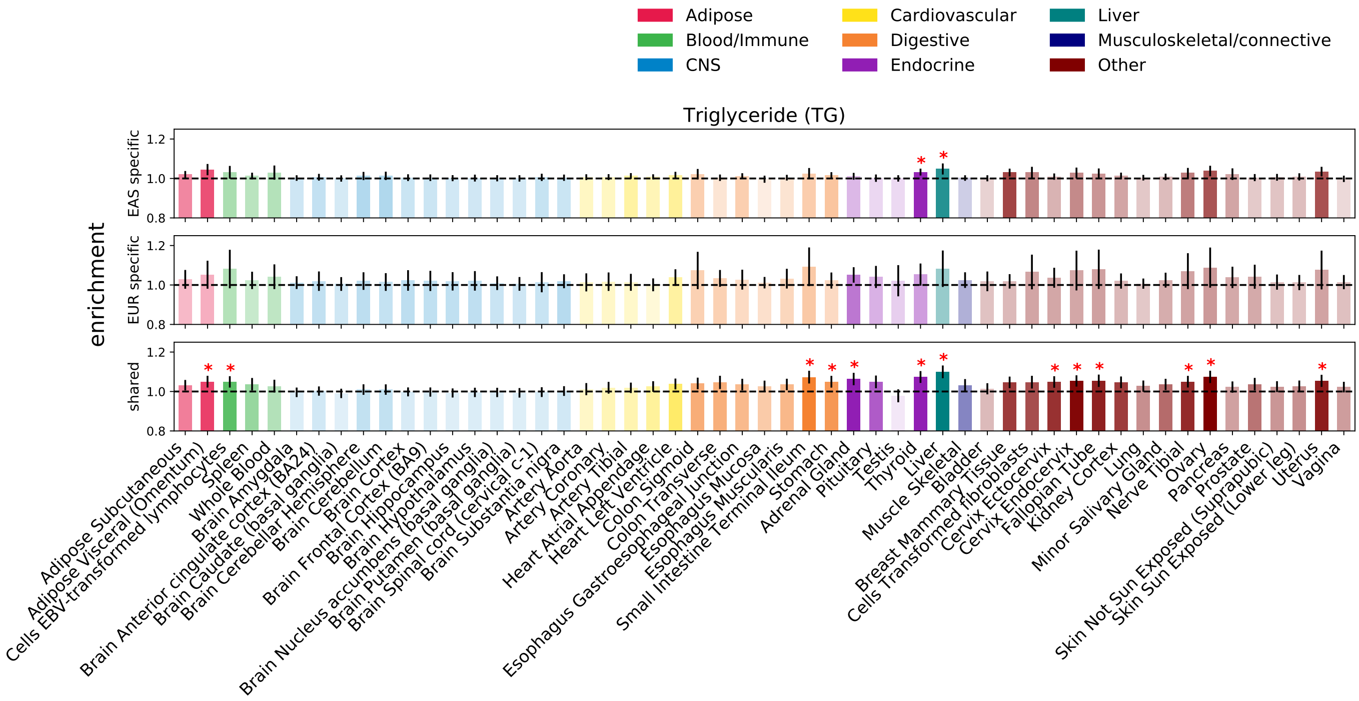

Figure S43: **Enrichment of population-specific and shared causal variants in specifically expressed genes annotation across 53 GTEx tissues.** Error bars represent 1.96 times the standard error on each side. The darker the color, the more significant an enrichment is. We mark enrichment with p-value less than $0.05/53$ with a star.

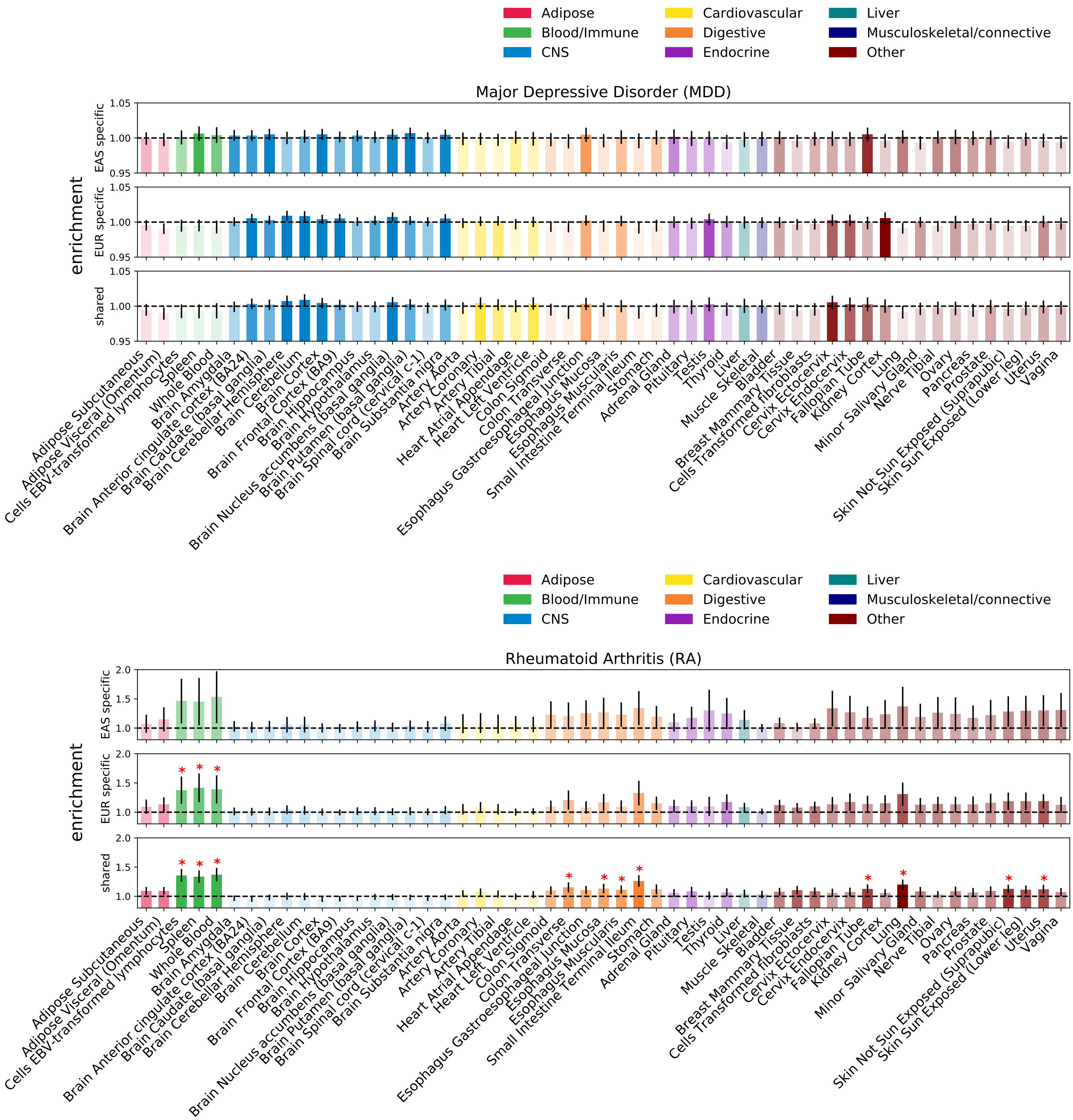

Figure S44: **Enrichment of population-specific and shared causal variants in specifically expressed genes annotation across 53 GTEx tissues.** Error bars represent 1.96 times the standard error on each side. The darker the color, the more significant an enrichment is. We mark enrichment with p-value less than $0.05/53$ with a star.

### 2 Supplemental Tables

| true_cau_status | Posterior > t | mean_l2_EAS | sem_l2_EAS | mean_l2_EUR | sem_l2_EUR | t |
|---|---|---|---|---|---|---|
| shared | shared | 6.79 | 0.08 | 6.73 | 0.08 | 0.25 |
| none | shared | 7.37 | 0.1 | 7.13 | 0.09 | 0.25 |
| EUR_only | shared | 6.87 | 0.21 | 6.44 | 0.21 | 0.25 |
| EAS_only | shared | 6.55 | 0.19 | 6.72 | 0.21 | 0.25 |
| shared | EAS_only | 6.57 | 0.23 | 6.75 | 0.25 | 0.25 |
| none | EAS_only | 6.54 | 0.22 | 6.58 | 0.25 | 0.25 |
| EAS_only | EAS_only | 6.49 | 0.21 | 6.61 | 0.24 | 0.25 |
| shared | EUR_only | 6.74 | 0.2 | 6.16 | 0.19 | 0.25 |
| none | EUR_only | 7.02 | 0.26 | 6.57 | 0.22 | 0.25 |
| EUR_only | EUR_only | 7.02 | 0.29 | 6.36 | 0.23 | 0.25 |
| EAS_only | EUR_only | 5.27 | 0.18 | 5.46 | 0.75 | 0.25 |
| shared | shared | 6.52 | 0.11 | 6.39 | 0.1 | 0.5 |
| EUR_only | shared | 6.72 | 0.34 | 6.12 | 0.33 | 0.5 |
| EAS_only | shared | 6.36 | 0.32 | 6.54 | 0.35 | 0.5 |
| none | shared | 7.2 | 0.2 | 7.09 | 0.18 | 0.5 |
| shared | EAS_only | 6.45 | 0.4 | 6.49 | 0.35 | 0.5 |
| none | EAS_only | 7.18 | 0.73 | 7.89 | 0.9 | 0.5 |
| EAS_only | EAS_only | 6.94 | 0.45 | 6.82 | 0.53 | 0.5 |
| EUR_only | EUR_only | 6.3 | 0.35 | 5.84 | 0.3 | 0.5 |
| shared | EUR_only | 7.27 | 0.39 | 6.4 | 0.31 | 0.5 |
| none | EUR_only | 8.1 | 0.91 | 7.23 | 0.72 | 0.5 |

Table S1: **Average LD scores of SNPs with posterior probability $> t$ for at least one causal configuration.** For each set of SNPs with posterior probability $> t$ (i.e. SNPs classified as shared, EAS-specific, or EUR-specific with respect to a given threshold), we stratified the SNPs by their true causal statuses and report the mean and S.E.M. of their EAS and EUR LD scores. Column 1 contains the true causal statuses; column 2 contains the causal configurations for which at least two SNPs have posterior probability $> t$.

### 3 Supplemental Note

#### 3.1 The multivariate Bernoulli (MVB) distribution

The multivariate Bernoulli (MVB) is a generalization of the Bernoulli for modeling the distribution of a binary vector of arbitrary size[2,3]. Let $\boldsymbol{B} \in \{0,1\}^p$ represent a random binary vector of size $p$ that follows an MVB distribution. The distribution of $\boldsymbol{B}$ can be described by $2^p$ probabilities, namely $\Pr(\boldsymbol{B} = 0, \cdots, 0), \cdots, \Pr(\boldsymbol{B} = 1, \cdots, 1)$, one for each of the $2^p$ possible realizations of $\boldsymbol{B}$[2,3]. Alternatively, one can adopt an index set representation of the binary vector $\boldsymbol{B}$, $\boldsymbol{A} = \{i : \boldsymbol{B}_i = 1\}$, the set of indices of 1's in $\boldsymbol{B}$, and represent the distribution of $\boldsymbol{B}$ as the ratio

$$\Pr(\boldsymbol{B}) = \Pr(\boldsymbol{A}) = \frac{\exp\left(\sum_{\boldsymbol{C} \subseteq \boldsymbol{A}} f_{\boldsymbol{C}}\right)}{\sum_{\boldsymbol{D}} \exp\left(\sum_{\boldsymbol{C} \subseteq \boldsymbol{D}} f_{\boldsymbol{C}}\right)} = \frac{\exp\left(S_{\boldsymbol{A}}\right)}{\sum_{\boldsymbol{D}} \exp\left(S_{\boldsymbol{D}}\right)}, \tag{1}$$

where $f_{\boldsymbol{C}}$ contains the natural parameters of the MVB[2,3], and $S_{\boldsymbol{A}} = \sum_{\boldsymbol{C} \subseteq \boldsymbol{A}} f_{\boldsymbol{C}}$.

We use the convention that the right-most bit in the binary vector is the first bit, and the left-most bit is the last bit. For the sake of convenience, we use binary string and index set representation of binary vectors interchangeably (e.g., both the binary string 011 and the index set $\{1, 2\}$ represent the binary vector $(0, 1, 1)$).

As a concrete example, consider a binary vector of size 2. The probabilities of each possible realization of a binary vector of size 2 under the MVB are

$$\begin{aligned}
\Pr(00) = \Pr(\phi) &= \frac{\exp(f_{00})}{\exp(f_{00}) + \exp(f_{00} + f_{01}) + \exp(f_{00} + f_{10}) + \exp(f_{00} + f_{01} + f_{10} + f_{11})} \\
\Pr(01) = \Pr(\{1\}) &= \frac{\exp(f_{00} + f_{01})}{\exp(f_{00}) + \exp(f_{00} + f_{01}) + \exp(f_{00} + f_{10}) + \exp(f_{00} + f_{01} + f_{10} + f_{11})} \\
\Pr(10) = \Pr(\{2\}) &= \frac{\exp(f_{00} + f_{10})}{\exp(f_{00}) + \exp(f_{00} + f_{01}) + \exp(f_{00} + f_{10}) + \exp(f_{00} + f_{01} + f_{10} + f_{11})} \\
\Pr(11) = \Pr(\{1,2\}) &= \frac{\exp(f_{00} + f_{01} + f_{10} + f_{11})}{\exp(f_{00}) + \exp(f_{00} + f_{01}) + \exp(f_{00} + f_{10}) + \exp(f_{00} + f_{01} + f_{10} + f_{11})}
\end{aligned} . \tag{2}$$

#### 3.2 Modeling GWAS summary statistics in two ancestral populations

##### 3.2.1 MVB prior for a SNP's causal status in two ancestral populations

We use a binary vector of size 2, $\boldsymbol{C}_i = (\boldsymbol{c}_{i1}, \boldsymbol{c}_{i2})$, to model the causal statuses of SNP $i$ in two ancestral populations. In total, there are 4 possible binary vectors of size 2: (1) if $\boldsymbol{C}_i = 00$, the SNP is causal in neither population; (2) if $\boldsymbol{C}_i = 01$, the SNP is causal in population 1 only; (3) if $\boldsymbol{C}_i = 10$, the SNP is causal in population 2 only; (4) and if $\boldsymbol{C}_i = 11$, the SNP is causal in both populations. $\boldsymbol{C}_i$ can be modeled using a multinomial distribution, Mult$(p_{00}, p_{01}, p_{10}, p_{11})$, where $p_{00}$, $p_{01}$, $p_{10}$, and $p_{11}$ represent the probability of each possible binary vector of size 2. Equivalently, one can model $\boldsymbol{C}_i$ through the MVB as,

$$\begin{aligned} \Pr(\boldsymbol{C}_i = 00) &= \frac{\exp(f_{00})}{\eta},\ \Pr(\boldsymbol{C}_i = 01) = \frac{\exp(f_{01} + f_{00})}{\eta} \\ \Pr(\boldsymbol{C}_i = 10) &= \frac{\exp(f_{10} + f_{00})}{\eta},\ \Pr(\boldsymbol{C}_i = 11) = \frac{\exp(f_{11} + f_{10} + f_{01} + f_{00})}{\eta}, \end{aligned} \tag{3}$$

where $\eta = \exp(f_{00}) + \exp(f_{01} + f_{00}) + \exp(f_{10} + f_{00}) + \exp(f_{11} + f_{10} + f_{01} + f_{00})$ is the normalization constant and $\boldsymbol{f} = (f_{00}, f_{01}, f_{10}, f_{11})$ are the parameters of the MVB (see Equation (2)).

Since the MVB distribution is invariant with respect to the parameter $f_{00}$, we enforce $f_{00} = 0$ as a convention[2]. The parameters $f_{01}$ and $f_{10}$ govern the probability of a SNP being causal in a single population, and $f_{11}$ governs the dependence of the causal statuses between two populations: $f_{11} = 0$ indicates independence and $f_{11} \neq 0$ indicates dependence[2,3]. Since the MVB parameters are real numbers (i.e. $\boldsymbol{f} \in \mathbb{R}^4$), they can be estimated using unconstrained optimization.

###### 3.2.2 Joint distribution of GWAS summary statistics in two ancestral populations

We model a phenotype in two ancestral populations using the linear models $\boldsymbol{Y}_1 = \boldsymbol{X}_1\boldsymbol{\beta}_1 + \boldsymbol{\epsilon}_1$ and $\boldsymbol{Y}_2 = \boldsymbol{X}_2\boldsymbol{\beta}_2 + \boldsymbol{\epsilon}_2$, where $\boldsymbol{Y}_1 \in \mathbb{R}^{n_1}$ and $\boldsymbol{Y}_2 \in \mathbb{R}^{n_2}$ are the phenotype measurements for $n_1$ in population 1 and $n_2$ individuals in population 2, respectively, $\boldsymbol{X}_1 \in \mathbb{R}^{n_1 \times p}$ and $\boldsymbol{X}_2 \in \mathbb{R}^{n_2 \times p}$ are column-standardized genotype matrices for $p$ SNPs, $\boldsymbol{\beta}_1 \in \mathbb{R}^p$ and $\boldsymbol{\beta}_2 \in \mathbb{R}^p$ are the standardized effect sizes of the $p$ SNPs in the two populations, and $\boldsymbol{\epsilon}_1 \in \mathbb{R}^{n_1}$ and $\boldsymbol{\epsilon} \in \mathbb{R}^{n_2}$ are environmental effects. We further assume that each row of $\boldsymbol{X}_1$ and $\boldsymbol{X}_2$ is drawn from a distribution with covariance $\boldsymbol{V}_1$ and $\boldsymbol{V}_2$, the $p \times p$ LD matrix in each population, respectively, and that for individual $n$, $\boldsymbol{\epsilon}_{1n} \sim N\left(0, \sigma_{e1}^2\right)$ and $\boldsymbol{\epsilon}_{2n} \sim N\left(0, \sigma_{e2}^2\right)$, where $\sigma_{e1}^2$ and $\sigma_{e2}^2$ represent variance of the environmental effects in population 1 and 2, respectively.

In a typical GWAS, one obtains association statistics (Z-scores) of every SNP as

$$\begin{aligned} \boldsymbol{Z}_1 &= \frac{1}{\sqrt{n_1}}\boldsymbol{X}_1^\intercal \boldsymbol{Y}_1 \\ \boldsymbol{Z}_2 &= \frac{1}{\sqrt{n_2}}\boldsymbol{X}_2^\intercal \boldsymbol{Y}_2 \end{aligned} \tag{4}$$

which have been shown to follow the multivariate normal distributions[4]

$$\begin{aligned} \boldsymbol{Z}_1|\boldsymbol{\beta}_1 &\sim N\left(\sqrt{n_1}\boldsymbol{V}_1\boldsymbol{\beta}_1, \sigma_{e1}^2\boldsymbol{V}_1\right) \\ \boldsymbol{Z}_2|\boldsymbol{\beta}_2 &\sim N\left(\sqrt{n_2}\boldsymbol{V}_1\boldsymbol{\beta}_2, \sigma_{e2}^2\boldsymbol{V}_2\right) \end{aligned} \tag{5}$$

Given the causal status vectors, $\boldsymbol{c}_1$ and $\boldsymbol{c}_2$, of every SNP in each population, one obtains the conditional distributions $\boldsymbol{Z}_1|\boldsymbol{\beta}_1, \boldsymbol{c}_1$ and $\boldsymbol{Z}_2|\boldsymbol{\beta}_2, \boldsymbol{c}_2$ as

$$\begin{aligned} \boldsymbol{Z}_1|\boldsymbol{\beta}_1, \boldsymbol{c}_1 &\sim N\left(\sqrt{n_1}\boldsymbol{V}_1(\boldsymbol{\beta}_1 \circ \boldsymbol{c}_1), \sigma_{e1}^2\boldsymbol{V}_1\right) \\ \boldsymbol{Z}_2|\boldsymbol{\beta}_2, \boldsymbol{c}_2 &\sim N\left(\sqrt{n_2}\boldsymbol{V}_2(\boldsymbol{\beta}_2 \circ \boldsymbol{c}_2), \sigma_{e2}^2\boldsymbol{V}_2\right) \end{aligned} \tag{6}$$

where $\circ$ denotes the Hadamard product[5].

Following Equation (6), one can evaluate the likelihood of $\boldsymbol{Z}_1$ and $\boldsymbol{Z}_2$ given the true causal effect size vectors $\boldsymbol{\beta}_1$ and $\boldsymbol{\beta}_2$. However, in reality the true causal effect size vectors are not given, and estimating these parameters from data will likely lead to over-fitting. Instead, we impose a normal prior on each causal SNP in $\boldsymbol{\beta}_1$ and $\boldsymbol{\beta}_2$ to obtain

$$\begin{aligned} \boldsymbol{\beta}_1|\boldsymbol{c}_1 &\sim N\left(\boldsymbol{0}, \frac{h_{g1}^2}{|\boldsymbol{c}_1|}\,\mathrm{diag}(\boldsymbol{c}_1)\right), \\ \boldsymbol{\beta}_2|\boldsymbol{c}_2 &\sim N\left(\boldsymbol{0}, \frac{h_{g2}^2}{|\boldsymbol{c}_2|}\,\mathrm{diag}(\boldsymbol{c}_2)\right), \end{aligned} \tag{7}$$

where $h_{g1}^2$, $h_{g2}^2$ are the SNP-heritability of the phenotype in the two populations and $|\boldsymbol{c}_1|$, $|\boldsymbol{c}_2|$ denote the number of 1's (i.e. the number of causal SNPs) in the binary vectors[6,7,8]. With the normal prior on $\boldsymbol{\beta}_1$ and $\boldsymbol{\beta}_2$, the conditional distributions $\boldsymbol{Z}_1|\boldsymbol{c}_1$ and $\boldsymbol{Z}_2|\boldsymbol{c}_2$ are

$$\begin{aligned} \boldsymbol{Z}_1|\boldsymbol{c}_1 &\sim N\left(\boldsymbol{0}, \boldsymbol{V}_1 + \sigma_1^2\boldsymbol{V}_1\,\mathrm{diag}(\boldsymbol{c}_1)\boldsymbol{V}_1\right), \\ \boldsymbol{Z}_2|\boldsymbol{c}_2 &\sim N\left(\boldsymbol{0}, \boldsymbol{V}_2 + \sigma_2^2\boldsymbol{V}_2\,\mathrm{diag}(\boldsymbol{c}_2)\boldsymbol{V}_2\right), \end{aligned} \tag{8}$$

where $\sigma_1^2 = \frac{n_1 h_{g1}^2}{|\boldsymbol{c}_1|}$ and $\sigma_2^2 = \frac{n_2 h_{g2}^2}{|\boldsymbol{c}_2|}$.

Incorporating the MVB prior on the causal status vectors, the joint distribution of $\boldsymbol{Z}_1$ and $\boldsymbol{Z}_2$, which is parameterized by the MVB parameters, $\boldsymbol{f} = (f_{00}, f_{01}, f_{10}, f_{11})$, is

$$\begin{aligned}\Pr(\boldsymbol{Z}_1, \boldsymbol{Z}_2; \boldsymbol{f}) &= \sum_{\boldsymbol{c}_1}\sum_{\boldsymbol{c}_2} \Pr(\boldsymbol{Z}_1, \boldsymbol{Z}_2, \boldsymbol{c}_1, \boldsymbol{c}_2; \boldsymbol{f}) = \sum_{\boldsymbol{c}_1}\sum_{\boldsymbol{c}_2} \Pr(\boldsymbol{Z}_1|\boldsymbol{c}_1)\Pr(\boldsymbol{Z}_2|\boldsymbol{c}_2)\Pr(\boldsymbol{c}_1, \boldsymbol{c}_2; \boldsymbol{f}) \\ &= \sum_{\boldsymbol{c}_1}\sum_{\boldsymbol{c}_2}\left[\begin{array}{c} N(\boldsymbol{Z}_1; \boldsymbol{0}, \boldsymbol{V}_1 + \sigma_1^2 \boldsymbol{V}_1 \operatorname{diag}(\boldsymbol{c}_1)\boldsymbol{V}_1) \times \\ N(\boldsymbol{Z}_2; \boldsymbol{0}, \boldsymbol{V}_2 + \sigma_2^2 \boldsymbol{V}_2 \operatorname{diag}(\boldsymbol{c}_2)\boldsymbol{V}_2) \times \prod_{i=1}^{p} \frac{\exp(S_{\boldsymbol{C}_i})}{\sum_{\boldsymbol{B}} \exp(S_{\boldsymbol{B}})} \end{array}\right]\end{aligned} \tag{9}$$

To model the joint distribution of GWAS summary statistics across $L$ LD-independent regions, we take the product of the probability of Z-scores across regions:

$$\begin{aligned}&\Pr(\boldsymbol{Z}_{1\{1,\cdots,L\}}, \boldsymbol{Z}_{2\{1,\cdots,L\}\}}; \boldsymbol{f}) = \prod_{l=1}^{L} \Pr(\boldsymbol{Z}_{1l}, \boldsymbol{Z}_{2l}; \boldsymbol{f}) \\ &= \prod_{l=1}^{L}\left\{\sum_{\boldsymbol{c}_{1l}}\sum_{\boldsymbol{c}_{2l}}\left[\begin{array}{c} N(\boldsymbol{Z}_{1l}; \boldsymbol{0}, \boldsymbol{V}_{1l} + \sigma_{1l}^2 \boldsymbol{V}_{1l} \operatorname{diag}(\boldsymbol{c}_{1l})\boldsymbol{V}_{1l}) \times \\ N(\boldsymbol{Z}_{2l}; \boldsymbol{0}, \boldsymbol{V}_{2l} + \sigma_{2l}^2 \boldsymbol{V}_{2l} \operatorname{diag}(\boldsymbol{c}_{2l})\boldsymbol{V}_{2l}) \times \prod_{i=1}^{p_l} \frac{\exp(S_{\boldsymbol{C}_{li}})}{\sum_{\boldsymbol{B}} \exp(S_{\boldsymbol{B}})} \end{array}\right]\right\}.\end{aligned} \tag{10}$$

#### 3.3 Model fitting using Expectation Maximization

##### 3.3.1 Expectation step

We use expectation-maximization (EM) to estimate the model parameters $\boldsymbol{f}$. First, we derive the complete log-likelihood of the data

$$\begin{aligned}&\ell\left(\boldsymbol{f}|\boldsymbol{Z}_{1\{1,\cdots,L\}}, \boldsymbol{Z}_{2\{1,\cdots,L\}}, \boldsymbol{c}_{1\{1,\cdots,L\}}, \boldsymbol{c}_{2\{1,\cdots,L\}}\right) \\ &= \log\left\{\prod_{l=1}^{L}\left[\begin{array}{c} N(\boldsymbol{Z}_{1l}; \boldsymbol{0}, \boldsymbol{V}_{1l} + \sigma_{1l}^2 \boldsymbol{V}_{1l} \operatorname{diag}(\boldsymbol{c}_{1l})\boldsymbol{V}_{1l}) \times \\ N(\boldsymbol{Z}_{2l}; \boldsymbol{0}, \boldsymbol{V}_{2l} + \sigma_{2l}^2 \boldsymbol{V}_{2l} \operatorname{diag}(\boldsymbol{c}_{2l})\boldsymbol{V}_{2l}) \times \prod_{i=1}^{p_l} \frac{\exp(S_{\boldsymbol{C}_{li}})}{\sum_{\boldsymbol{B}} \exp(S_{\boldsymbol{B}})} \end{array}\right]\right\} \\ &= \sum_{l=1}^{L}\left[\log N(\boldsymbol{Z}_{1l}; \boldsymbol{0}, \boldsymbol{V}_{1l} + \sigma_{1l}^2 \boldsymbol{V}_{1l} \operatorname{diag}(\boldsymbol{c}_{1l})\boldsymbol{V}_{1l}) + \log N(\boldsymbol{Z}_{2l}; \boldsymbol{0}, \boldsymbol{V}_{2l} + \sigma_{2l}^2 \boldsymbol{V}_{2l} \operatorname{diag}(\boldsymbol{c}_{2l})\boldsymbol{V}_{2l})\right] \\ &\quad + \sum_{l=1}^{L}\sum_{i=1}^{p_l} S_{C_{li}} - \log\left(\sum_{\boldsymbol{B}} \exp(S_{\boldsymbol{B}})\right)\sum_{l=1}^{L} p_l.\end{aligned} \tag{11}$$

In the expectation step of the EM algorithm, one finds the expectation of the log-likelihood with respect to the causal status vectors $\boldsymbol{c}_{1\{1,\cdots,L\}}$, $\boldsymbol{c}_{2\{1,\cdots,L\}}$, conditioned on the current estimate of the model parameters $\boldsymbol{f}^{(t)}$,

$$\begin{aligned}
Q\left(\boldsymbol{f}|\boldsymbol{f}^{(t)}\right) &= \mathrm{E}\left[\ell\left(\boldsymbol{f}|\boldsymbol{Z}_{1\{1,\cdots,L\}},\boldsymbol{Z}_{2\{1,\cdots,L\}},\boldsymbol{c}_{1\{1,\cdots,L\}},\boldsymbol{c}_{2\{1,\cdots,L\}}\right)\right] \\
&= \sum_{l=1}^{L}\sum_{\boldsymbol{c}_{1l},\boldsymbol{c}_{2l}} \Pr\left(\boldsymbol{c}_{1l},\boldsymbol{c}_{2l}|\boldsymbol{f}^{(t)},\boldsymbol{Z}_{1l},\boldsymbol{Z}_{2l}\right)\left[\begin{array}{c}\log N(\boldsymbol{Z}_{1l};\boldsymbol{0},\boldsymbol{V}_{1l}+\sigma_{1l}^2\boldsymbol{V}_{1l}\operatorname{diag}(\boldsymbol{c}_{1l})\boldsymbol{V}_{1l}) \\ +\log N(\boldsymbol{Z}_{2l};\boldsymbol{0},\boldsymbol{V}_{2l}+\sigma_{2l}^2\boldsymbol{V}_{2l}\operatorname{diag}(\boldsymbol{c}_{2l})\boldsymbol{V}_{2l})\end{array}\right] \\
&\quad + \sum_{l=1}^{L}\sum_{\boldsymbol{c}_{1l},\boldsymbol{c}_{2l}} \Pr\left(\boldsymbol{c}_{1l},\boldsymbol{c}_{2l}|\boldsymbol{f}^{(t)},\boldsymbol{Z}_{1l},\boldsymbol{Z}_{2l}\right)\left(\sum_{i=1}^{p_l} S_{\boldsymbol{C}_{li}}\right) - \log\left(\sum_{\boldsymbol{B}}\exp(S_{\boldsymbol{B}})\right)\sum_{l=1}^{L} p_l,
\end{aligned} \tag{12}$$

where $\Pr\left(\boldsymbol{c}_{1l},\boldsymbol{c}_{2l}|\boldsymbol{f}^{(t)},\boldsymbol{Z}_{1l},\boldsymbol{Z}_{2l}\right)$ is

$$\Pr\left(\boldsymbol{c}_{1l},\boldsymbol{c}_{2l}|\boldsymbol{f}^{(t)},\boldsymbol{Z}_{1l},\boldsymbol{Z}_{2l}\right) = \frac{\Pr\left(\boldsymbol{c}_{1l},\boldsymbol{c}_{2l},\boldsymbol{Z}_{1l},\boldsymbol{Z}_{2l}|\boldsymbol{f}^{(t)}\right)}{\sum_{\boldsymbol{b}_{1l},\boldsymbol{b}_{2l}}\Pr\left(\boldsymbol{b}_{1l},\boldsymbol{b}_{2l},\boldsymbol{Z}_{1l},\boldsymbol{Z}_{2l}|\boldsymbol{f}^{(t)}\right)}. \tag{13}$$

##### 3.3.2 Maximization step

The goal of the maximization step is to find

$$\boldsymbol{f}^{(t+1)} = \operatorname{argmax}_{\boldsymbol{f}} Q\left(\boldsymbol{f}|\boldsymbol{f}^{(t)}\right) = \operatorname{argmax}_{\boldsymbol{f}} g(\boldsymbol{f}) \tag{14}$$

where

$$g(\boldsymbol{f}) = \sum_{l=1}^{L}\sum_{\boldsymbol{c}_{1l},\boldsymbol{c}_{2l}} \Pr\left(\boldsymbol{c}_{1l},\boldsymbol{c}_{2l}|\boldsymbol{f}^{(t)},\boldsymbol{Z}_{1l},\boldsymbol{Z}_{2l}\right)\left(\sum_{i=1}^{p_l} S_{\boldsymbol{C}_{li}}\right) - \log\left(\sum_{\boldsymbol{B}}\exp(S_{\boldsymbol{B}})\right)\sum_{l=1}^{L} p_l, \tag{15}$$

removing the irrelevant constant in $Q(\boldsymbol{f}|\boldsymbol{f}^{(t)})$.

Evaluating $g(\boldsymbol{f})$ involves a summation over all possible causal status vectors, which has time complexity on the order of $O(2^{2p_l})$ and is intractable. Instead, we recognize that

$$\begin{aligned}
g(\boldsymbol{f}) &= \sum_{l=1}^{L}\sum_{\boldsymbol{c}_{1l},\boldsymbol{c}_{2l}} \mathrm{E}\left[\sum_{i=1}^{p_l} S_{\boldsymbol{C}_{li}}\right] - \log\left(\sum_{\boldsymbol{B}}\exp(S_{\boldsymbol{B}})\right)\sum_{l=1}^{L} p_l \\
&\approx h(\boldsymbol{f}) = \sum_{l=1}^{L}\left[\frac{1}{J}\sum_{j=1}^{J}\left(\sum_{i=1}^{p_l} S_{\boldsymbol{C}_{li}^{(j)}}\right)\right] - \log\left(\sum_{\boldsymbol{B}}\exp(S_{\boldsymbol{B}})\right)\sum_{l=1}^{L} p_l,
\end{aligned} \tag{16}$$

where $\boldsymbol{C}_{li}^{(j)} = \left(\boldsymbol{c}_{1i}^{(j)},\boldsymbol{c}_{2i}^{(j)}\right)$ represents the causal status of the $i$-th SNP at locus $l$ in the two populations, from the causal status vectors, $\boldsymbol{c}_1^{(j)}$, $\boldsymbol{c}_2^{(j)}$, sampled from the posterior distribution

$\Pr(\boldsymbol{c}_{1l}, \boldsymbol{c}_{2l} | \boldsymbol{Z}_{1l}, \boldsymbol{Z}_{2l}, \boldsymbol{f}^*)$. We use Gibbs sampling to efficiently sample causal status vectors from the posterior (see Section 3.4).

It can be shown that the following parameter updates maximizes $h(\boldsymbol{f})$,

$$\begin{aligned}
\boldsymbol{f}_{00}^{(t+1)} &= 0, \\
\boldsymbol{f}_{01}^{(t+1)} &= \log \bar{q}_{01} - \log \bar{q}_{00}, \\
\boldsymbol{f}_{10}^{(t+1)} &= \log \bar{q}_{10} - \log \bar{q}_{00}, \\
\boldsymbol{f}_{11}^{(t+1)} &= \log \bar{q}_{11} - \log \bar{q}_{01} - \log \bar{q}_{10} + \log \bar{q}_{00},
\end{aligned} \tag{17}$$

where $\bar{q}_{00}$, $\bar{q}_{01}$, $\bar{q}_{10}$, and $\bar{q}_{11}$ represent the average count of 01, 10, and 11 causal status at a single SNP in two ancestral populations across MCMC samples from the Gibbs sampler (see Section 3.4).

#### 3.4 Sampling causal status vectors from posterior distribution

We use Gibbs sampling to sample $\boldsymbol{C} = (\boldsymbol{c}_1, \boldsymbol{c}_2)$ from the posterior distribution,

$$\boldsymbol{C} \sim \Pr(\boldsymbol{C} | \boldsymbol{f}, \boldsymbol{Z}_1, \boldsymbol{Z}_2) \propto \Pr(\boldsymbol{Z}_1, \boldsymbol{Z}_2, \boldsymbol{C} | \boldsymbol{f}). \tag{18}$$

For notational simplicity, we drop the index $l$ representing different loci. To advance the Markov chain from step $j$ to step $j+1$ in Gibbs sampling, at step $j$ we select SNP $k$ and evaluate the probability of the four possible cross-population causal configurations at that SNP,

$$\begin{aligned}
&\Pr\left(\boldsymbol{Z}_1, \boldsymbol{Z}_2, \boldsymbol{C}_k = 00, \boldsymbol{C}_{\neg j}^{(j)} | \boldsymbol{f}\right) \ \Pr\left(\boldsymbol{Z}_1, \boldsymbol{Z}_2, \boldsymbol{C}_k = 01, \boldsymbol{C}_{\neg j}^{(j)} | \boldsymbol{f}\right) \\
&\Pr\left(\boldsymbol{Z}_1, \boldsymbol{Z}_2, \boldsymbol{C}_k = 10, \boldsymbol{C}_{\neg j}^{(j)} | \boldsymbol{f}\right) \ \Pr\left(\boldsymbol{Z}_1, \boldsymbol{Z}_2, \boldsymbol{C}_k = 11, \boldsymbol{C}_{\neg j}^{(j)} | \boldsymbol{f}\right),
\end{aligned} \tag{19}$$

where $\boldsymbol{C}_{\neg j}^{(j)}$ denotes the rest of the causal configurations, excluding that of SNP $k$ in the $j$-th step. We then sample $\boldsymbol{C}^{(j+1)}$ based on the following probability

$$\Pr\left(\boldsymbol{C}^{(t+1)} = \left(\boldsymbol{C}_k = \boldsymbol{b}', \boldsymbol{C}_{\neg j}^{(j)}\right)\right) = \frac{\Pr\left(\boldsymbol{Z}_1, \boldsymbol{Z}_2, \boldsymbol{C}_k = \boldsymbol{b}', \boldsymbol{C}_{\neg j}^{(j)} | \boldsymbol{f}\right)}{\sum_{\boldsymbol{b}} \Pr\left(\boldsymbol{Z}_1, \boldsymbol{Z}_2, \boldsymbol{C}_k = \boldsymbol{b}, \boldsymbol{C}_{\neg j}^{(j)} | \boldsymbol{f}\right)}. \tag{20}$$

To evaluate $\Pr(\boldsymbol{Z}_1, \boldsymbol{Z}_2, \boldsymbol{c}_1, \boldsymbol{c}_2 | \boldsymbol{f}) = \Pr(\boldsymbol{Z}_1 | \boldsymbol{c}_1) \Pr(\boldsymbol{Z}_2 | \boldsymbol{c}_2) \Pr(\boldsymbol{c}_1, \boldsymbol{c}_2 | \boldsymbol{f})$, we note that previous work has shown that

$$\Pr(\boldsymbol{Z}_1|\boldsymbol{c}_1) = N\left(\boldsymbol{Z}_1|\boldsymbol{0}, \boldsymbol{V}_1 + \sigma_1^2\boldsymbol{V}_1^2\right) \propto \frac{N\left(\boldsymbol{Z}_{1\boldsymbol{c}_1}|\boldsymbol{0}, \boldsymbol{V}_{1\boldsymbol{c}_1} + \sigma_1^2\boldsymbol{V}_{1\boldsymbol{c}_1}^2\right)}{N\left(\boldsymbol{Z}_{1\boldsymbol{c}_1}|\boldsymbol{0}, \boldsymbol{V}_{1\boldsymbol{c}_1}\right)}, \tag{21}$$

where $BF_1 = \frac{N\left(\boldsymbol{Z}_{1\boldsymbol{c}_1}|\boldsymbol{0}, \boldsymbol{V}_{1\boldsymbol{c}_1} + \sigma_1^2\boldsymbol{V}_{1\boldsymbol{c}_1}^2\right)}{N\left(\boldsymbol{Z}_{1\boldsymbol{c}_1}|\boldsymbol{0}, \boldsymbol{V}_{1\boldsymbol{c}_1}\right)}$ is the Bayes factor at only the causal SNPs, reducing the time complexity of evaluating the probability from $p^3$ to $p_{\text{causal}}^3$. Let $\boldsymbol{V}_{1\boldsymbol{c}_1} = \sum_{i=1}^{p_{\text{causal}}} w_i \boldsymbol{u}_i \boldsymbol{u}_i^\intercal$ be the eigenvalue decomposition of $\boldsymbol{V}_{1\boldsymbol{c}_1}$, where $w_i$ and $\boldsymbol{u}_i$ are the eigenvalues and eigenvectors of $\boldsymbol{V}_{1\boldsymbol{c}_1}$. We further note that $BF_1$ can be expressed as

$$\begin{aligned} BF_1 &= \frac{\det(\boldsymbol{V}_{1\boldsymbol{c}_1} + \sigma_1^2\boldsymbol{V}_{1\boldsymbol{c}_1}^2)^{-\frac{1}{2}} \exp\left[-\frac{1}{2}\boldsymbol{Z}_{1\boldsymbol{c}_1}^\intercal(\boldsymbol{V}_{1\boldsymbol{c}_1} + \sigma_1^2\boldsymbol{V}_{1\boldsymbol{c}_1}^2)^{-1}\boldsymbol{Z}_{1\boldsymbol{c}_1}\right]}{\det(\boldsymbol{V}_{1\boldsymbol{c}_1})^{-\frac{1}{2}} \exp\left(-\frac{1}{2}\boldsymbol{Z}_{1\boldsymbol{c}_1}^\intercal\boldsymbol{V}_{1\boldsymbol{c}_1}^{-1}\boldsymbol{Z}_{1\boldsymbol{c}_1}\right)} \\ &\propto \left(\prod_{i=1}^{p_{\text{causal}}} \frac{1}{1+\sigma_1^2 w_i}\right)^{\frac{1}{2}} \exp\left[\frac{1}{2}\sum_{i=1}^{p_{\text{causal}}} \frac{\sigma_1^2}{1+\sigma_1^2 w_i}\left(\boldsymbol{Z}_{1\boldsymbol{c}_1}^\intercal \boldsymbol{u}_i\right)^2\right], \end{aligned} \tag{22}$$

avoiding numerical instability introduced by small eigenvalues. The Bayes factor for $\boldsymbol{Z}_{2\boldsymbol{c}_2}$ can be obtained using the same approach.

#### 3.5 Posterior probability of each SNP to be ancestry-specific or shared

We use posterior probability of each SNP to be ancestry-specific or shared to quantify evidence of ancestry-specific or shared genetic architecture at single-SNP resolution. Specifically, we evaluate

$$\Pr(\boldsymbol{C}_i = \boldsymbol{b}|\boldsymbol{Z}_1, \boldsymbol{Z}_2, \boldsymbol{f}^*) \tag{23}$$

for $\boldsymbol{b} \in \{01, 10, 11\}$ at each SNP $i$, where $\boldsymbol{f}^*$ denotes the estimated MVB parameter. We show below that the per-SNP posterior probability in Equation (23) can be evaluated using the Gibbs sampling procedure outlined in Section 3.4. First, we note that

$$\begin{aligned} \Pr(\boldsymbol{C}_i = \boldsymbol{b}|\boldsymbol{Z}_1, \boldsymbol{Z}_2, \boldsymbol{f}^*) &= \sum_{\boldsymbol{C}_{\neg i}} \Pr(\boldsymbol{C}_i = \boldsymbol{b}, \boldsymbol{C}_{\neg i}|\boldsymbol{Z}_1, \boldsymbol{Z}_2\boldsymbol{f}^*) \\ &= \sum_{\boldsymbol{C}_{\neg i}} \Pr(\boldsymbol{C}_i = \boldsymbol{b}|\boldsymbol{C}_{\neg i}, \boldsymbol{Z}_1, \boldsymbol{Z}_2\boldsymbol{f}^*) \Pr(\boldsymbol{C}_{\neg i}|\boldsymbol{Z}_1, \boldsymbol{Z}_2\boldsymbol{f}^*) \\ &= \mathrm{E}\left[\Pr(\boldsymbol{C}_i = \boldsymbol{b}|\boldsymbol{C}_{\neg i}, \boldsymbol{Z}_1, \boldsymbol{Z}_2\boldsymbol{f}^*)\right] = \mathrm{E}\left[\mathrm{E}[\mathbb{1}_{\{\boldsymbol{C}_i=\boldsymbol{b}\}}|\boldsymbol{C}_{\neg i}, \boldsymbol{Z}_1, \boldsymbol{Z}_2\boldsymbol{f}^*]\right] \\ &= \mathrm{E}[\mathbb{1}_{\{\boldsymbol{C}_i=\boldsymbol{b}\}}|\boldsymbol{Z}_1, \boldsymbol{Z}_2\boldsymbol{f}^*] \approx \frac{\sum_{j=1}^J \mathbb{1}_{\{\boldsymbol{C}_i^{(j)}=\boldsymbol{b}\}}}{J}, \end{aligned} \tag{24}$$

where $C^{(j)}$ is the $j$-th causal status vectors (out of a total of $J$ samples) sampled from the posterior distribution $\Pr(C|Z_1, Z_2, f^*)$ (see Section 3.4). To ensure stable estimates of the posterior probability, we run the Gibbs sampling procedure 20 times and report the average posterior probability.

##### 3.6 Defining approximately independent LD blocks in two populations

We adapted LDetect[9] to define blocks of SNPs that are approximately independent in both East Asian and European populations. Briefly, LDetect is a method to define approximately independent blocks of SNPs in a single population that estimates a regularized LD matrix for the population and identifies block-diagonal structures[9]. To define blocks of SNPs that are approximately independent in two populations, we first compute regularized LD matrices for both populations ($V_{\text{EAS}}$ and $V_{\text{EUR}}$) following the LDetect procedure[9]. We then construct a new matrix ($V_{\text{trans}}$) by taking, for each pair of SNPs, the larger of the East Asian and European LD estimates:

$$V_{\text{trasn},ij} = \begin{cases} V_{\text{EAS},ij} & \text{if} |V_{\text{EAS},ij}| > |V_{\text{EUR},ij}| \\ V_{\text{EUR},ij} & \text{if} |V_{\text{EUR},ij}| > |V_{\text{EAS},ij}| \end{cases}. \quad (25)$$

The matrix $V_{\text{trans}}$ is block diagonal due to shared recombination hot spots in both populations. We then applied the LDetect procedure[9] to define block structures in $V_{\text{trans}}$. Using this approach, we identified 1,368 LD blocks (2Mb wide on average) that are approximately independent in both East Asian and European populations.

##### 3.7 Simulation framework

We used real genotypes on chromosome 22 from CONVERGE[10] and the UK Biobank[11] to simulate GWAS summary statistics for East Asian and European populations. We used genotype data from the 1000 Genomes Project (Phase 3)[12] as the reference panel. Since SNPs in perfect LD have identical Z-scores, we performed minimal LD pruning (at an $R^2$ threshold of 0.95) on the reference LD matrices using PLINK 1.9[13]. We also removed strand-ambiguous SNPs and SNPs with minor allele frequency (MAF) less than 1% in either population, resulting in a total of 8,599 SNPs on chromosome 22.

We simulated phenotypes based from the linear models $Y_1 = X_1\beta_1 + \epsilon_1, Y_2 = X_2\beta_2 + \epsilon_2,$

where the effects at causal SNPs in each population, $\boldsymbol{\beta}_{1\boldsymbol{c}_1}$ and $\boldsymbol{\beta}_{2\boldsymbol{c}_1}$, are drawn from

$$\boldsymbol{\beta}_{1\boldsymbol{c}_1} \sim N\left(\mathbf{0}, \frac{h_{g1}^2}{|\boldsymbol{c}_1|}\boldsymbol{I}\right), \boldsymbol{\beta}_{2\boldsymbol{c}_2} \sim N\left(\mathbf{0}, \frac{h_{g2}^2}{|\boldsymbol{c}_2|}\boldsymbol{I}\right), \quad (26)$$

and effects at non-causal SNPs are set to 0. Here, $\boldsymbol{c}_1$ and $\boldsymbol{c}_2$ are the index sets of causal SNPs in each population. We simulated the environmental effect for the $n$-th individual in each population as $\boldsymbol{\epsilon}_{1n} \sim N(0, 1-h_{g1}^2)$ and $\boldsymbol{\epsilon}_{2n} \sim N(0, 1-h_{g2}^2)$. Finally, we compute Z-scores for all SNPs following Equation (4).
